# Neural sensing of surface mechanics modulates proprioceptive activity and locomotion in *Caenorhabditis elegans*

**DOI:** 10.64898/2026.03.05.709070

**Authors:** Aleksandra Pidde, Montserrat Porta-de-la-Riva, Costanza Agazzi, Carmen Martínez-Fernández, Al-ice Lorrach, Ashutosh Bijalwan, Neus Sanfeliu-Cerdán, Alba Calatayud-Sanchez, Eric Torralba-Sales, Ravi Das, José J. Muñoz, Michael Krieg

## Abstract

Locomotion — whether walking, running, or crawling — depends on the precise coordination of forces between the body and its surroundings. Two critical factors in this process are the force that resists the relative motion between two bodies, and mechanosensation, the body’s ability to sense and respond to mechanical forces. Together, they allow organisms to move efficiently, adapt to varying environments, and maintain balance. Here we show that the ‘gentle touch’ receptor neurons (TRNs) in the *Caenorhabditis elegans* body wall are sensitive to dynamic surface traction. Using a combination of calcium recordings and traction force microscopy in freely moving animals, microfluidics, and whole connectome computer simulations, we show that MEC-4 DEG/ENaC ion channel activity depends on the crawling velocity and friction force. Mutations disrupting MEC-4 activity and body wall mechanoreceptor function produce lethargic worms with impaired proprioceptive regulation, suggesting functional coupling between surface mechanoreceptors and proprioceptors. Our data reveal a new role for classical touch receptors in locomotion and critically define the mechanical modality sensed by skin mechanosensors.

## Main Text

Mechanosensation allows us to explore the immediate physical environment (*1*). In addition to conscious touch, mechanosensation plays a key role in communication, bonding, and social comfort (*2,3*), and provides essential mechanical information about the surface when we walk (*4*). This feedback mechanism is crucial for adaptive locomotion and proprioception, enabling individuals to respond to uneven terrain and varying support surfaces (*5, 6*). Although they play a critical role in the somatosensory system, the cellular and mechanical mechanisms of how animals process plantar feedback and which mechanical stimuli activate these receptors remains understudied and therefore poorly understood.

*Caenorhabditis (C.) elegans* senses mechanical forces delivered to their body wall with a set of different mechanoreceptors (*1, 7*). Depending on the stimulus magnitude, these mechanoreceptors have been coined harsh and gentle touch receptors. The six neurons responsible for sensing gentle mechanical stimuli are called PVM, PLM(L/R), ALM(L/R) and AVM and tile the body in anterior and posterior receptive fields (Fig. 1a; (*7*)). However, and despite being mechanosensitive (*8*), laser ablation of PVM resulted in a negligible effect on the behavioral response to touch (*9*), questioning its role in mechanosensation. What other stimuli, beyond touch, might TRNs have evolved to sense? Importantly, mutations in genes that cause loss of sensory function cause a striking lethargic phenotype (*9–11*). Likewise, loss of the pore-forming subunit of the mechanoelectrical transduction channel *mec-4* (*12, 13*), a member of the Degenerin/Epithelial sodium/acid sensing ion channel (DEG/ENaC/ASiC) family, leads to changes in the generation of locomotor patterns, suggesting that TRNs induce a behavioral response to a changing physical environment (*14, 15*). Lastly, it was also shown that TRNs, through MEC-4 ion channels sense mechanical shear stresses (*16*), exhibiting higher activity when stimuli are delivered at a higher rate (*8, 17*), similar to human mechanoreceptors involved in touch (*18*). Together, this may hint towards a role of these neurons in locomotion, through sensing dynamic mechanical properties such as substrate traction forces.

**Fig. 1.**
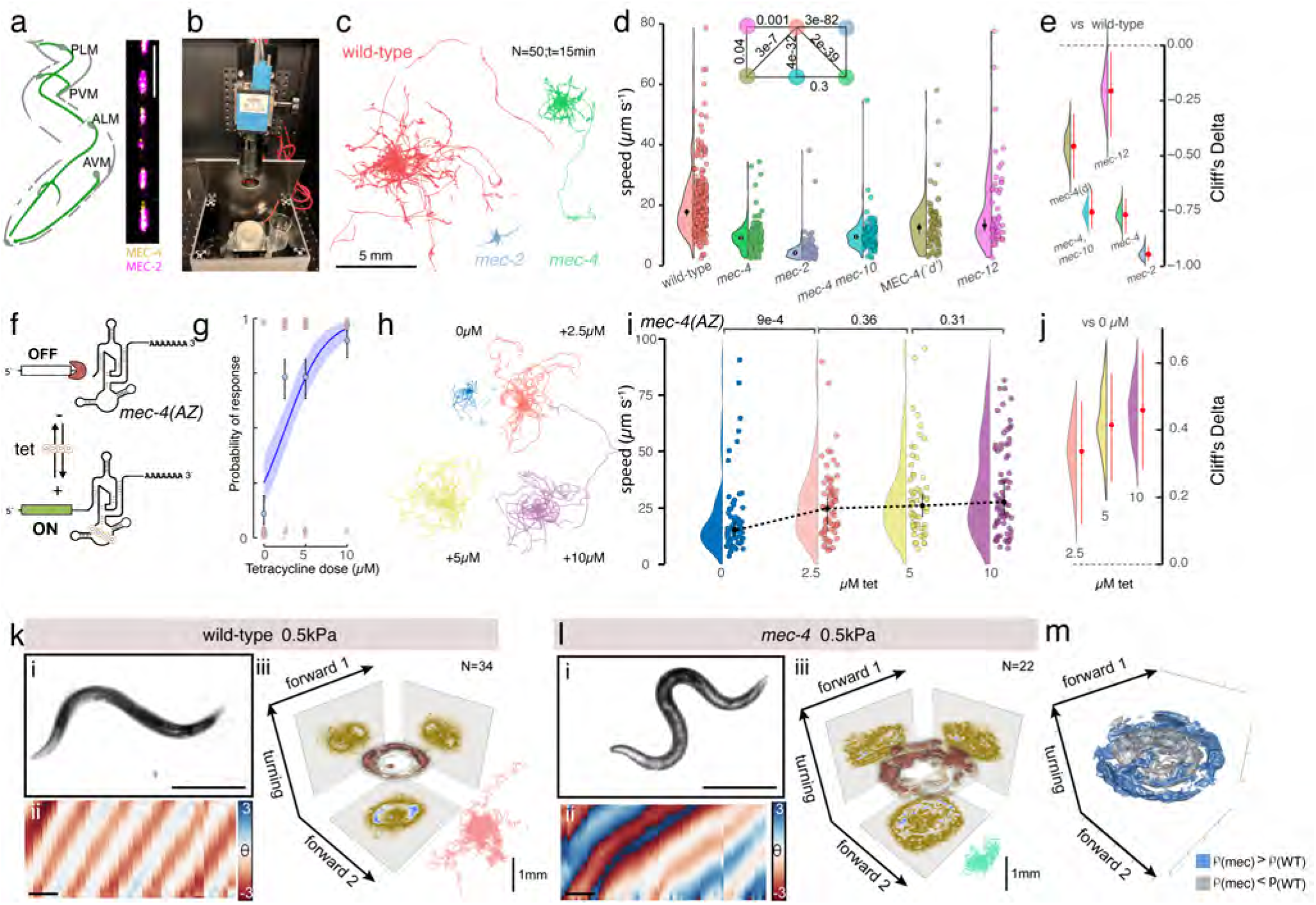
Body-wall mechanoreceptor function promotes locomotion in *C. elegans*. **a,** Schematic of the worm with the location of the body wall mechanoreceptors shown in green (left) and a representative micrograph of MEC-2/MEC-4 colocalization in the ALM neurite (right). Scale bar = 5µm. **b,** Photograph of the light and vibration-isolated small animal behavior tracking platform. **c,** Trajectories of 15 min locomotion behavior for wild-type animals, *mec-2* and *mec-4* mutants. *N* = 50 animals per condition on NGM plates supplemented with food. **d,** Distribution of the crawling speed from wild-type and various *mec* mutant genes visualized as a violin plot. *MEC-4(‘d’)* indicates the degenerin mutation leading to loss of TRNs in adult animals (*e1611*, A713V, (*56*)). Black solid circles show medians, range indicates 95% confidence interval. For exact number of measurements and replicates, consult Supplementary Table 3. Inset: Two-sided Kolmogorov-Smirnov test p-values comparing each pair of distributions (numbers inside the matrix). Colors correspond to genotype groups consistently across violins, estimation plots, and the KS matrix. **e,** Estimation plots of bootstrapped Cliff’s Delta (red points indicate median, red bars show 95% bias-corrected confidence intervals) for each comparison relative to the reference genotype. A dashed horizontal line at zero indicates no effect. **f,** Schematic of the aptazyme (az), and the acute rescue experiment using tetracycline to block it. **g,** Representative results of a touch assay in animals expressing *mec-4(az)* at increasing tetracycline concentrations. The probability of responding in the 10-touch assay increased with dose (log-odds = 0.416 ± 0.066 per µM, z = 6.34, *p* = 2.3 10^−10^), with notable animal-to-animal variability (random intercept SD = 0.89). Points show individual trial outcomes (0 = no response, 1 = response). The solid line represents the predicted probability from a mixed-effects binary logistic regression with tetracycline dose as a fixed effect and animals as random intercepts, and the shaded area indicates the 95% confidence interval. Blue circle and range indicate mean SD. **h,** Trajectories acquired for 15 min from 30 animals under each condition analyzed for *mec-4(az)* and the indicated concentrations of tetracycline. **i,** Violin plot of crawling speed in *mec-4(az)* animals under the stated tetracycline concentrations. Black circles show medians, range indicates 95% confidence interval. *P*-values comparing each pair of distributions calculated from a two-sided KS-test. See Supplementary Table 3 for measurement details. **j,** Estimation plots of bootstrapped Cliff’s Delta (red points indicate median, red bars show 95% bias-corrected confidence intervals) for each comparison relative to the reference genotype. A dashed horizontal line at zero indicates no effect. **k-m,** Animal body posture during locomotion on elastic substrates. Representative photograph (i, scale bar = 200um) and curvature kymograph (ii, scale bar = 2s) of a wild-type (k), and *mec-4* mutant (l), animal crawling a on 0.5 kPa elastic hydrogel substrate and (iii) their joint probability distribution (equivalent to a discrete 3D histogram) in the Eigenworm subspace (see Methods). Trajectories in the lower right indicate track length for N=34 (22) animals.**m**, 3D plot of the statistically significant differences comparing the joint probability distribution of wild-type (k), and *mec-4* mutant (l) animals crawling on 0.5 kPa substrates. Silver voxels indicate higher density for wild-type animal, blue voxels indicate higher density for *mec-4* mutants with *p <*0.01.

Here we synthesize this large body of previous work with new evidence that gentle body wall mechanoreceptors sense friction with the surrounding environment and use MEC-4 as a velocity-dependent force sensor. In particular, we define a role for PVM in sensing substrate compliance, a neuron that specifically activates in response to navigation in a soft environment. Using traction force microscopy, we map the forces that are exerted by the animal on the substrate and show that direct stimulation of PVM is reflected in different proprioceptive circuits, in a *mec-4* dependent manner. Based on these data and whole connectome simulations, we propose that the gentle body wall mechanoreceptors provide essential information about substrate mechanics, crosstalk with proprioceptive neurons, akin to mammalian plantar mechanoreceptors.

## Results

### Loss of body-wall mechanoreception leads to lethargic animals

In addition to their failure to respond to external touch to the body wall, *mec-4* mutants are known to be lethargic (*9*). To corroborate and extend these observations, we first recorded 15 min long videos in an isolated chamber, protecting animals from external light, vibrations, and variations of temperature and humidity (Fig. 1b, see Methods) to understand whether the loss of mechanosensitivity to touch is correlated to a loss of locomotor activity. We tracked individual animals lacking TRNs (MEC-4(’d’)) and various mutant alleles with loss-of-function *mec-4*, *mec-10*, *mec-2* and *mec-12*. Although some of these genes are expressed in other neurons than TRNs (*19*), they all functionally intersect in TRNs and share a common role in mechanotransduction. Their loss of function causes a strong, almost complete absence of touch sensitivity and touch-induced currents (*13*). We found that hermaphrodite mutants of all alleles are significantly less mobile than wild-type animals of similar age on food (Fig. 1c-e, Supplementary Video 1), affecting both forward and backward locomotion (Extended Data Fig. 1a). As previously described in (*9*), this difference is strongly reduced off food, whereas *mec-4* and *mec-2* mutant males retain their mobile activity on food (Extended Data Fig. 1b, and ref. (*9*)). Moreover, locomotor defects can be induced either by ectopic expression of a miniSOG transgene (*20*) to ablate individual TRNs upon blue-light illumination (Extended Data Fig. 1c) or by blocking synaptic transmission through TRN-specific expression of tetanus toxin (TeTx, Extended Data Fig. 1d), suggesting that vesicular neurotransmission underlies TRN-mediated locomotion phenotypes. Because animals expressing miniSOG in the dark displayed already light-independent defects, we sought to determine whether the effect arises from the chronic absence of mechanoreception or can be induced acutely. To do so, we expressed the anion channelrhodopsin ACR1 in TRNs using the UAS/Gal4 system (*21*). Upon green-light stimulation, ACR1 activation transiently hyperpolarized TRNs and reliably produced a lethargus-like state, demonstrating that acute reversible silencing of mechanoreceptor neurons is sufficient to elicit this behavior (Extended Data Fig. 1e,f). As described previously (*22–24*), the Mec mutant phenotypes are not caused by defects in muscle function or general paralysis because all mutants retain normal locomotion on standard NGM agar, with no gross differences from wild-type when stimulated with cues other than gentle touch. Thus, our data suggest that mechanosensitivity in TRNs is linked to spontaneous locomotor behavior.

We then asked if MEC-4-induced neuronal activity patterns can induce changes in synaptic plasticity and whether their absence causes lethargic phenotypes. Previously, a repetitive optogenetic stimulus administered to channelrhodopsin 2-expressing (ChR) motor neurons in the midbody was shown to induce the bending motion of the head to a new frequency (*25, 26*). TRNs do not display rhythmic activity patterns but occasional stochastic calcium spikes in their sensory dendrite in immobilized wild-type animals (Extended Data Fig. 2a). Thus, we sought to emulate a stochastic activity pattern in animals expressing the blue light gated ion channel ChR2 in TRNs (*24*). When applying a random optogenetic stimulus to *mec-4* mutant animals for 48 h (Extended Data Fig. 2b), we were unable to overcome the lethargic phenotype associated with it (Extended Data Fig. 2c). Together, our findings indicate that the entrainment protocols used here are insufficient to explain plasticity-related phenotypes in the absence of MEC-4. However, we cannot rule out the possibility that MEC-4-dependent plasticity requires longer or developmentally timed stimulation protocols.

To provide direct evidence that MEC-4 function enhances agile locomotion, we engineered an aptamer ribozyme (*27*) in the 3’UTR of the endogenous *mec-4* locus that leads to steady and efficient degradation of its mRNA (Fig. 1f). Indeed, the introduction of the ribozyme led to the disruption of MEC-4 protein, as visualized by the absence of fluorescent signal of a mScarlet::MEC-4 fusion (see Methods and below). Consequently, these animals do not respond to external touch (Fig. 1g) and are lethargic (Fig. 1h-j). Importantly, the ribozyme’s function to cleave itself can be disabled upon the addition of low concentrations of tetracycline and the addition of these concentrations of tetracycline itself had little effect on locomotion velocity in wild-type animals (Supplementary Fig. 1a), supporting specificity of the aptazyme-mediated rescue. Using RT-PCR, we verified that the addition of tetracycline rescued *mec-4* RNA expression (Supplementary Fig. 1b-d), and assayed these animals in a classical touch assay. Indeed, animals not only responded to external touch after tetracycline addition (Fig. 1g), but their increase in mechanosensitivity was accompanied by an increase of MEC-4 protein expression (Supplementary Fig. 1d), coinciding with robust rescue of the lethargic locomotion phenotype (Fig. 1h-j). Thus, this acute rescue effect from its endogenous locus strongly supports the causal role of MEC-4 mechanosensitivity in locomotion. Consequently, we hypothesize that TRNs are mechanosensors on the body wall involved in detecting the mechanical load of the environment to induce animal locomotion - similar to plantar mechanoreceptors in mammals.

### External mechanoreception adjusts proprioceptive feedback

To understand whether MEC-4, through expression in body wall mechanosensors, plays a role in substrate sensing, we studied the crawling behavior of wild-type and *mec-4* mutant young adult animals onto polyacrylamide (PAA) hydrogels with different nominal stiffness, ranging from 0.5kPa to 100kPa (Fig. 1k-m, Extended Data Fig. 3a, see Methods). Because the animals did not move on PAA gels with abundant food, and the bacterial lawn interfered with high-resolution traction force microscopy, we carried out these experiments in the absence of a food source (OP50 bacteria). We did not find a significant difference in the locomotion gait of wild-type control animals crawling on the different stiffness substrates, suggesting that wild-type animals are able to adapt to these environments (Extended Data Fig. 3a, c). However, *mec-4* mutant animals crawled with significantly higher body curvature on softer gels (Fig. 1k-m, compare Supplementary Video 2 and 3), while on substrates with a higher stiffness, animals lacking MEC-4 crawled with significantly lower curvature (Extended Data Fig. 3e) and are slightly slower compared to wild-type controls (Extended Data Fig. 3b). This suggests that MEC-4, using TRNs as a neuronal substrate, senses the mechanical properties of the environment and thus may provide input to proprioceptive neurons to control during locomotion.

Intriguingly, the structural connectome reveals numerous synaptic contacts between PLM/PVM and PDE and DVA (Fig. 2a; (*28*)), suggesting potential functional coupling between TRNs and proprioceptors. To obtain a systems-preview on the functional connections between the touch circuit and the proprioceptive feedback circuit, we adapted a computational model incorporating the full synaptic and gap junctions connectome (Fig. 2b). This model has previously been used to understand the influence of individual neurons on forward locomotion, using motor neuron activity to predict muscle contraction and thus behavior (*29, 30*). Here, we specifically asked whether loss of TRN function alters motor neuron population dynamics when projected into a low-dimensional neural manifold, analogous to the eigenworm representation of body posture (*31, 32*). Indeed, computationally ‘injected’ constant current to PLM is sufficient to generate stable periodic activity in motor neurons, indicating an ability to generate sustained rhythmic dynamics as emergent property of the neural network (Fig. 2c).

**Fig. 2.**
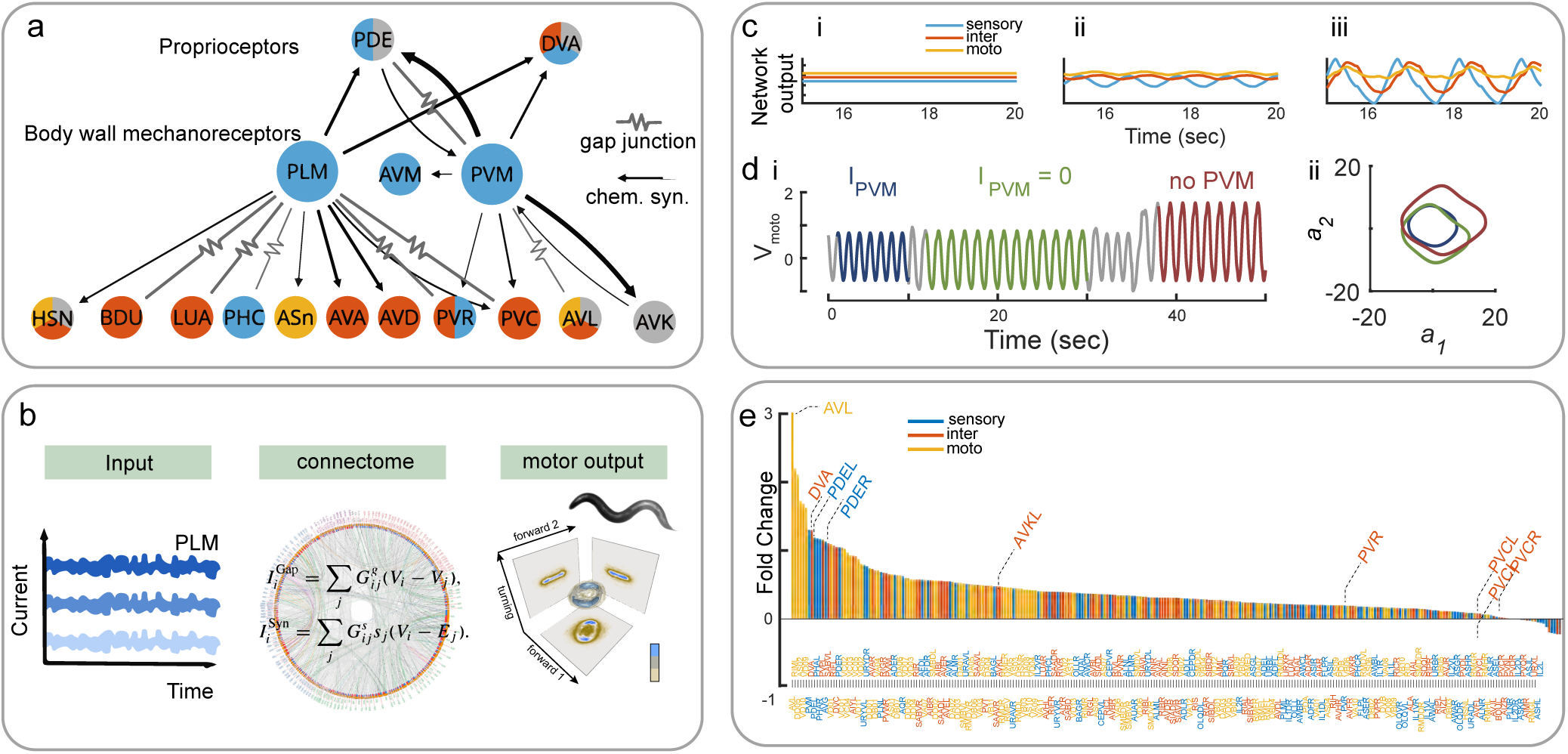
Whole neural network dynamics *in silico* suggest crosstalk between mechanosensory circuit and proprioceptive neurons. **a,** Wiring diagram of PVM and PLM. Color of the nodes indicates type of neuron (blue = sensory; orange = inter; yellow = motor; grey = modulatory neurons). Thickness of the edges indicates absolute number of electrical or chemical synapses between the connected neurons. Adapted from (*28*) and nemanode.org. **b,** Computational pipeline to estimate the functional connectomics after constant current input (low, medium, high) into PLM. The output of the connectome is quantified as a single value decomposition of the combined motor neuron activity. **c,** Average neuronal network activity in each neuron class after applying constant stimulation current with different amplitudes to PLML/R. i) low, ii) intermediate and iii) high current input. Different colors indicate sensory (blue), inter (orange) and motor neurons (yellow). **d,** i) Average motor neuron voltage oscillations of the network receiving both PLM driving current plus a ‘mechanosensitive’ PVM current (1% of the PLM driving current, dark blue), after acute removal of PVM current, resembling the *mec-4* mutation (olive green), and after PVM ablation (see Methods 3.1, dark red). ii) Projection of the combined motor neuron output into singular vectors for all three conditions. Note, the pronounced increase in limit cycle after removal of the current from PVM and PVM ablation, representing the increase in the body curvature. **e,** Fold-change in the amplitude of oscillations in voltage activity for all neurons following the removal of PVM input (*I*_PVM_ = 0) (as in **d** i, olive green). Proprioceptors DVA and PDER/L are among the most severely affected.

Next, we asked how the network behaves when the activity of ALM, AVM, and PVM is silenced — mimicking the loss of *mec-4*. Removing the input from the anterior touch neurons (ALM/AVM) collapses the motor neuron oscillations, driving the system into a fixed point in the 2D subspace spanned by singular vectors (Supplementary Fig. 2). Also, when all input connections into PLM are silenced, we observe a drastic reduction of the motor neuron population dynamics. This is consistent with the lethargic phenotype observed for the mutations affecting TRN function (Fig. 1c-e). The simulation output was markedly different when we removed the current from PVM: the amplitude of the oscillations in many neurons was increased (Fig. 2d i), leading to a larger limit cycle in the low dimensional phase space (Fig. 2d ii). This suggests a surprising role for PVM in the regulation of body curvature.

In order to investigate the influence of the PVM neuron on the motor circuit in our simulation, we quantified the change in activity upon removal of the mechanosensitive current (emulated as 1% of the PLM driving current) from the PVM. Unexpectedly, removal of the PVM input current led to a global increase in neuronal activity. Although most neurons exhibited elevated activity levels, the largest changes were observed in the proprioceptive neurons DVA and PDE (Fig. 2e). Together, these results point toward a functional role of posterior body wall mechanoreceptors in regulating the activity of DVA and PDE proprioceptors through network-level interactions rather than purely excitatory drive. This outcome is counterintuitive given the presumed modulatory role of PVM and suggests that, despite the structural connectome indicating direct coupling, the network may operate in a regime where PVM input normally exerts a net suppressive or stabilizing influence on downstream circuits.

### Touch receptors functionally couple to mechanical proprioceptors

We tested this prediction directly and investigated PDE and DVA activity after stimulating TRNs mechanically within a microfluidic pressure trap (Fig. 3a, b and ref. (*8*)). To do so, we applied a high-frequency ‘buzz’ to the cuticle within the receptive field of PVM (*33*) and recorded DVA and PDE activity using a GCaMP6s calcium reporter (*32*) (Supplementary Video 4 and 5). We used a panneuronal calcium reporter in conjunction with the Neuropal landmark transgenes (*34*) to identify and visualize PDE activity (inset in Fig. 3a panel ii), together with PVM, PVD and SDQ on the left side of the worm body (or just PVD and PDE on the right side). Intriguingly, we could also clearly distinguish mechanoreceptor activity in PVD and PVM, but not SDQ (Figs. 3a panel ii, Supplementary Video 4), while PDE peak currents lagged behind that of other sensory neurons with a temporal delay of ≈ 200 ms. PDE responded consistently and strongly to mechanical stimuli (Fig. 3c, d) in a *mec-4* dependent manner (Fig. 3c, e). To understand if PDE receives postsynaptic input from posterior touch receptor neurons, we repeated these experiments in a transgenic animal expressing TeTx in TRNs, a highly potent inhibitor of synaptic transmission. Consistent with the hypothesis that TRNs provide synaptic input to PDE, we observed a marked reduction in mechanically evoked PDE calcium responses upon TeTx-mediated silencing of synaptic transmission in TRNs (Fig. 3f, Extended Data Fig. 4a, b). In contrast, PVM remained mechanically responsive (Extended Data Fig. 4c), indicating that the defect is selective for downstream neurons rather than a general impairment of mechanotransduction. The residual PDE activity may therefore arise from intrinsic mechanosensitivity of PDE itself or from synaptic or extrasynaptic inputs originating from neurons other than TRNs. Together, our data shows that PDE is postsynaptic and functionally connected to posterior TRNs. DVA also showed a multifaceted, context dependent response to mechanical input, which required functional MEC-4 and synaptic function (Fig. 3g-j). In many cases the DVA neuron responded with a decrease in activity that recovered during the interstimulus interval (Fig. 3g, h and Supplementary Video 5). Mutating *mec-4* and silencing TRN synaptic output with TeTx dampened DVA’s deactivation, yet paradoxically seemed to increased both the number and magnitude of secondary activation responses following mechanical stimulation (Fig. 3i, j). This effect of TeTx expression on PDE and DVA was different to the avoidance behavior to touch, as animals expressing TeTx in TRNs remained largely sensitive to mechanical stimuli (TeTx: 7.9±1.5; ctrl: 8.8±1.2; mean±SD, p=0.11; two-sided t-test). This indicates that increased TRN output causes a transient activation of PDE and a suppression of DVA activity. Because the loss of DVA activity is related to increases in body curvature (*32*) our results suggest that body wall mechanosensation finetunes proprioceptive gain within the circuit. Together, these data show that TRNs are functionally connected to proprioceptive neurons, and therefore lack of input from TRNs to proprioceptors may lead to altered locomotion behavior.

**Fig. 3.**
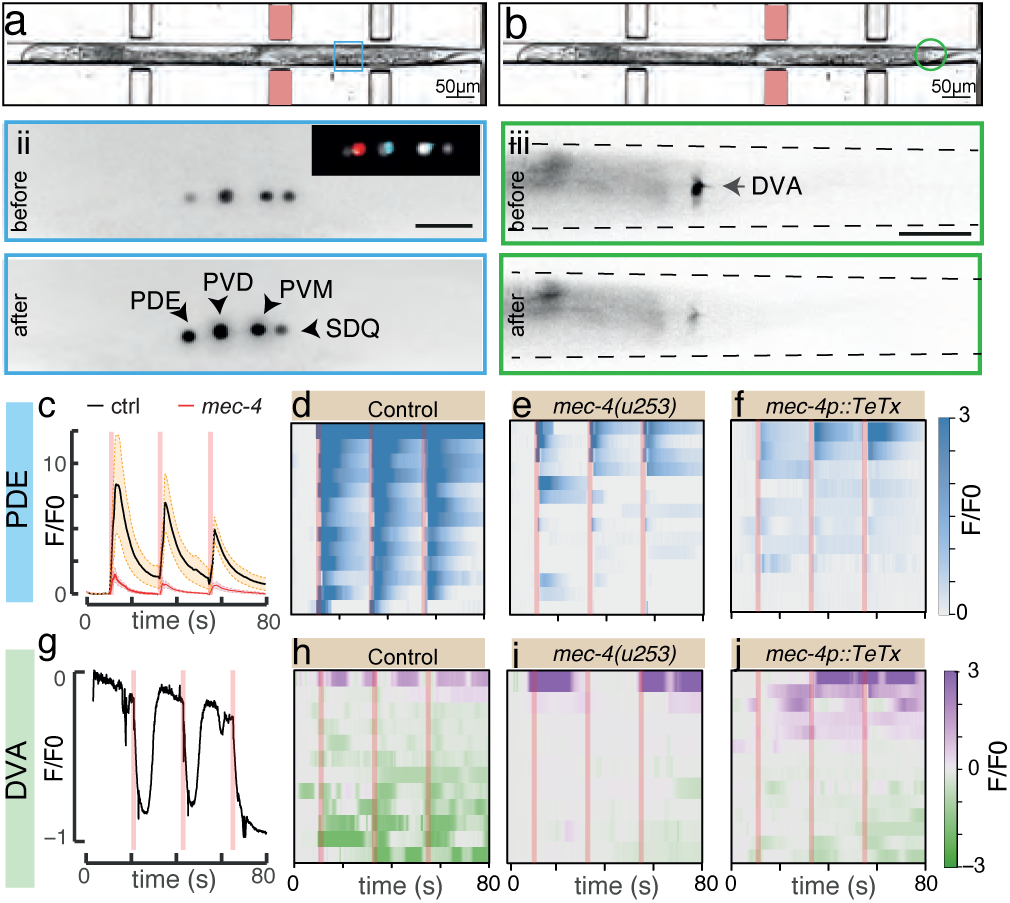
Stimulation of bodywall mechanosensors activate proprioceptive circuits. **a, b** (i) Representative image of a single animal acquired at 10x magnification in the microfluidic device, highlighting the actuator and the location of (a) PDE (blue square) and (b) DVA (green circle). The actuator used for each neuron is highlighted by a red rectangle. (ii) Fluorescent micrograph with (a) PDE and (b) DVA before and after stimulation at the posterior body part. Scale bar = 10µm. The inset in (ii) shows a snapshot of the mNeptune and mtag-BFP fluorescent tags of the Neuropal transgene taken on the same animal to unequivocally identify the four posterior midbody neurons on the left side. Dashed lines in b indicate the walls of the trapping channel. **bc** Average calcium activity of (i) PDE in control (orange) and *mec-4* (pink) mutant animals. Vertical lines indicate onset and duration of the pressure pulse. **d-f** Stacked kymographs of the Ca^2+^ activity in (d) wild-type animals (N=13), (e) *mec-4* mutant (N=14) and (f) animals expressing tetanus toxin (TeTx) in touch receptor neurons (*mec-4p(TeTx)*, N=10). **g** Representative recording of GCaMP6s signal in DVA after mechanical stimulation of the posterior body wall at times indicated by the red vertical bars. **h-j,** Stacked kymographs of the Ca^2+^ recording in DVA for (h) wild-type (N=27), (i) *mec-4* mutants (N=9) and (j) animals expressing TeTx in TRNs (N=14). Red vertical bars indicate time and duration of the mechanical stimulus.

### Touch receptor neurons sense substrate compliance

Our connectome-level simulations (Fig. 2) paired with experimental data (Figs. 1 and 3) suggest that the body wall mechanoreceptors directly sense the mechanical properties of the environment during locomotion. If so, we expect differences in neuronal activity in different mechanical environments. To test this notion, we used animals expressing GCaMP6s in TRNs of wild-type and *mec-4* mutant animals, and recorded calcium fluctuations in all anterior and posterior TRNs in freely moving animals on different PAA gels with an automated closed-loop image acquisition pipeline (see Methods 2.4). In general, we observed larger and more frequent calcium transients in moving wild-type compared to *mec-4* on all gels in all body wall mechanoreceptor neurons (Fig. 4a-c and Extended Data Fig. 5, Supplementary Video 6 and 7). The transients appeared larger on the softer gels, especially for the posterior neurons PLM and PVM (Fig. 4d and Extended Data Fig. 5). We occasionally observed increases in calcium activity that occurred secondary to self-inflicted contacts (e.g., nose-body contact), but also seemed uncorrelated with any behavioral syllable. Apart of infrequent spontaneous calcium activity confined to the neurites of TRNs, we did not observe calcium transients in completely immobilized animals (Supplementary Fig. 3a), we hypothesized that activity is related to the velocity of the animals. Indeed, time series of calcium activity and the instantaneous velocity co-varied (Fig. 4c, e), but only on softer substrates. To determine the causal and temporal relationships between calcium activity in PVM, PLM and ALM, and the animal’s velocity on different substrates, we computed the cross-covariance between these signals (see Methods) in both wild-type and *mec-4* mutant animals. We observed two dominant peaks in the covariance of the calcium signals with the tangential velocity component, indicating that the TRNs exhibit a nested and context-dependent activity (Fig. 4e).

**Fig. 4.**
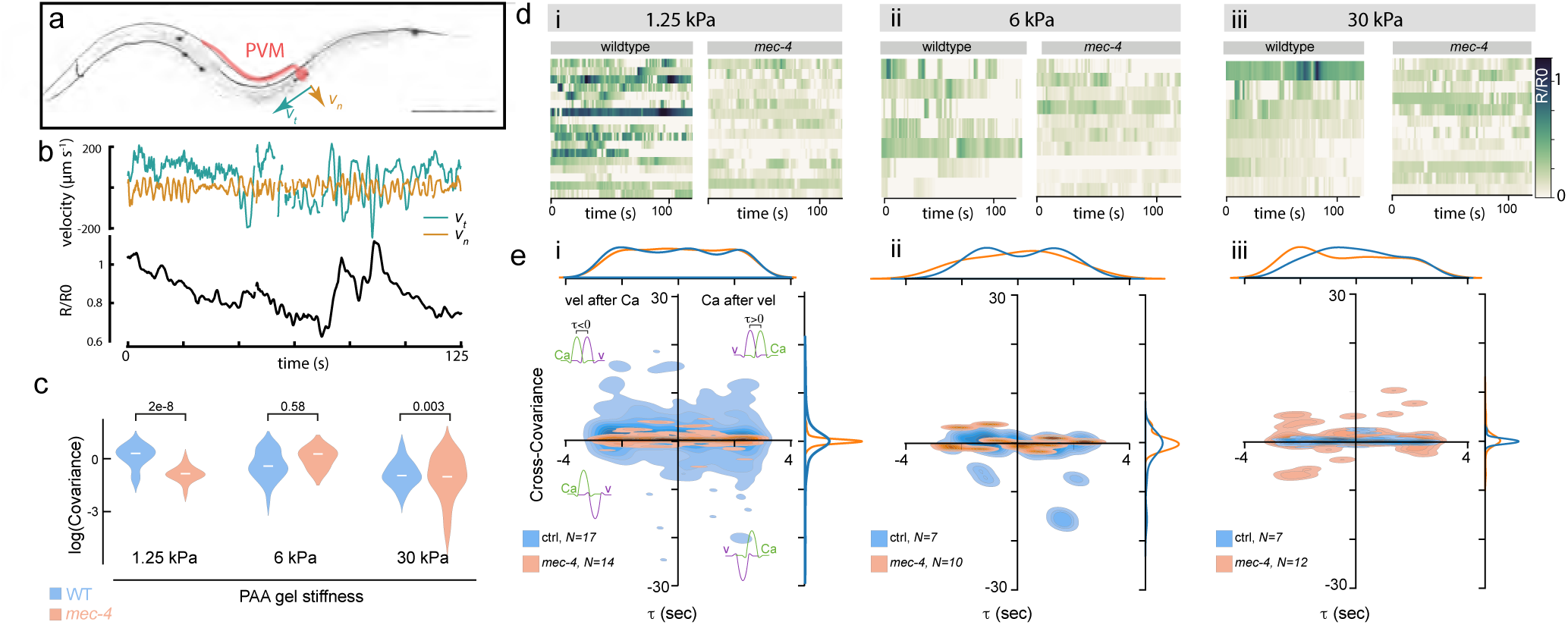
PVM senses substrate compliance. **a,** Schematic of the worms and the location of PVM neuron. Normal and tangential velocity vectors (*v_n_* & *v_t_*, respectively) at the location of the PVM neuron are indicated as arrows. Scale bar = 100µm. **b,** Representative instantaneous tangential (*v_t_*) and normal (*v_n_*) components of the velocity on substrates of soft stiffness (upper) and recording of normalized, ratiometric calcium activity (R/R0, see Methods) in PVM (lower). **c,** Covariance of PVM calcium activity and forward velocity on all substrates for control and *mec-4* mutant animals (see also Extended Data Figure 5 for comparison with ALM and PLM). *p*-values derived from a two-sided Wilcoxon rank-sum test. **d,** Kymograph of the ratiometric calcium activity extracted from PVM expressing GCaMP6s and tagRFP-T in freely moving animals (ctrl vs *mec-4*) on different substrates. i) 1.25kPa; ii) 6kPa; iii) 30kPa. **e,** Cross-covariance between PVM calcium activity and forward crawling velocity on i) 1.25kPa; ii) 6kPa; iii) 30kPa substrates of ctrl and *mec-4* mutant animals suggesting plausible causal relation between neural activity and velocity. Negative timelags indicate PVM calcium transients that precede bouts in velocity (e.g. due to self-inflicted body touch), whereas a positive timelag indicates that calcium transients follow bouts in velocity. Inset inside each quadrant indicates the temporal relation between the velocity and the calcium signals. See also Supplementary Video 6. *N* =number of animals analyzed. Top and right axes indicate the projected densities, aka histograms, to indicate the 1D-distribution of the measurements.

We first observed a pronounced neuronal activity pattern preceding bursts in locomotor velocity (negative lags in the covariance analysis), indicating that TRN activation is associated with subsequent increases in speed due to self-inflicted touches. This is consistent with their established role in the gentle-touch escape response. A similar relationship was observed for PVM, despite ongoing debate regarding its contribution to forward escape behavior (*35, 36*) (see Supplementary Discussion, Fig. 4e and Supplementary Fig. S4).

In addition, the covariance analysis revealed a significant peak at positive lags - most prominently for PVM, but not for ALM or PLM - indicating neuronal activity following increases in velocity (Fig. 4e; Extended Data Fig. 5; Methods). Notably, this relationship was absent when neuronal activity was compared to the normal (perpendicular) components of velocity, highlighting the directional specificity of locomotion-induced mechanical stimuli (Extended Data Fig. 6), but also on stiffer substrates.

Together, these findings suggest that PVM senses compliance of the physical environment and encodes the animal’s velocity relative to its substrate. Strikingly, these covariance signals were completely absent in *mec-4* mutant animals, suggesting that MEC-4 acts as a velocity-dependent sensor of substrate mechanics (Fig. 4e). Thus, our data show that body wall mechanoreceptors, especially PVMs, exhibit calcium signals that co-vary with tangential velocity on soft substrates, consistent with velocity-modulated mechanosensitivity, leading to a nested, context-dependent neuronal activity.

A key open question was whether the gentle body wall mechanoreceptors (TRNs) are activated primarily by shear stresses generated during crawling against a substrate, or whether viscous drag on the body wall in a fluid is sufficient to elicit activation. To disentangle these possibilities, we immersed individual animals in biologically inert fluids of defined viscosity (Halocarbon oil, 190 mPa s). These animals maintained rhythmic crawling movements, albeit with increased body curvature (Extended Data Fig. 7a), allowing us to compare neuronal responses in the presence of drag forces but in the absence of substrate friction.

We imaged calcium activity in anterior and posterior TRNs during locomotion in oil and compared this to activity on agar substrates. While occasional transients were detected, especially in PLM, the overall TRN activity was strongly reduced and covaried significantly less with locomotor kinematics (Extended Data Fig. 7b–d) compared to crawling on soft PAA gels (Fig. 4).

But how much drag force does the animal experience moving through the viscous medium? Using resistive force theory for a slender cylinder of length *L* = 1mm and radius *a* ≈ 40 *µ*m, moving laterally at velocity v=50µm/s in a fluid of viscosity *η*=0.19Pa·s, we obtain a transverse drag per unit length of 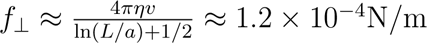.

Over a TRN-relevant length scale of Δ*x*=50*µ*m this corresponds to a local load of only *F_viscous_* = *f*_⊥_Δ*x* ≈ 6nN. This value is at least an order of magnitude below the ≈100-200nN indentation forces reported to reliably activate TRNs with a glass probe (*13*) supporting the conclusion that viscous drag is insufficient to robustly engage TRNs.

However, during crawling on agar, frictional forces can be estimated as *F_friction_* = *µN*, where *µ* is the effective coefficient of friction and N is the normal load. Traction force studies report effective frictional forces in the range of 10-1000µN per animal (*37, 38*). On the scale of a 50 µm TRN segment, this corresponds to hundreds of nN of local load - well above the activation threshold for MEC-4 - dependent mechanotransduction.

Together, both the experimental recordings and the force estimates indicate that viscous drag alone is not sufficient to strongly activate TRNs. Instead, TRNs are robustly engaged by shear stresses that arise from frictional coupling to a soft, viscoelastic substrate during crawling.

### Disruption of MEC-4 in PVM elicits a lethargic locomotion phenotype

Up to here, our results support a model in which PVM detects substrate-dependent mechanical cues during crawling. We next asked if the PVM neuron has a specific role and modulates locomotion through expression of MEC-4 ion channels. To address this question, we generated a transgenic strain in which a split Cre recombinase (*39*) was co-expressed under the control of two promoters that intersect uniquely with *mec-4* expression in PVM neurons (Fig. 5a, b). This construct was introduced into a genetic back-ground in which the endogenous *mec-4* locus was flanked by loxP sites (Fig. 5b). After assessing the specific recombination of the two loxP sites specifically in PVM (*32, 40*) (Fig. 5c) we confirmed the loss of MEC-4 expression in PVM using a functional split-mScarlet–tagged MEC-4 allele (*41*), which exhibits wild-type punctate localization (Fig. 5d) and touch sensitivity (responding to 8.85±1.23 out of ten touches, mean±SEM, N=20 animals), comparable to control animals with either split-CRE or floxed *mec-4* locus alone (Fig. 5e). Finally, we asked whether selective disruption of *mec-4* in PVM induces a lethargic locomotion phenotype. Compared to animals that expressed the split CRE alone or the loxP-flanked *mec-4* allele alone, double-transgenic animals lacking *mec-4* specifically in PVM exhibited significantly reduced locomotor speed and increased lethargy (Fig. 5f). Together with our calcium-imaging data, these results indicate that MEC-4 expression in PVM has a role in substrate mechanosensation important for animal locomotion.

**Fig. 5.**
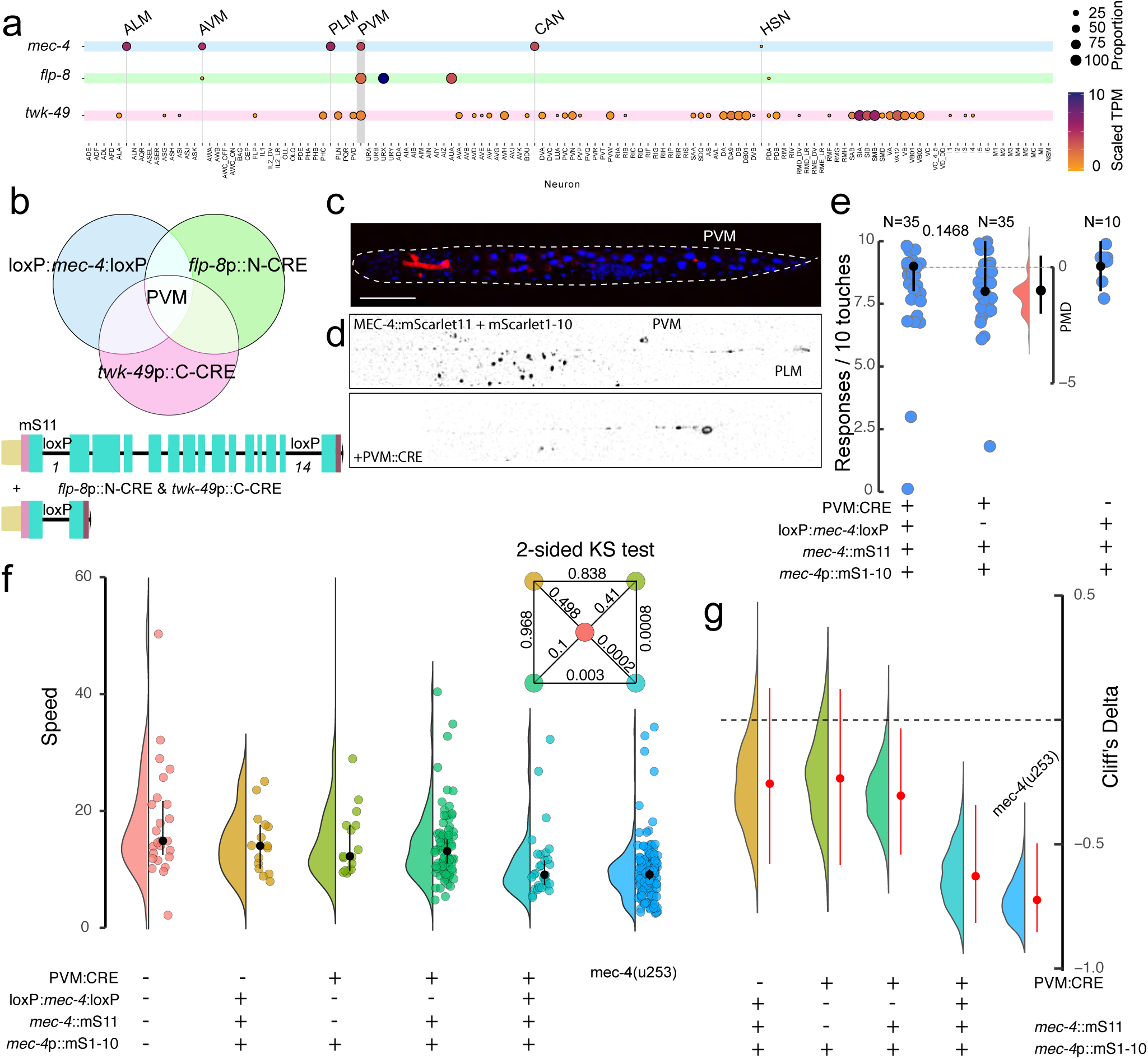
MEC-4 disruption in PVM causes lethargic locomotion phenotype. **a,** Single-cell RNA expression profile of *mec-4*, *flp-8* and *twk-49* in all neurons of adult hermaphrodites, indicating their overlap exclusively in PVM. TPM = transcripts per million. Data taken from (*19*). **b,** Design of the strategy. A split CRE recombinase will be expressed in the loxP-flanked *mec-4* transgenic under the control of *flp-8p* (N-terminal domain) and *twk-49* (C-terminal domain) promoters, leading to the disruption of the *mec-4* gene. LoxP sites were introduced by CRISPR into intron 1 and itron 14. The 5’ end is fused in frame to the 11th strand of mScarlett (ws11) to visualize protein localization upon complementation with mScarlett1-10. **c,** Representative image of a CRE reporter animal, permitting the visualization of CRE activity through a blue-to-red colorswitch. Scale bar = 100 µm. **d,** Representative images of control and PVM::CRE animal with intact and disrupted MEC-4 expression in PVM visualized by presence or absence of mScarlet. **e,** Average touch-evoked escape responses to a sequence of 10 consecutive touches in control animals versus animals in which *mec-4* is disrupted specifically in PVM. Black circle indicates the median ± 95% confidence intervall of the median. p-value indicated on top of the bracket derived from a two-sided Mann-Whitney U-test. Number of tested animals indicated above each category. Floating axis shows the bootstrapped effect size as paired mean difference (PMD). **f,** Violin plot of the distributions of worm locomotion speeds with the conditional *mec-4* ablation exclusively restricted to the PVM neuron across different genotypes. Individual data points representing the average speed of each animal and black dots indicating medians 95% confidence internal. PVM::CRE = split CRE using the *flp-8p* and *twk-49p* promoters; mS11::*mec-4*= barrel 11 of mScarlet fused to the N-terminus of MEC-4; *mec-4*p:mS1-10 = barrels 1-10 of mScarlet expressed under the *mec-4* promoter to reconstitute the mScarlet fluorescent protein. Inset: Two-sided Kolmogorov-Smirnov test p-values comparing each pair of distributions (numbers inside the matrix), highlighting the degree of distributional overlap between genotypes. Colors correspond to genotype groups consistently across violins, estimation plots, and the KS matrix. **g,** Estimation plots of bootstrapped Cliff’s Delta (red points indicate median, red bars show 95% bias-corrected confidence intervals) for each comparison relative to the reference genotype. A dashed horizontal line at zero indicates no effect.

### MEC-4–dependent mechanosensation of traction forces shapes locomotion

Effective crawling locomotion on a substrate requires sufficient friction or traction to generate forward movement. Traction along normal direction in one part of the body is needed to generate the propulsive force required to overcome resistive friction force acting on other regions of the body (*42*). Thus, optimization becomes a paradoxical process — minimizing traction at some parts and directions of the system while maximizing it in other regions and orientations. How is this achieved in moving animals? Our data up to here suggest that TRNs serve as the neuronal substrate to sense the friction force in a velocity-dependent manner.

To visualize and quantify the worm’s interaction with its substrate while crawling, we performed a traction force microscopy experiment and imaged the displacement of fluorescent microspheres on the surface of viscoelastic PAA gels (Fig. 6a, b; Supplementary Video 8). Here, bead displacements are proportional to the friction forces between the animal and the surface. Assuming a linear resistive force model, these forces are proportional to the friction coefficients and the velocity of the body part in contact. Intriguingly, we observed that the tractions produced by the crawling animals are not homogeneous, but display greater amplitudes near the head (Fig. 6c) and higher bead displacement, thus, different tractions associated with different parts of the body (Fig. 6d, e). To interpret this spatial anisotropy in traction forces, we next turn to resistive force theory (*43*), which provides a mechanical framework to relate body motion with substrate interactions. According to resistive force theory, the perpendicular (normal) drag force experienced by *C. elegans* during movement in a viscous environment is greater than the parallel (tangential) drag force. To check the validity of this assumption, we decomposed the traction field into its components perpendicular and tangential to the worm’s centerline. In fact, we found that the force components are well correlated with the movement of the worm (Fig. 6f) and also that the vectors associated with the perpendicular motion are twice as large as the one with tangential motion. This suggests that worm locomotion can be described by applying linear resistive force theory (*43*). To find the profile of the friction forces and coefficients along the worm’s body we set up a mechanical model that includes different friction coefficients along tangential and normal directions of the worm midline (Fig. 6g, see Section 3). We used this model and an optimal control formulation (*44*) for solving the following inverse problem: infer the friction coefficients and muscle torques at each time and location from the experimental measures of worm positions. Consistent with the tractions derived experimentally (Fig. 6f) and from our model (Fig. 6g), the solution of the inverse problem reveals that friction coefficients are spatially heterogeneous along the worm’s body (Fig. 6h), suggesting that this inhomogeneity contributes to optimized locomotion. Strikingly, when the same inferred internal torques were applied to a mechanical model with homogeneous and isotropic friction coefficients, the simulated animals failed to generate substantial forward propulsion (Fig. 6i; Supplementary Video 9). This raises the question: do TRNs encode information about local substrate coupling, allowing the worm to regulate its effective traction through changes in posture and velocity?

**Fig. 6.**
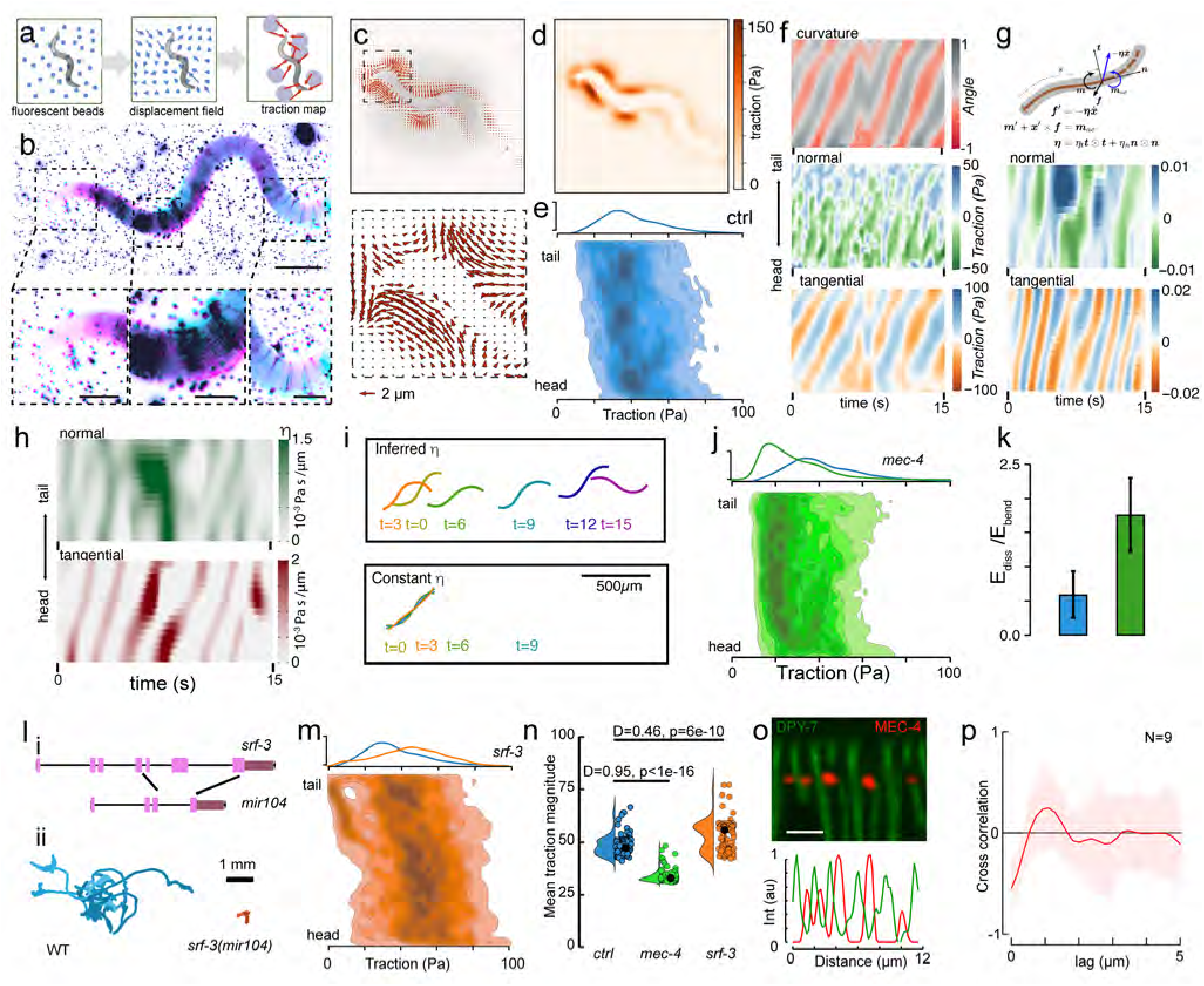
MEC-4 senses substrate traction forces. **a,** Schematic of the traction force experiment. Local displacements of the gel-embedded microspheres in the vicinity of the worm body are transformed into traction field. **b,** Representative image sequence for two successive frames false-colored from a video of a worm crawling on an elastic substrate. Scale bar = 100 µm. Insets show the displacement of the gel-embedded microspheres as the animal is crawling on the substrate. Pink, frame #*n*; Cyan, frame #*n* + 1 **c,** Representative displacement field of a worm crawling on the gel. Length of the arrows correspond to bead displacements. Inset shows magnified regions in the upper map. Scale bar = 40µm. **d,** Color map showing the uneven distribution of tractions exerted on the substrate. **e,** Quantification of the mean traction force vs body coordinate for wild-type animals (*N* = 8) on 1.25kPa PAA. **f,** Kymograph of the curvature along the body coordinate and the corresponding normal (perpendicular to the centerline, but in plane of the surface) and tangential traction map as a function of time. **g,** Explanation and formulation of the mechanical model and optimal control problem. Kymographs represent the time course of the normal and tangential tractions inferred from the worm bending/locomotion data. **h,** Representative kymographs the time course of the perpendicular and tangential friction coefficient inferred from the worm bending/locomotion data. **i,** Simulations of an animal with the inferred inhomogeneous friction coefficient and an animal with a constant friction coefficient, using the same inferred muscle torques. As shown, the homogeneous case does not lead to locomotion using the experimentally determined bending moments. See also Supplementary Video 9. **j,** Quantification of the mean traction force vs body coordinate for *mec-4* mutant animals (N = 9) crawling on E=1.25kPa substrates. **k,** Ratio of dissipated and bending energy for wild-type and *mec-4* mutant animals (N=360 points). **l,** i) Genomic organization of the *srf-3* locus and the location of the *mir104* allele. ii) Displacement of the *srf-3* and wild-type animals on 1.25 kPa substrates in the 5 min experimental window. **m,** Quantification of the mean traction force vs body coordinate for *srf-3* mutant animals (N = 9) on 1.25 kPa PAA. **n,** Comparison of the average spatial traction magnitude measured for the three genotypes on 1.25 kPa PAA gels. Each point indicates the average of each body coordinate for each strain. A linear mixed-effects model revealed significant effects of strain (*F*_2_,_·_ = 5.21), body position along the worm (*F*_1_,_·_ = 1419.26), and their interaction (*F*_2_,_·_ = 179.94) on measured traction forces. **o,** Representative confocal image and intensity distribution of a worm labeled with DPY-7::GFP and MEC-4::mScarlet to visualize colocalization of the cuticle furrows and mechanoreceptor channels, respectively. Scale bar = 5 µm. **p,** Average cross-correlation of the two signals obtained from N=9 animals.

We thus repeated the traction force experiment in the *mec-4* mutant. Interestingly, we found that the traction forces of the *mec-4* mutants were generally lower and more homogeneous than those of wild-type animals, especially on soft substrates (Fig. 6j and the Extended Data Fig. 8). However, the magnitude and distribution of the traction force was similar to that of the wild-type on stiffer substrates (Extended Data Fig. 8).Because *mec-4* mutants exhibit defects in posture and traction force generation, these findings suggest that MEC-4 is required to optimize locomotion, particularly on soft substrates. To further support this interpretation, we calculated both the bending energy and the energy dissipated during locomotion. We found that energy dissipation is substantially increased in *mec-4* animals (Fig. 6k and Methods). The elevated energy dissipation in *mec-4* mutants, combined with the higher and more spatially organized tractions in wild-type animals, suggests that mechanosensory feedback through MEC-4 improves the mechanical efficiency of locomotion.

Lastly, we sought to identify the molecular components that mediate frictional coupling between the worm and its substrate, providing a physical basis for the generation of traction forces. The cuticle of *C. elegans* is coated with an elaborate glycocalyx (*45*). SRF-3, a nucleotide sugar transporter in the Golgi apparatus, is expressed in hypodermal seam cells, leading to altered cuticle glycoconjugates on the animal’s surface. While some *srf* mutants show severe locomotion defects and fragile cuticles, *srf-3* mutants are touch sensitive (8.4±1, mean±SD, N=30 animals) and appear wild-type despite altered surface antigenicity (*46*). Using CRISPR/Cas9, we created a *srf-3* loss-of-function allele (Fig. 6l) which led to strong cuticle staining with wheat germ agglutinin (Extended Data Fig. 9a,b) as described before. Even though crawling speed on agar was slightly lower (Extended Data Fig. 9c), neither body shape nor cuticle structure was affected by *srf-3* loss of function, as determined by quantifying *dpy-7* distribution, a label for cuticle furrows (Extended Data Fig. 9d-f). We then tested their locomotion on PAA gels using traction force microscopy. Notably, *srf-3* mutants struggled to move forward (Fig. 6l), and generated higher traction across all PAA gels independent of their stiffness, suggesting increased friction forces with the substrate (Fig. 6m, Extended Data Fig. 8).

Importantly, the traction between the different strains varied significantly along the body of individual animal (linear mixed effects model, see Methods; body position: F = 1419.26, p *<* 0.001) and differed between strains (strain: F = 5.21, p *<* 0.01). Importantly, the interaction between strain and body position was significant (strain × body position, F = 179.94, p < 0.001), indicating that the strains differed significantly in spatial traction distribution along the body, not merely a global shift in traction magnitude (Fig. 6n). Visual inspection of the predicted marginal means showed that strains diverged primarily in the head and midbody regions, while converging near the tail. These results suggest that mechanical interaction of the body wall is modulated by *srf-3* and *mec-4* sensory dependend, but differences are localized rather than uniform across the body.

Intriguingly, despite the strong traction of the animal with the substrate, due to the low velocity of their movement we did not observe calcium fluctuation in PVM (Extended Data Fig. 10). This indicates that the cuticle-substrate interaction affects calcium signalling in TRNs during locomotion.

Lastly, in an effort to understand how animal use their TRNs to sense surface mechanics, using fluorescent marker strains, we quantified the relative distribution of MEC-4 ion channels and DPY-7 cuticular collagens, a marker for circumferential furrows (*47*). We found MEC-4 localized in discrete puncta that on average did not colocalize with the furrow marker DPY-7 (Fig. 6o, p), suggesting that MEC-4 colocalizes with the cuticle ridges. This observation evokes an intriguing model: given that MEC-4 ion channel complexes aligned beneath circumferential furrows, it is plausible that cuticle ridges act as sites of frictional force transfer from the substrate, thereby channeling mechanical stresses directly to MEC-4.

## Discussion

Our findings reveal that *C. elegans* use their gentle mechanoreceptor neurons to sense mechanical stresses rates during crawling locomotion. We demonstrated that these neurons actively detect forces between the body and the substrate, fine-tuning postural gait through proprioceptive feedback. Active sensing involves organisms or systems that generate movements to obtain sensory information about their environment, rather than passively receiving input (*48*). In active sensing, the organism or system takes control of how it explores and interacts with its surroundings to optimize information gathering. Animals such as rodents move their whiskers to sense textures (*49*), while human fingertip ridges enhance high-frequency vibrations during contact, improving tactile perception (*50*). Similarly, we here propose that *C. elegans* touch receptor neurons are active sensors to facilitate substrate sensing, during which forward motion helps to gather detailed information about the mechanics of the environment. Friction forces are complex and depend on the contact area (thus load), asperities of the interface and surface chemistry (*51*). The structured cuticle of *C. elegans* may therefore act similar to the ridges on human fingertips to increase friction forces between the skin and the surfaces, providing dynamic frequency-dependent sensory input. In ventral mechanosensors of wild-type animals such as PVM and AVM, MEC-4 aligns with the cuticle annuli (ridges between furrows, Figure 6o, p). Because these cuticle structures are absent in the lateral neurons (which we found to be less sensitive to substrate mechanics), we propose that mechanical forces are conveyed through the cortical fibrous organelles, cable-like structures of the epidermis that are aligned with the cuticular annuli, and thereby coupled to mechanosensitive ion channels in the touch receptor neurons (*52*).

TRNs respond rate-dependently, making them largely unresponsive to slow or stationary stimuli (*8, 17*). They only activate during a change in stimulus, or, movement. When an organism moves across a surface, it generates traction forces that vary with velocity. This traction force is due to the relative velocity of the substrate and changes in space and time. We propose that it is this change in traction, caused by movement, that generates a dynamic force on neurons. This time-varying force prevents the mechanosensitive channels from closing (*17*), keeping enough of them in an active state to ensure that a strong signal is produced. Mechanically, our simulations demonstrate two necessary frictional conditions for effective crawling: the tangential friction coefficient (and tractions) must be low enough to overcome drag and normal friction must be high enough to provide propulsion. Wild-type worms seem to modulate these two conditions efficiently by sensing and adapting their traction force. Intriguingly, lab cultivated strains seem to have lost the higher effective drag anisotropy, which means the wild isolates crawl with less surface slip (*43*). Some partial explanation for this modulation may be found by resorting to more sophisticated friction laws, such as lubrication theory (*53*), where frictional coefficients depend on the curvature of the groove and the worm cross-section. Consequently, by changing the curvature of the cross-section while bending, the worm may be able to modify the contact area and the drag coefficient ratio between normal and tangential friction coefficients. *mec-4* animals are sensory deficient and thus are not able to properly synchronize their muscle activity with their curvature profile. Interestingly though, *srf-3* mutants may crawl less on all substrates assay, despite having a higher traction force. This points to a scenario where the defective glycocalyx leads to stonger adhesion/friction of the animal with the substrate such that it appears to move less.

We have shown that *C. elegans* actively produces inhomogeneous traction on elastic substrates and thus shares the characteristic with other limbless animals, such as snakes (*54*). Because the highest traction is exerted in the anterior part, animals can use the muscles in the head to actively steer toward the substrate to maximize sensory information. In the absence of this information the *mec* mutants ‘loose orientation’, do not actively adjust friction forces and cease crawling.

In absence of food on soft viscoelastic substrates, body wall mechanoreceptors, especially PVMs, exhibit velocity-dependent activity consistent with sensing substrate mechanics; the magnitude and expression of these phenotypes are modulated by feeding state, sex, and substrate. Why are TRNs, and PVM in particular, exquisitely sensitive to soft substrates but not to stiffer substrates? TRNs are not the only body-wall mechanosensors, and PVD neurons are known as high-threshold mechanosensors. This lower sensitivity may, therefore, enable mechanosensation on stiffer substrates. Together, a degenerate circuit system (*55*) and molecules with increasing sensitivity may therefore ensure optimal locomotion in varying and dynamic mechanical environments.

## Acknowledgments

The authors thank the NMSB lab and Benjamin Podbilewicz for discussions and comments on the manuscript and the *Caenorhabditis* Genetics Center for providing some strains (CGC, supported by the National Institutes of Health, P40 OD010440). Mechanical and electronic workshops at ICFO for manufacturing customized parts.

## Funding

AP acknowledges financial support from Grant FJC2021-047089-I funded by MCIN/AEI/ 10.13039/501100011033 and by the European Union “NextGenerationEU/PRTR”, MK acknowledges financial support from the Human Frontiers Science Program (RGP021/2023), MCIN/ AEI/10.13039/501100011033/ FEDER “A way to make Europe” (PID2024-157334OB-I00, CNS2022-135906), “Severo Ochoa” program for Centres of Excellence in R&D (CEX2024-001490-S)[MICIU/AEI/10.13039/501100011033], from Fundació Privada Cellex, Fundació Mir-Puig, and from Generalitat de Catalunya through the CERCA and Research program. ICFO is the recipient of the Severo Ochoa Award of Excellence of MINECO (Government of Spain). JJM and AB acknowledge Severo Ochoa program (CEX2018-000797-S). JJM is also financially supported by grant PID2020-116141GB-I00 funded by MCIN/AEI, and grant SGR 01049 from the local government Generalitat de Catalunya.

## Authors contributions

Investigation - including calcium imaging in freely moving worm - AP. Building the microscope - CA, AP. Locomotion experiments - AP, RD, ET, MPR. Traction force microscopy - AP. Optogenetics - AP. Strain generation - AP, MPR, CMF, NSC. Confocal microscopy - AP, MPR, NSC. Microfluidics - AL, ACS, MK. Connectome modeling - AP. Implementing the software for close-loop experiment - AP. Formal analysis - AP, MK, AB and JJM. Methodology - AP, JJM, MK. Conceptualization - MK, JJM. Funding acquisition - MK, JJM. Supervision - MK, JJM. Writing, original draft - MK, AP. Visualization - MK, AP.

## Competing interests

The author have no competing interests to declare.

## Data and materials availability

All numerical source data is presented in the supplementary material of the manuscript or available from the corresponding author. Curvature matrices were deposited on zenodo.org (doi: 10.5281/zenodo.18153808).

## Code availability

The codes are deposited on: https://gitlab.icfo.net/NMSB/traction-force-worm

https://gitlab.icfo.net/NMSB/track-live-close-loop

https://gitlab.icfo.net/NMSB/net-sim-worm

https://gitlab.icfo.net/NMSB/cal-free-m-worm

## 3 Extended Data Figures

**Extended Data Fig. 1.**
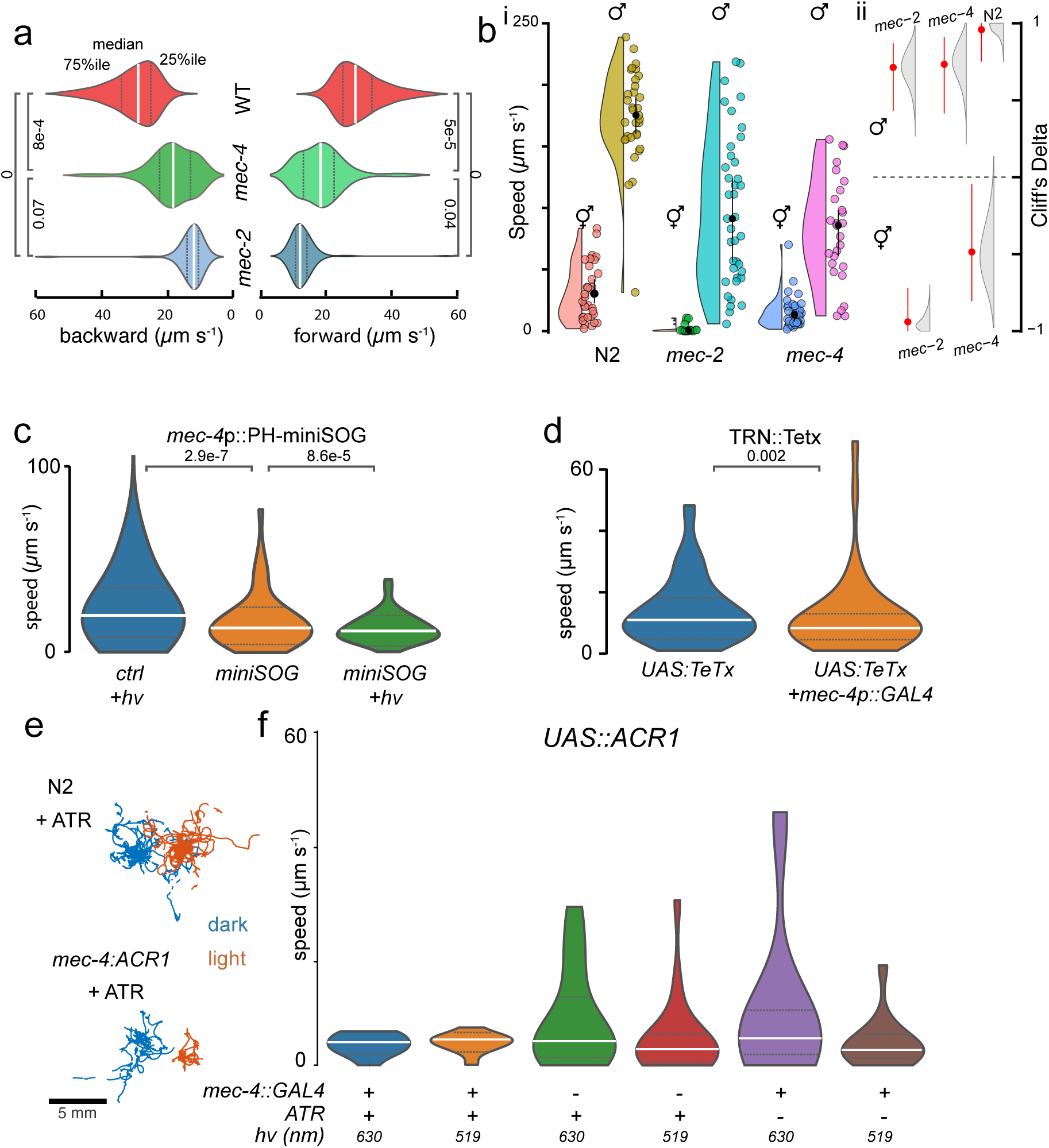
Disruption of gentle ‘touch’ receptors causes lethargus. **a,** Violin plots showing the distribution of the backward and forward velocity. White lines show medians; dotted lines represent the 25th and 75th percentiles. *p*-values for linear mixed model for nested comparisons, comparisons between dispersion/forward/backward velocity respectively, significant in red. ctrl vs mec-4:3.90 × 10^−18^*/*1.27 × 10^−28^*/*1.02 × 10^−29^ ctrl vs mec-2: 1.23 × 10^−43^*/*2.59 × 10^−123^*/*2.31 × 10^−131^ mec-4 vs mec-2: 4.63 × 10^−3^*/*3.35 × 10^−13^*/*1.36 × 10^−14^ **b,** i) Distribution of the crawling speed comparing wild-type hermaphodites and males with males and hermaphrodites from *mec-2* and *mec-4* mutant alleles. P-values above two neighboring distribution were derived from a two-sided Dunn-test with Bonferroni adjustments for multiple comparison. For exact number of measurements and replicates, consult Supplementary Table 3. ii) Estimation plots of boot-strapped Cliff’s Delta (red points indicate median, red bars show 95% bias-corrected confidence intervals) for each comparison relative to the reference genotype. A dashed horizontal line at zero indicates no effect. Animals were tracked from 2-min videos using the Manual Tracking plugin in ImageJ (v1.54p). Euclidean distances between consecutive x,y-positions were calculated every second in Python. Instantaneous velocity was computed as distance divided by the 1-s interval, and mean velocity per animal was plotted as violin plots, with individual data points shown for each worm. **c,** Chronic ablation of TRNs using miniSOG and blue light. Blue violin indicates the wild-type animals under 4 min of continuous illumination of blue light. Orange violin indicates miniSOG expressed in TRNs under the *mec-4* promoter but kept in the dark before recording behavior under red light. Green violin indicates miniSOG expressing animals under continuous illumination with blue light. **d,** Average crawling speed of animals expressing tetanus toxin light chain (UAS::TeTx) in TRNs using the *mec-4*p::GAL4 driver. Blue violin indicates UAS:TeTx effector alone; orange violin indicates combined expression of TRN:Gal4 driver and TeTx effector. TRN silencing reduces movement compared to controls. **e,** Individual locomotion trajectories for wild-type N2 animals and animals expressing ACR1 in touch receptor neurons, supplemented with ATR. Each mutant strain exhibits reduced locomotion relative to wild-type upon illumination, consistent with impaired TRN function. **f,** Acute suppression of TRN activity leads to immediate lethargus. Average crawling speed of animals expressing Anion channelrhodopsin 1 (ACR1) in TRNs under the indicated combinations of UAS::ACR1, *mec-4*p::GAL4, ATR (±) and illumination wavelength (h*ν* = 519 nm or 630 nm). Conditions that permit light-activated ACR1 activity (ATR present + appropriate wavelength) show a pronounced decrease in crawling speed versus matched controls (ATR- or non-activating wavelength), demonstrating wavelength- and cofactor-dependent modulation of behavior. For all panels, white lines show medians; dotted lines represent the 25th and 75th percentiles.

**Extended Data Fig. 2.**
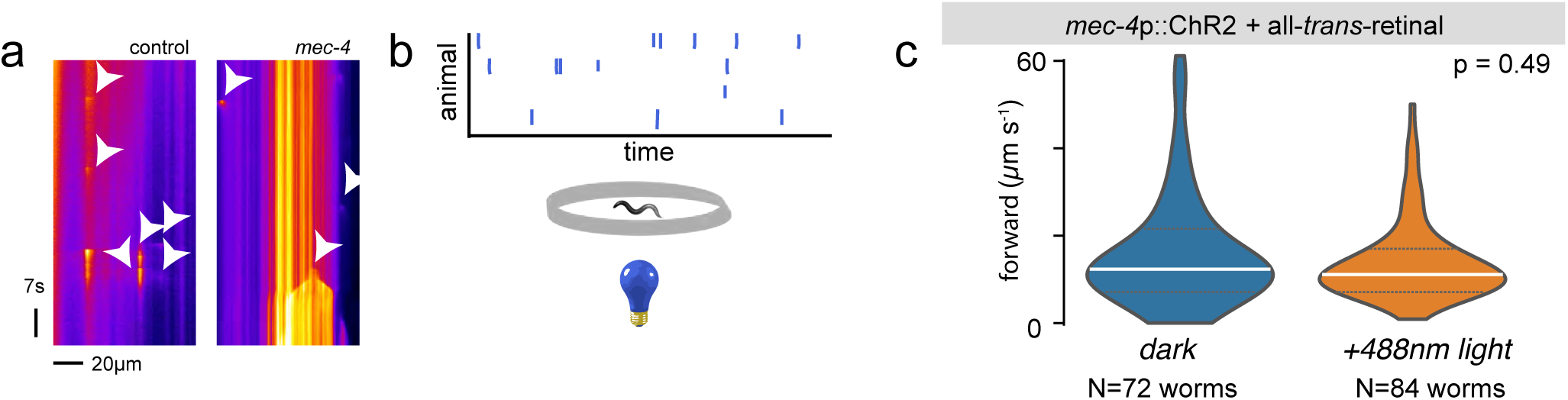
Random, ectopic activation of TRNs does not suppress *mec-4* induced lethargus. **a,** Spontaneous calcium fluctuations in PLM neurites in immobilized wild-type (left) and *mec-4* (right) mutant animals **b,** Schematics of the experiment, where a cohort of animals is illuminated with a Poisson-distributed stimulus for 48h. **c,** Average crawling speed of *mec-4* animals expressing Channel-rhodopsin2 in TRNs under the indicated stimulation protocol. Note, even though animals immediately respond to blue light, their spontaneous locomotion behavior is unchanged, indicating that occasional neuronal activity throughout larval development is not sufficient to suppress the *mec-4* induced lethargus.

**Extended Data Fig. 3.**
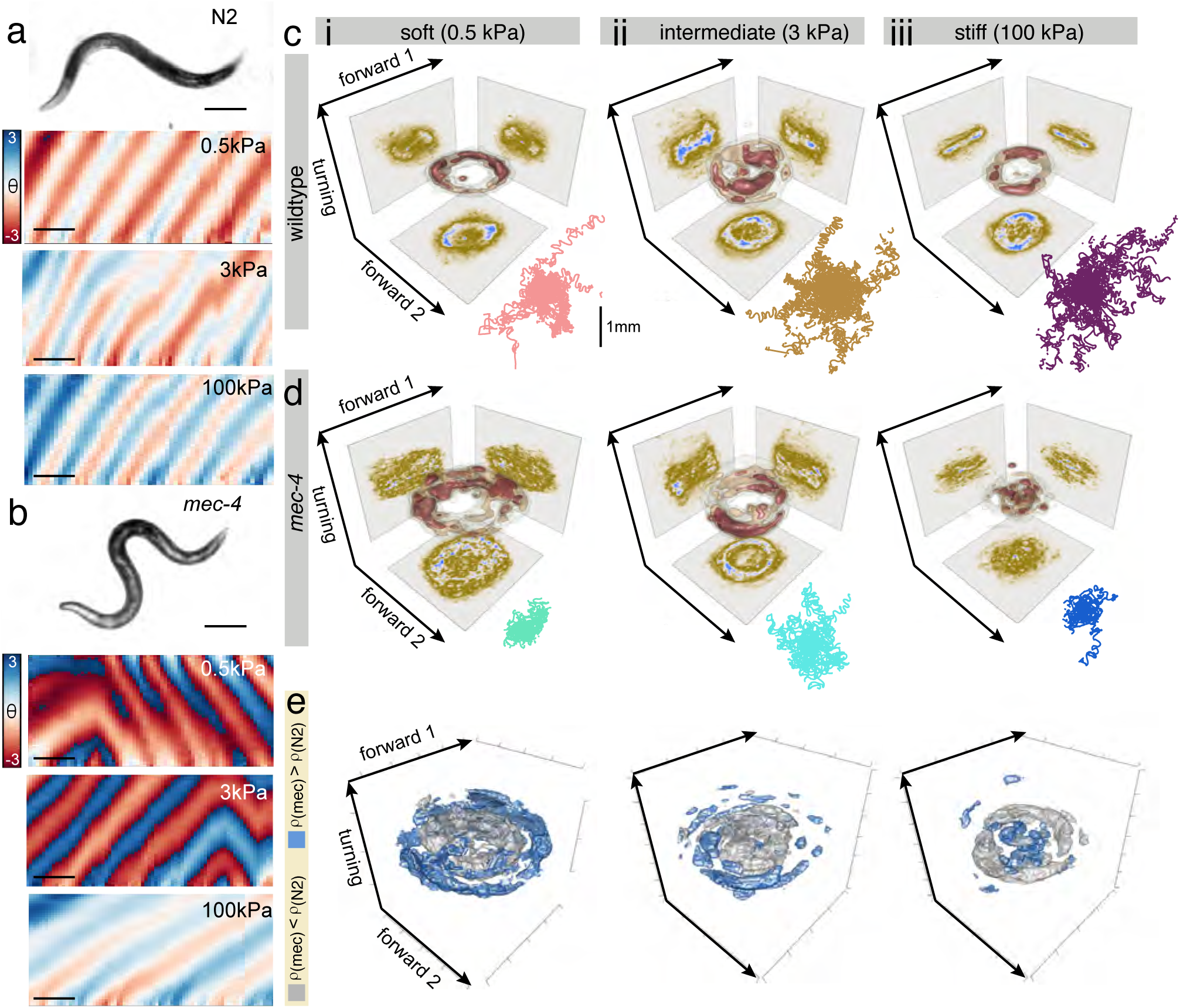
Animals adjust their gait on soft substrates in a MEC-4 dependent manner. **a, b,** Representative photograph of a wild-type (a) and *mec-4* mutant (b) animal crawling on soft substrates (0.5 kPa, scale bar = 100 µm). Representative curvature kymographs for the three different elastic matrices tested are shown below. Scale bar = 2s. **c, d,** Locomotion of (c) wild-type and (d) *mec-4* mutant animals on three different polyacrylamide gel substrates with increasing stiffness. Joint probability distribution (equivalent to a discrete 3D histogram) of the two forward and turning modes in the eigenworm space. Color scale = (brown, low density; blue, high density). A larger expansion of the manifold indicates higher body curvature during locomotion. Colored lines below the 3D histogram indicate the trajectories of the animal displacements. **e,** 3D plot of the statistically significant differences comparing the joint probability distribution shown in (c) and (d), comparing control and *mec-4* mutant animals crawling on soft (0.5 kPa), intermediate (3k Pa) and stiff ((100 kPa) substrates. Silver voxel indicates higher density for N2 control animal, blue voxels indicate higher density for *mec-4* mutants with *p <*0.01.

**Extended Data Fig. 4.**
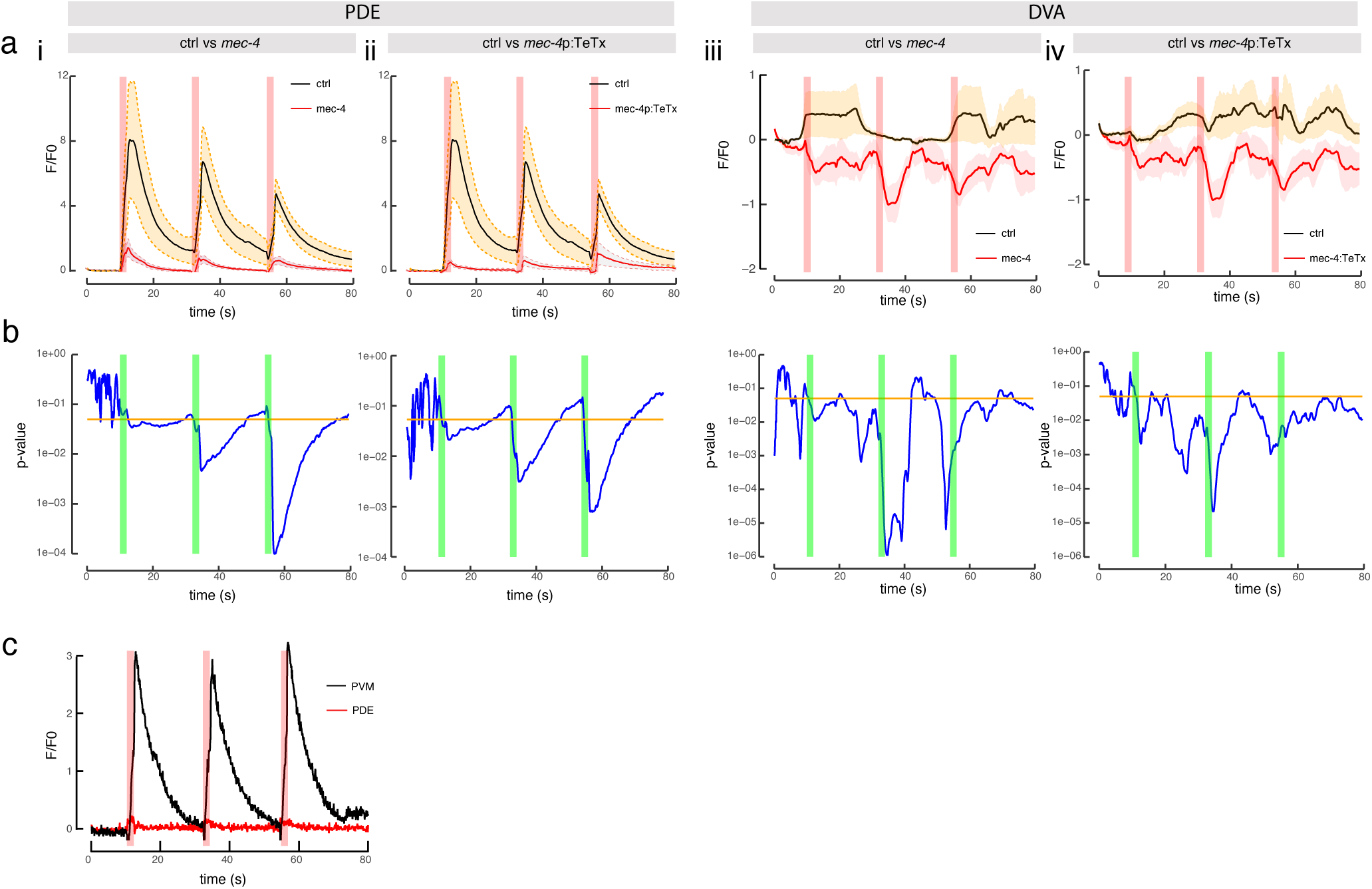
Statistical comparison of the calcium response after direct mechanical stimulations. **a,** Plots showing the average fluorescence intensity of the calcium recordings between the indicated genotypes. Recordings of PDE in (i) ctrl vs *mec-4* and (ii) ctrl vs *mec-4p*:TeTx expressing animals. Recordings of DVA in (iii) ctrl vs *mec-4* and (iv) ctrl vs *mec-4p*:TeTx expressing animals. **b,** Graphs showing a one-sided p-value (left axis) derived from a t-test statistics and the average calcium dynamics (right axis) vs time (bottom) for the indicated comparisons. The running, time-dependent *p*-value is indicated as blue line, the alpha-level of significance indicated as orange line. We consider the recorded time point different to the control when the blue line drops below the orange level of significance. **c,** Representative recording of PVM and PDE in a mec-4p::TeTx transgenic animal.

**Extended Data Fig. 5.**
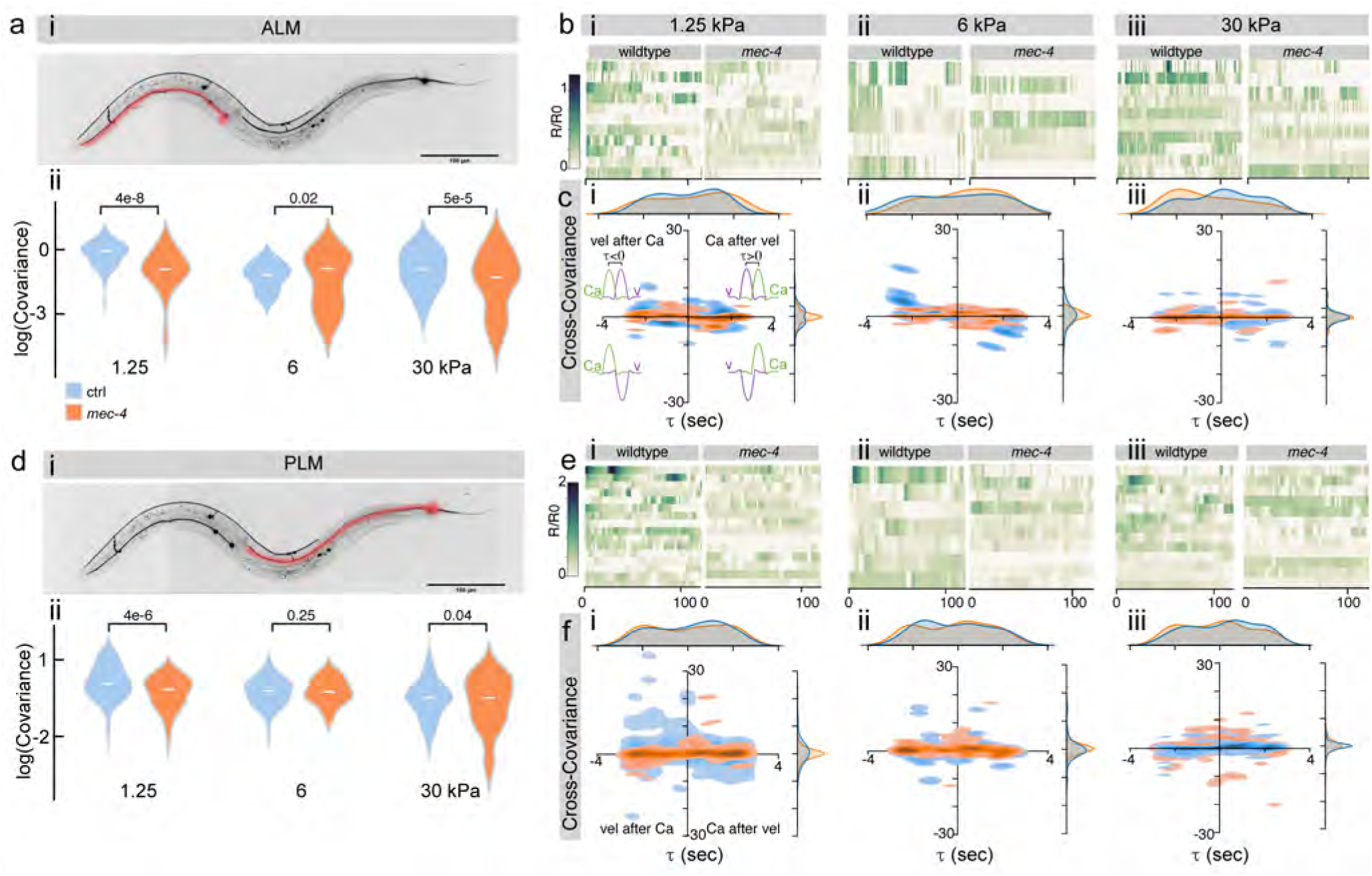
ALM and PLM calcium imaging in freely moving animals on different substrates. **a,** (i) Sketch of an animal with ALM highlighted in red. (ii) Covariance between forward velocity and calcium activity in ALM. p-values derived from a two-sided Wilcoxon rank-sum test. **b,** Kymographs of wild-type and *mec-4* mutant ALM activity during free locomotion on PAA gels with different stiffness. i) 1.25kPa; ii) 6kPa; iii) 30kPa. **c,** Cross-Covariance of ALM activity and forward locomotion velocity crawling on i) 1.25kPa; ii) 6kPa; iii) 30kPa. hydrogels. Inset inside the four quadrant of panel (i) indicates the temporal relation between the velocity and the calcium signals. **d,** i) Sketch of an animal with PLM highlighted in red. (ii) Covariance between forward velocity and calcium activity in PLM. p-values derived from a two-sided Wilcoxon rank-sum test. **e,** Kymographs of wild-type and *mec-4* PLM activity during free locomotion on PAA gels with different stiffness. i) 1.25kPa; ii) 6kPa; iii) 30kPa. **f,** Cross-Covariance of PLM activity and forward locomotion velocity crawling on i) 1.25kPa; ii) 6kPa; iii) 30kPa hydrogels.

**Extended Data Fig. 6.**
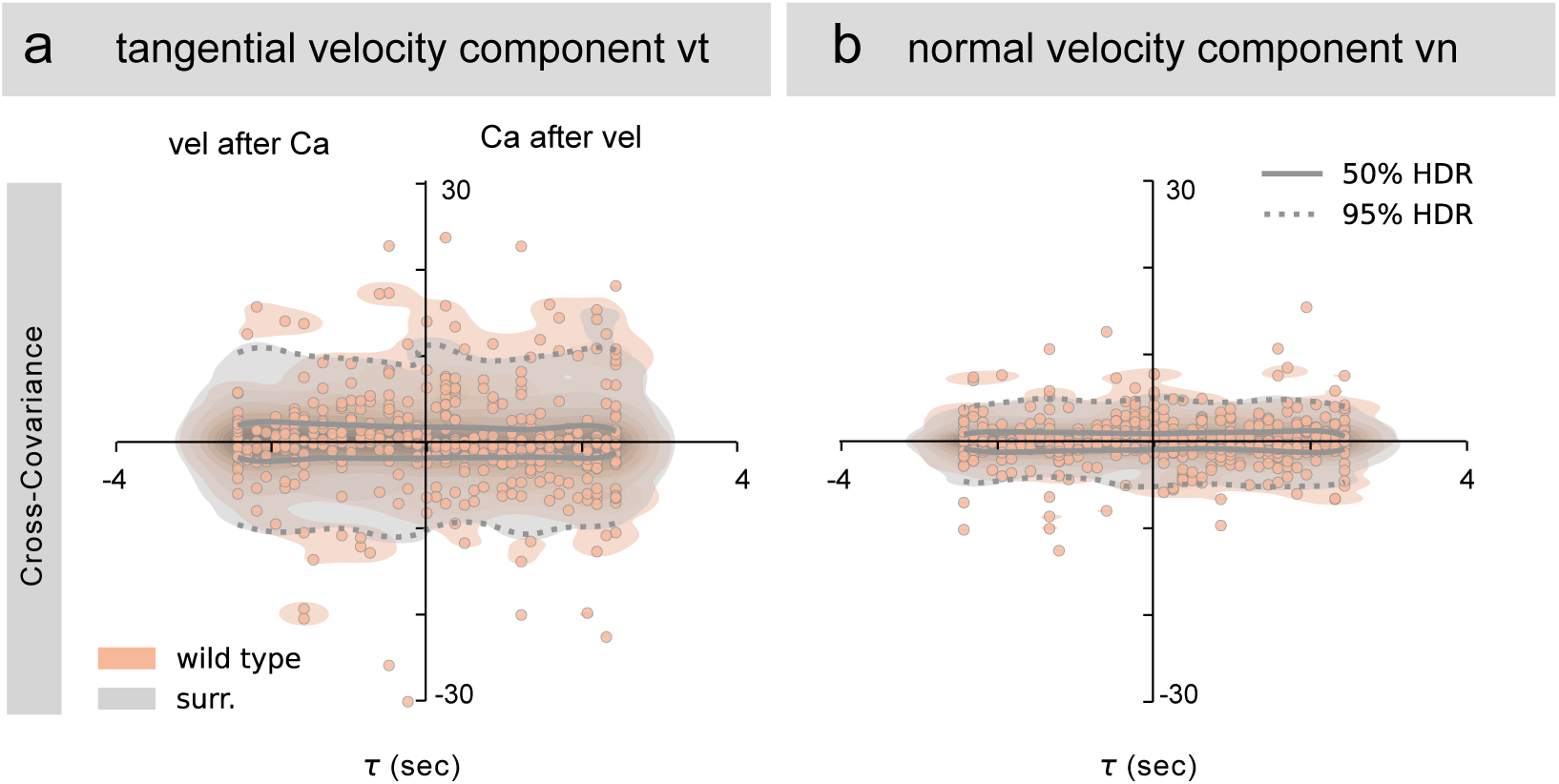
PVM activity co-varies with tangential (and not normal) components of velocity. **a, b**, Cross-covariance plots between PVM calcium dynamics and velocity components indicating the position of the cross covariance extrema for 5-seconds windows for wild-type worms and inter-subject surrogates for (a) tangential, v*_t_* and (b) normal, v*_n_* velocity components respectively. Surrogate analysis indicates that the local extrema in (a) are due to a real covariance between the time series, rather than due to internal properties of the individual time series.

**Extended Data Fig. 7.**
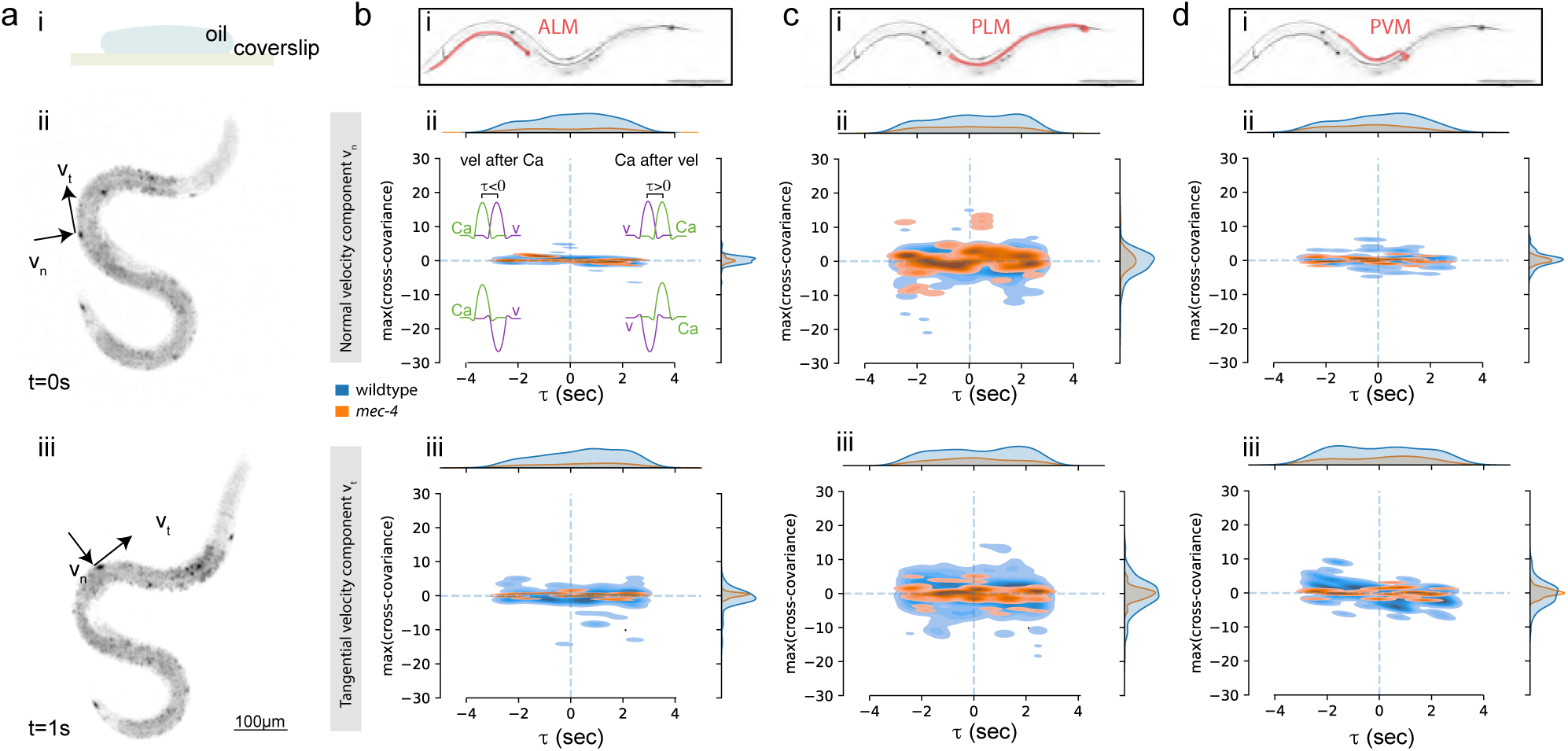
Viscous drag does not activate body wall mechanosensors. **a,** i) Schematic of the experiment of an animal immersed inside a flattened drop of Halocarbon oil. Representative snapshots of an animal expressing GCaMP6s in TRNs crawling inside the oil droplet at two different time-points 1 s apart. The net velocity was decomposed into the normal and tangential component to visualize causal dependence between viscous drag and calcium activity. **b-d,** i) Schematic representing the (b) ALM, (c) PLM and (d) PVM neuron. ii-iii) Plots showing the maximum cross-covariance of the calcium signal with the (ii) normal and the (iii) tangential velocity component. Blue, control; orange, *mec-4*. Negative timelags indicate calcium transients that precede bouts in velocity (e.g. due to self-inflicted body touch), whereas a positive timelag indicates that calcium transients follow bouts in velocity. Inset inside each quadrant of panel (b ii) indicates the temporal relation between the velocity and the calcium signals.

**Extended Data Fig. 8.**
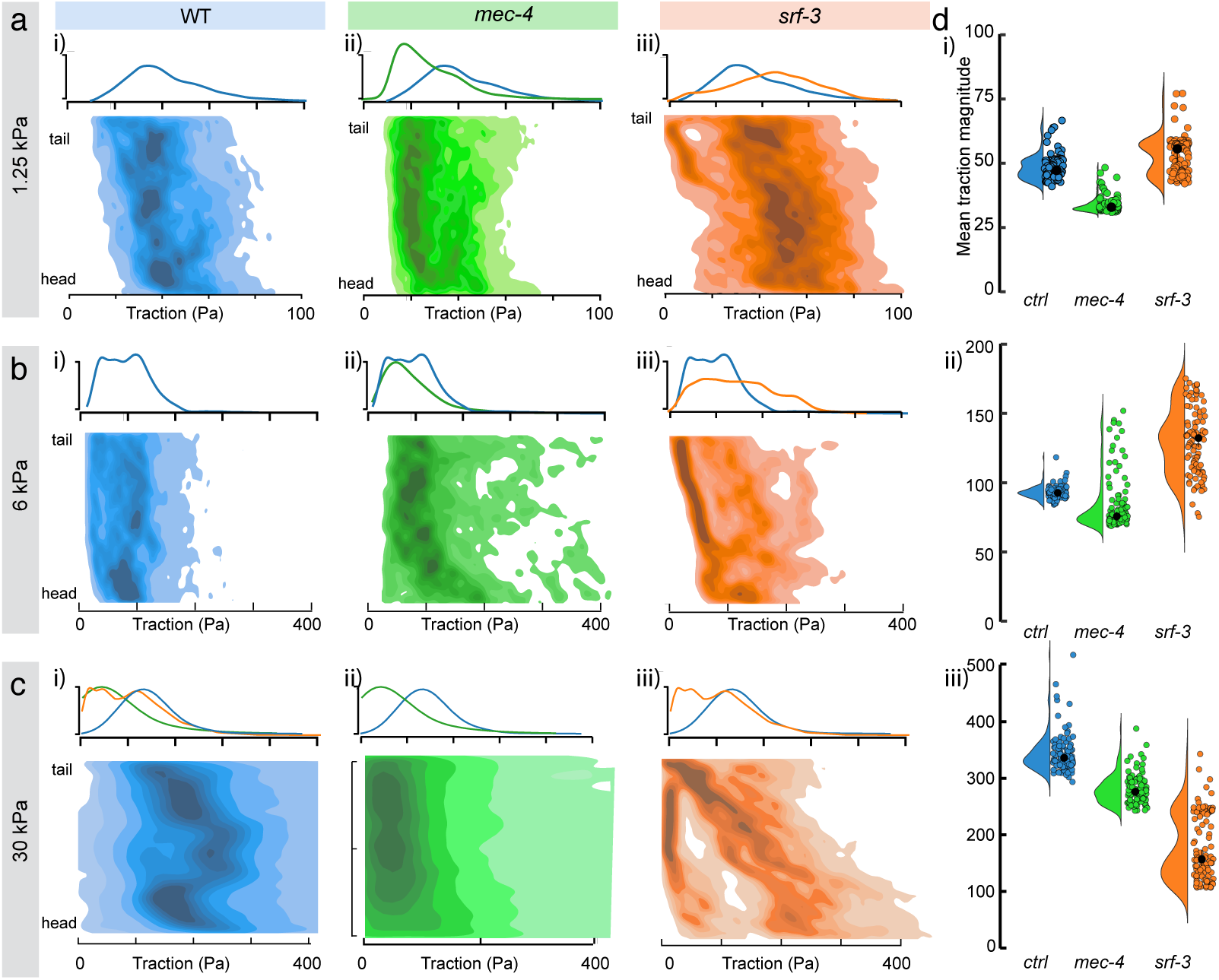
Traction forces of *mec-4* and *srf-3* mutants on different substrates. **a-c,** Distribution of traction forces along the body coordinate for (i) wild-type, (ii) *mec-4* and (iii) *srf-3* animals crawling on 1.25 kPa (a), 6 kPa (b) and 30 kPa (c) stiff polyacrylamide gels. Note, tractions and gel stiffness have the same units for stress and Young’s modulus, respectively. Densities correspond to number of observations in a smoothed KDE. **d,** Statistical comparison of the tractions per strain and along body coordinate using a linear mixed-effects model. Each point indicates the average of each body coordinate for each strain. (i–iii) Analysis revealed a significant effect of body position (all p *<* 2 × 10^−16^) and a significant strain × body position interaction at all substrate stiffnesses (all p *<* 2 × 10^−16^). A main effect of strain was observed only at i) 1.25 kPa (F_2_,_23_._4_ = 10.3, p = 6 × 10^−3^), but not at ii) 6 kPa (F_2_,_18_._1_ = 2.98, p = 0.076) or iii) 30 kPa (F_2_,_18_ = 0.24, p = 0.76). Body position effects were: i) 1.25 kPa: F_1_,_2571_ = 1416.1; ii) 6 kPa: F_1_,_2076_ = 1319.1; iii) 30 kPa: F_1_,_2076_ = 400.2. Interaction effects were: i) 1.25 kPa: F_2_,_2571_ = 179.9; ii) 6 kPa: F_2_,_2076_ = 279.8; iii) 30 kPa: F_2_,_2076_ = 218, indicating that strain-dependent differences in traction are spatially localized along the body axis.

**Extended Data Fig. 9.**
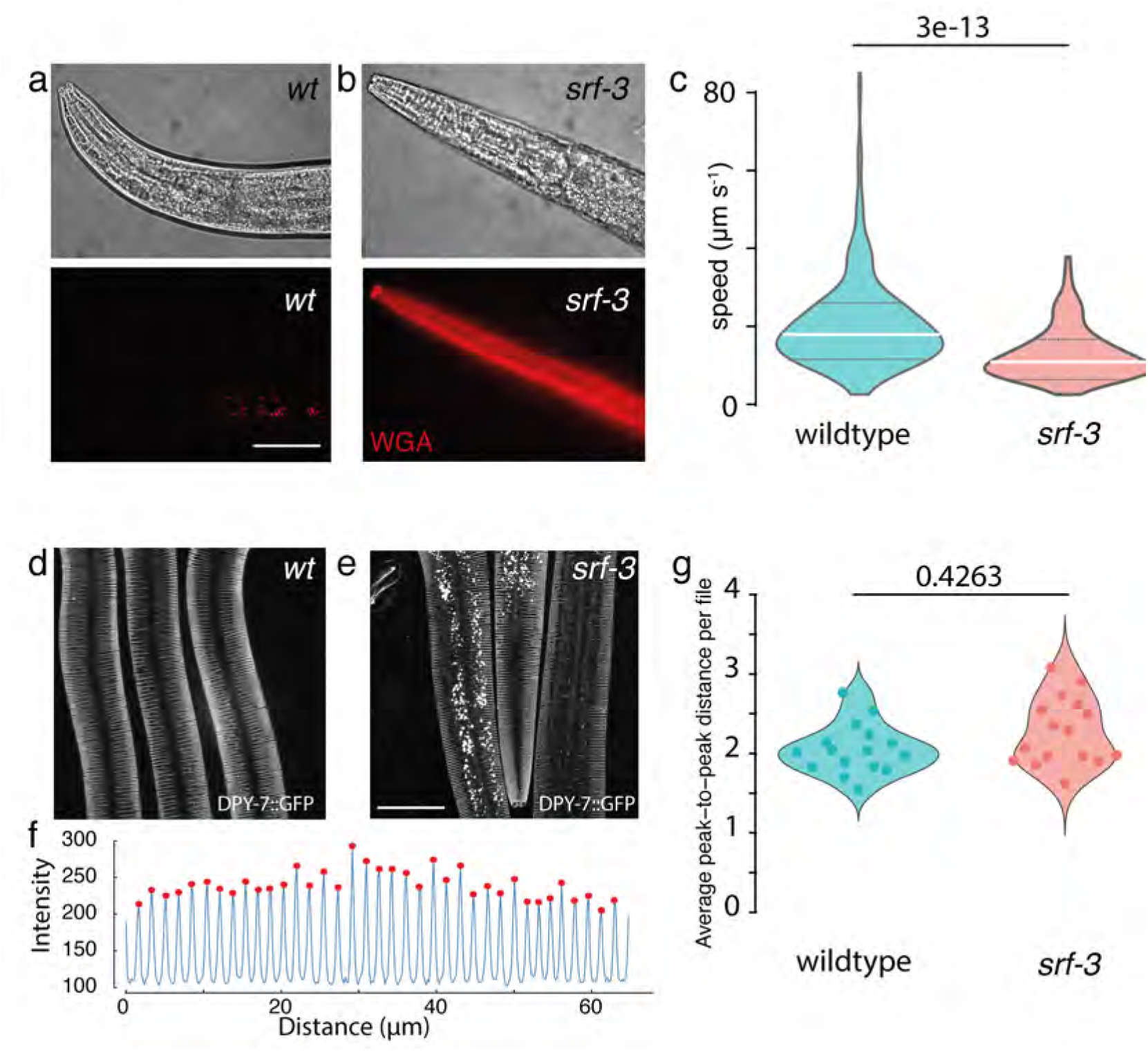
Characterization of *srf-3* mutant. **a, b,** Staining of (a) wild-type and *srf-3(mir104)* animals with wheat germ agglutinin (WGA). Note, the absence of the *srf-3* - dependent gly-cocalyx leads to strong WGA staining of the collagen cuticle. **c,** Comparison of the crawling speed between wild-type (N=266) and *srf-3* mutant (N=112) animals on normal NGM agar. Two sided KS test. N=number of animals. **d, e** Representative images of DPY-7::GFP expressing (d) wild-type and (e) *srf-3* mutant animals. **f,** Intensity profile of the DPY-7::GFP transgenic cuticle, labeling the furrows. **g,** Comparison of the DPY-7::GFP interval between wild-type and *srf-3* mutant animals. p-value derived from the two-sided KS-test, N= number of analyzed animals.

**Extended Data Fig. 10.**
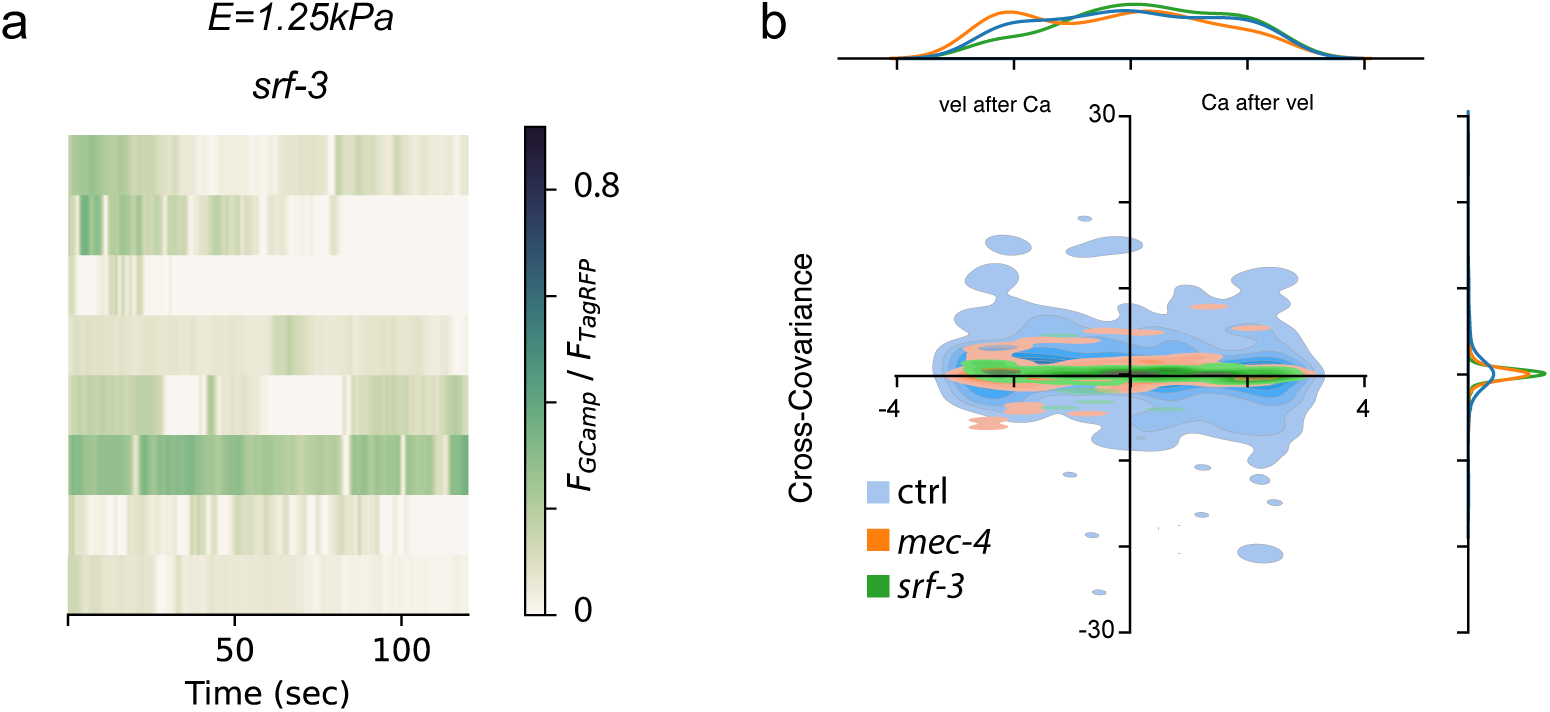
Calcium dynamics in the *srf-3* mutant. **a,** Calcium dynamics in PVM of *srf-3* mutant is absent in moving worms. Kymographs of PVM activity during free locomotion on 1.25kPa PAA gels. *srf-3* lacks characteristic activity ‘spikes’ present in wild-type. **b,** Cross-covariance plot indicating that *srf-3* lacks peaks characteristic for wild-type worms.

## Supplementary Material

## 1 Methods

### 1.1 Animal maintenance

Animals were grown and maintained according to standard protocols (*57*). Synchronization was performed as stated previously (*58*) and animals were left in the 20°C incubator until they reached the young adult stage (approximately 52 hours, subjected to strain variability).

### 1.2 Molecular biology, transgenesis and genome editing

All plasmids listed in Supplementary Table 1 were generated using the Gibson assembly method. All coding sequences were verified by sequencing. Transgenic strains were generated by microinjection into the worm gonad (*59*). When multicopy extrachromosomal arrays were integrated, the UV/TMP strategy was used (*24*). CRISPR edits were performed as described previously (*60*). MSB952 was generated using FLint as described (*61*). Unless otherwise specified, crRNAs, tracrRNAs, ssODN, gBlocks and Cas9 were purchased from IDT Technologies. All resulting strains were verified by sequencing.

#### Conditional knockout of *mec-4* in PVM

50 ng/*µ*l of each pNMSB184 and pNMSB185 (See Supplementary Table 1) were coinjected with 25 ng/*µ*l of pCFJ68 (*unc-122*p::GFP) into SV2049 worms giving rise to MSB1492. Integration of this array produced strain MSB1501.

#### Expression of tetanus toxin in TRNs

Background strains MSB273 (for DVA) or MSB661 (for PDE) were injected with 20 ng/*µ*l pNMSB186 (*mec-4*p::mScarlet::SL2::TeTx), 1 ng/*µ*l pNM50 (*myo-2*p::GFP) and 80 ng/*µ*l of 1kb Plus DNA ladder (Invitrogen) to generate MSB1505 and MSB1542 respectively. MSB1524 was the result of integrating the array in MSB1505.

#### Split mScarlet::MEC-4

To visualize endogenous MEC-4 expression, we used a split fluorescent protein approach (*41*) in which we introduced the 11th *β*-barrel of mScarlet before the first exon of the gene, with a GGSGG flexible linker between them. We used two crRNAs (#51: 5’ CGGATGGGTCCCGAAGGTGT 3’ and #52: 5’ TGTACTCGGATGGGTCCCGA 3’) targeting a few bases upstream of the first exon of *mec-4* and a ssODN (tgctattttttttatcgctatcaagttatagaatgTACACCGTCGTCGAGCAATACGAGAAGTCCGTCGCCCGTCACTGCACCGGAGGAggtggatctggaggtatgtcatggatgcaaaacctgaaaaactaccaGcaTctCcgAgaTccatccgagtacatgtcccaggtttatggagaccc) (barrel 11 UPPERCASE) with 35 bp homology region upstream and downstream of the insertion and 5 silent mutations (UPPERCASE) to prevent targeting of the crRNAs to the ssODN. The resulting strain (MSB797) was outcrossed once to the parental KG1180 to remove the dpy phenotype, leading to MSB931. It was then injected with *mec4*p::mScarlet 1–10 (pNS50, 10 ng/*µ*l) as an extrachromosomal array to restrict expression selectively to TRNs, together with a co-injection marker for green coelomocytes (pCFJ68, *unc-122*p::GFP, 30 ng/*µ*l), leading to MSB932 *mir50*(mScarlet(11)::*mec-4*); *mirEx384*(*mec-4*p::mScarlet1-10; *unc-122*p::GFP). The extrachromosomal array was UV integrated leading to MSB943 *(mir50*(mScarlet(11)::*mec-4*); mirIs93(*mec-4*p::mScarlet1-10; *unc-122*p::GFP). Both strains were sensitive to gentle touch to the body wall: MSB932 9.35±1.42; MSB943: 8.85±1.23 (mean±SEM, N=number of individual animals).

#### *mec-4*::active aptazyme//*mec-4(mir85)*

We introduced an active aptazyme (*27*) 16 bp downstream the *mec-4* stop codon by using a crRNA (#78: 5’ tttaagacacaacattgcaa 3’) annealing at the 3’ UTR of the gene. A 35 bp homology arms, double stranded DNA amplified from a gBlock with KOD Hot Start DNA Polymerase (Novagen, ref. 71086) was used as repair template (5’ tgcgtctggatctttctgaatttgttttttcttgtCAAACAAACAAAGGCGCGTCCTGGATTCGTACAAAACATACCAGATTTCGATCTGGAGAGGTGAAGAATACGACCACCTGTACATCCAGCTGATGAGTCCCAAATAGGACGAAACGCGCTCAAACAAACAAAtttaaagttaccattgcaatgttgtgtcttaaaataaaaatttacatgagaataatt 3’, aptazyme in UPPERCASE). With this approach we generated two strains, MSB1137 on wild-type background, and MSB1549, with the mScarlet::*mec-4* background (MSB943). Functionality of the aptazyme to reduce RNA levels was tested in RT-PCR on MSB1137 (Suppl. Fig. 1b, c). Briefly, worms incubated for 5 hours with the stated Tetracycline concentration were pelleted and washed twice with M9. 7 volumes of TRIReagent (TR-118 MRC) were added to the packed pellets and sored at −80°C. Next, 5 cycles of freeze/thawing/vortexing were performed and samples allowed to stand at room temperature (RT) for 5 minutes. Addition of 0.2 ml of chloroform per ml of Trizol used was carried out before a 2-3 minute RT incubation followed by a 15 minute centrifugation at 4°C and 12000xg. The aqueous phase was transferred to a new tube and 0.5 ml of isopropanol per 1 ml of Trizol added. after a 10 minute RT incubation, samples were centrifuged for 15 minutes at 12000xg and 4°C. Supernatant was then removed and pellet washed with 1 ml 75% ethanol centrifuged for 5 minutes, 4°C and 7500xg. Supernatant was removed and pellet allowed to air-dry before using RNAse-free water to resuspend. Quality and quantity was assessed on a NanoDrop with serial dilutions. RT-PCR was performed using Superscript IV One-step RT-PCR System (Invitrogen, ref. 12595100). Primers for *mec-4* amplification were FW 5’-TCTGACTCATATTGACCCTGCGTTTGG-3’ and RV 5’-TGGTTGTTGGCATCTGCAACGG-3’, giving rise to a 540 bp fragment. For *dat-1* amplification, primers were FW 5’-AGAACAGTGGTCTGGAAAGCTGGAC-3’ and RV 5’-GCCCATGGCAGGTTAAAACTGAATGAAG-3’ and the resulting amplicon was 350 bp long. The recovery of the mScarlet fluorescence signal (MSB1549, Suppl. Fig. 1d) was performed as previously reported (*41*). Briefly, adult worms were allowed to lay eggs on 10 *µ*M tetracyline plates and the F1 mounted, once young adults (YA, 20°C), on 2% agarose pads in 3 mM levamisole (*58*). Fluorescence images were taken using an inverted confocal microscope (Nikon Ti2 Eclipse) with a 60×/1.4 NA oil immersion lens. mScarlet was excited using the 561 nm laser at maximum laser power and transmitted through a 594 nm emission filter. Exposure time was 200 ms. 100–200 ms, depending on the strain to image.

#### Deletion allele *srf-3(mir104)*

We generated a deletion mutant for *srf-3* in the wild-type background covering from exon 5 until the end of the gene. We used two crRNAs, one targeting intron four (#87:5’ gtttgtgtgtaatgatgacg 3’) and the other at the end of the last exon (7, #88: 5’ CGAGATTCCGTCTA-CAAAAG 3’. We also used a repair template ssODN (5’ taatatacagacgagaatttacagacaaaacgcctTtAtCacacTcaaaccatgaaagttgcggcgtctaaatgtgaagctg 3’) with 35 bp homology regions and four mutations (UPPER-CASE) to generate a premature stop codon. However the sequencing results reported a slightly different repair, still generating a premature stop codon: 5’ taatatacagacgagaatTTACAGACAAAACGCCTcTTAGACGTATCTCATTACGATCATCACGATCATCGATCATTACGATCATCATTACACACaaacCATGAAAGTTGCGGCGTCTAAATGTGAAGCTG 3’

#### Deletion allele *mec-10(mir110)*

We proceeded to generate a deletion mutant for *mec-10* in the *mec-4(u253)* background (TU253). We used two crRNAs, one targeting in the first exon (#81:5’ GGTTGAAATTTTGACATTCG 3’) and another overlaping the last exon and the 3 ‘UTR of the gene #82: 5’ GAgagccatgagaaaaaagg 3’. We also used a repair template ssODN (5’-tccgttgattctgttatttgtacaaaattcaaaaaagccatgagaaaaaaggaggtgcaactcattttat-3’) with 35 bp homology regions upstream and downstream of the gene, completely knocking out the ORF of the gene. The sequence of the ssODN allowed us no use no mutations to prevent accidental targeting by crRNAs.

#### Floxed *mec-4*

On the MSB943 strain (mScarlet::*mec-4*) we first introduced a loxP site in intron 1 using crRNA #104 (5’ acgtaattttcaaattgcaa 3’) and ssODN agaagtacaagtgagtttacgtaattttcaaattgATAACTTCGTATAGCATACATTATACGAAGTTATcaacggtaaattttatttttgtatgcaattgtatt as homology repair template (loxP site in UPPERCASE). Once verified by sequencing (MSB1509[*mec-4(mir50mir117)*]) we introduced the second loxP site with crRNA #105 (5’ CACTGAATCAGAGGCTTATG 3’) and repair template 5’gttgaattttgaaatgctcactgaatcagaggcttaCggagtgagtATAACTTCGTATAGCATACATTATACGAAGTTATtgaaaaatttaggcaaagaacgtgtatcaaagaaa 3’ (loxP site in UPPERCASE) with 1 point mutation (UPPERCASE C). We named the final allele *mec-4(mir50mir117mir118)* and generated strain MSB1514.

### 1.3 Locomotion experiments

#### 1.3.1 Cohort tracking

##### Imaging setup

Locomotion experiments were carried out in a home-built setup (Fig. 1 b), with a large format sCMOS camera (Imaging Source, DMK38UX255) using imaging source acquisition software and the objective with adjustable magnification (Fujifilm Fujinon, CF25Za-1S). Before the experiment, the plates were transferred to the imaging chamber and left to settle, for 30 min on NGM plates and 10 min on PAA gels. The recordings were carried out for 15 min on NGM plates and 10 min on PAA gels, unless otherwise stated, sampled at 20 Hz. The PAA gel recording was performed inside an additional isolated chamber with a water reservoir to avoid drying of the PAA gels. Brightfield illumination with white-light LED of intensity 3.7 *µW/mm*^2^ was used (measured in the worm plane) unless otherwise stated. In the tetracycline conditional rescue experiment, the dark-field red LED ring-shaped illumination with the intensity 1.85 *µW/mm*^2^. In the optogenetic experiments with *ACR-1*⇒ *TRNs*, dark field illumination light with the intensity of 0.15 *µW/mm*^2^ for green, 519 nm and 0.11 *µW/mm*^2^ for red, 519 nm was used.

##### PAA gels

PAA plates were prepared in different acrylamide / bisacrylamide ratios (as tabulated below), resulting in different stiffness moduli (*62*). The PAA gels were obtained by mixing the solutions of PBS, acrylamide (40%), bisacrylamide (2%), ammonium persulfate (APS) and tetramethylethylenediamine (TEMED), as listed in Tab. 2.3.1. 500*µL* of the mix was placed in the glass well plates, immediately covered with coverslip glass and left to polymerize for about an hour. The plates were stored at 4 °C dipped in PBS solution until use and used within a week. For the spontaneous locomotion assay, plates were seeded with 50 *µL* concentrated OP50 and left in the hood until dry.

| | Acrylamide<br>(40%) $\mu\text{L}$ | Bisacrylamide<br>(2%) $\mu\text{L}$ | PBS<br>$\mu\text{L}$ | APS<br>$\mu\text{L}$ | TEMED<br>$\mu\text{L}$ |
| --- | --- | --- | --- | --- | --- |
| 0.5 kPa | 50 | 7.5 | 437 | 2.5 | 0.25 |
| 1.25 kPa | 68.75 | 12.5 | 416 | 2.5 | 0.25 |
| 6 kPa | 93.75 | 15 | 388.5 | 2.5 | 0.25 |
| 30 kPa | 93.75 | 80 | 323.5 | 2.5 | 0.25 |
| 100 kPa | 150 | 110 | 227.25 | 2.5 | 0.25 |

##### Locomotion on tetracycline NGM plates

To block the self-cleavage of the tetracycline aptazyme, NGM agar containing different concentrations of tetracycline were prepared fresh before each experiment. The stock solution of tetracycline was distributed on the surface of the plates to reach the final concentrations of 2.5, 5 and 10 *µ*M. The animals were synchronized, seeded and left at 20°C incubator until reached late L4 stage. Then they were transferred to tetracycline plates and left for about 4.5-5 h. The locomotion experiments were performed as descroibed earlier. The recordings were carried on in earlier described setup for 15 min.

##### Data processing

The videos were processed with Tierpsy Tracker (*63*). The parameters were adjusted for each video; example parameters file can be found in supplementary material. The trajectories of the crossed worms were linked manually. The tracks were further processed with customised Python scripts, then the violin plots with average for each individual were plotted using seaborn package.

#### 1.3.2 Single animal imaging on PAA substrates

Single animal imaging and tracking was performed as previously described (*32*). Briefly, 3-5 min long videos of young adult synchronized animals transferred to non-seeded polyacrylamide gels (without bacteria as food source) were recorded at 25 fps using a home-built tracking platform. Imaging processing was performed using eigenworms with custom MATLAB scripts. To compare genotypes, the behavior of the locomotion of the animal was decomposed using the Eigenworm analysis (*31*). To construct and visualize the 3D distributions of the modes (a_1_, a_2_, a_3_), the 3D kernel density estimate of the first three modes was calculated in R using the `ks` package (*64*) with an unconstrained plug-in selector bandwidth. We choose to indicate the 10, 25 and 50% contours of the highest density regions in the manifold and the 2D projections of the floating data cloud along the corresponding planes. To compare two different data sets and test for the null hypothesis that the two kernel density functions are similar, the local density between two distributions was tested using a binned kernel density estimator. For this, we used the `kde.local.test` function from the `ks` package in **R** (*64*) to a) converted all n data points of each distribution to counts on a 100×100×100 binning grid and embedded into a matrix C, b) evaluate the kernel function at these grid points to embed them into a matrix K and c) then obtain the binned density estimator f from a sequence of discrete convolutions of C and K. The exact formulation of that procedure and a detailed presentation of the algorithm can be found in ref. (*65*). For all grid points in which p >0.01, we accept the null hypothesis that the two densities at this grid point for the two distributions are the same. For all other grid points, a polarity is assigned and plotted as two different colors in a 3D voxelgram (e.g. Fig. 1 & Extended Data Fig. 3), indicating that x_1_ > x_2_ or x_1_ < x_2_.

### 1.4 Calcium imaging in freely moving worm

#### 1.4.1 Imaging setup

Imaging was performed using an in-home built dual-camera upright epifluorescent microscope, with two Hamamatsu ORCA-Fusion cameras C14440-20UP, Lumencore LED Light source Spectra X, with cyan LED, 475/28nm, at a final light intensity of up to 47.5mW in the worm plane, and green LED, 555/28nm, at the final light intensity of up to 83mW at the worm plane, measured with a Thorlabs microscope slide power meterhead (S170C) attached to PM101A power meter console. The hardware was configured in master-slave configuration, with one camera functioning as a master, sending TTL pulse to the second camera and LED light source. This helped minimizing the phototoxicity. In the epifluorescence path, the dichroic filter AHF FF493/574-Di01-25×36 was used. Emitted fluorescence was then split into two cameras using dichroic beamsplitter AHF T 560 LPXR and filters ET510/20m and ET620/60nm (both Chroma, commercially purchased from AHF) for GCamp and TagRFP respectively. The stage Märzen-häuser Wetzlar SCANplus IM 130×85 S/No. 17120519 was used together with TANGO 2 Desktop controller Art. No.: 31-76-150-1802. Z-stage controller PI C-863.12 C-863 Mercury Servo Controller was used for vertical adjustments. The objective used for imaging Nikon S Plan Fluor 20x/0.7, air.

#### 1.4.2 Automatic worm tracking napari-based plugin Track-live

We implemented Track-live plugin in napari (*66*) using napari multithreading to 1) manage the communication between two cameras, 2) reading and manipulating the XY stage position to keep the worm within the field of view, 3) live display of the videos that were being acquired, and 4) allowing a user to interact with the interface (e.g. terminate the acquisition earlier, switch to manual XY tracking). LED and XYstage were controlled through Micromanager (*67*) and the former also through the Lumencor-CIA plugin, and were further passed through the Pycromanager (*68*) communication bridge to Track-live. The cameras were controlled through Python Hamamatsu SDK. After read-out from the camera the images were binned twice for further processing and analysis. The live image analysis, enabling XY stage adjustments to follow the worm, relied on finding the centroid position of the binary mask obtained by thresholding the blurred auto-fluorescence image from a ‘red-filter camera’. The threshold was set to the 90th percentile value from the image when the user pressed the tracking button, ensuring that the worm is within the field of view. Tracking in XY plane was occasionally prone to detect artifacts, therefore it was at times switched to manual. Z adjustments were done manually. Exposure time was set to 20 ms, ensuring the reasonable trade-off between image quality and blurring as the worm moves. The videos were acquired for 2 min, at a sampling frequency 8 fps. The videos were stored in the program memory, and saved to drive after the acquisition was finished. The code is available at https://gitlab.icfo.net/NMSB/track-live-close-loop

#### 1.4.3 Video processing

##### Neural tracking & neural activity

The cell bodies of the TRNs were identified within each frame and their trajectories were linked using a simple LAP tracker implemented in the TrackMate Fiji plugin (*69*). The trajectories were then manually curated and cleaned. Each neuron was then manually assigned to one of the class AVM, ALM, PVM or PLM based on the location. In order to approximate neural activity, circular areas around the cell body (with radius 10 pix, which corresponds to 6.5*µm*) were cropped in each frame, averaged over the ROIs for the green and red channels separately, F*_ROI_*. For each channel, background fluorescence was defined for each frame as the average of all pixels below the 20th percentile F*_bckg_*, while worm fluorescence was defined as the average of all pixels between the 50th and 70th percentiles F*_worm_*. The fluorescence intensity was estimated as:

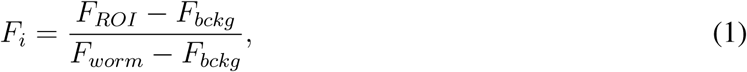

for both channels, *i* ∈ {*GCamp, T agRFP*}. The neural activity was then approximated as the ratio of the ‘green’ and ‘red’ fluorescence values *r* = *F_GCamp_/F_T agRF P_*.

##### Skeletonisation

In the first step, each frame from the red channel was denoised using `fastNlMeansDenoising` function form Open-cv Python package, and then thresholded using `cv2.threshold` and manually selected optimal threshold. The binary masks obtained were then dilated using `cv2.dilate` function to fill in the holes in the binary mask. Then all contours found with `cv2.findContours` function, with the area that reached 10.000 pixels or more, were kept as potential worm candidates and otherwise ignored. The edges of the masks were smoothed using the sequence of functions `cv2.GaussianBlur`, `skimage.exposure.rescale_intensity`, and then `cv2.threshold`. The binary masks were then skeletonized `cv2.ximgproc.thinning`, fil_finder pipeline, which included `preprocess_image, create_mask, medskel, analyze_skeletons` functions. The skeleton was then identified as the longest path found and interpolated to 50 points using cubic spline interpolation. Frames, where the skeletons were shorter than 750 pixels (∼ 490*µm*) or longer than 130% of the average, were considered unreliable and removed from further analysis. Skeleton finder is not sensitive to worm orientation, which can lead to systematic flips in the head-tail orientations. In order to account for that, the skeletons were then oriented from head to tail by minimizing the Cartesian distances head-head and tail-tail between the consecutive frames, and by using the neuron positions within the body, e.g. PLM is located in the tail, while ALM/AVMs are located in the anterior body.

#### 1.4.4 Time-series analysis

For each worm, each frame and for each neuron, tangential and normal directions were found using the skeleton point closest to the cell body. The velocity was then decomposed into tangential and normal components, *v_t_* and *v_n_*, respectively. In further analysis 5-second windows were considered. For each of them, the cross-covariance and cross-covariance coefficient between the ratio *r* and the tangential velocity *v_t_* was estimated using `scipy.signal.correlate`, mean-subtracted signals, and ‘valid’ modality of the cross-covariance function `scipy.signals.correlate(r, v, mode=’valid’)`, where r (1st argument) is shorter, 5 sec and v (2nd argument) is longer 10 sec and their middle points corresponds to the same point in time. The maxima of the cross-covariance, including both the peak value and its corresponding time lag *τ*, were grouped and smoothed using a kernel density estimator (`seaborn.kdeplot`). Per convention, we assigned positive time lags when the velocity precedes TRN activity; and negative lags when the TRN activity precedes reversals or increases in velocity (see also Supplementary Fig. 4). Next, we statistically verified whether the peaks originate from concurrent activity in both time series of calcium and velocity, or occur rather as inner properties of each time series. For details see Section 2.8.2 and Extended Data Fig. 6. The codes for sections 2.4.3 - 2.4.4 are available under the link: https://gitlab.icfo.net/NMSB/cal-free-m-worm

### 1.5 Optogenetics

#### 1.5.1 TRNs killing using PH-miniSOG

In order to obtain acute killing of the touch cells, a strain expressing mini Singlet Oxygen Generator in the TRNs was used, CZ23281 (*70*). MiniSOG generates the reactive oxygen species (ROS) singlet oxygen after illumination with blue light. Illumination of neurons expressing miniSOG targeted to the membrane (PH-miniSOG) causes neuronal death. The extrachromosomal worms were exposed to blue light 470 nm of the intensity 1.7 mW/mm^2^ (Mightex, Series Light Source Control Module) for 4 min around 18 hours before locomotion experiment. The light was mounted on specially designed box, 10 cm above the surface of the plate that ensured the uniform illumination of the whole plate. After the illumination the worms were kept in the darkness until the locomotion experiment.

#### 1.5.2 Stimulation of the touch receptor neurons with ChR2

In order to investigate if the lethargic phenotype can be rescued by chronic stimulation of the Touch Receptor Neurons we crossed strains AQ2334 and GN693 to generate MSB990, expressing channel-rhodopsin (ChR2(H134R)) under the control of the*mec-4p* promoter in the *mec-4(u253)* mutant background. L1 worms were seeded on plates with 100 µL of OP50 mixed with ATR to final concentration of 0.1 mM prepared as described in (*71*). The animals were then stimulated with 455nm LED (Osram Oslon power cluster; ILR-ON16-DEBL-SC211-WIR200) light powered by R&S NGE100B Power supply series. The power supply was controlled by customised python code, to generate the poisson-like light pattern with average interval between pulses of 5 min, each pulse lasted 3 seconds. The worms were stimulated for about 50h, transferred to plates without ATR and assayed in the locomotion setup using dark field ring-shaped red light illumination (as described earlier) about 2h later.

#### 1.5.3 Silencing of the touch receptor neurons with ACR1

MSB1244 worms expressing *mec-4*p::cGAL4 and UAS::ACR1 were obtained by crossing the previously described MSB952 strain (*61*) to MSB927 (*mirEX238*[*mec-4*p::cGAL4+*myo-2*p:mCherry]). Plates with ATR were prepared as specified in the previous section and MSB1244 L4 worms were seeded on them. After 16h incubation at 20°C animal were recorded for 15 min and analysed as described.

### 1.6 Calcium imaging in the microfluidic body wall touch device

#### Preparation of the mold using dry etching

The mold was fabricated by Reactive Ion Etching (RIE) on a silicon master wafer. Deep RIE using the Bosch process is a microfabrication technique that allows etching silicon substrates and obtain high aspect ratio features such as the one used in this work, where microchannels of around 50 µm height were separated by actuators of 10 µm, which required narrow deep trenches with an aspect ratio of 5:1 in the mould. This method consisted in alternating steps of etching and passivation that provided highly selective etching and resulted in well-defined and vertical walls.

Before etching, the desired design was patterned on 4-inch silicon wafer (525 ± 25 µm, MicroChemicals, GmbH) as a resist mask by conventional photolithography. Briefly, a 22 µm layer of negative resin AZ-125NXT (AZ Electronic Materials GmbH) was spinned on the wafer at 3000 rpm for 20 s (SPIN150x Spin coater, Polos-SPS Europe B.V.) and baked for 15 min at 125 °C. Then, the body-touch design was exposed using a maskless aligner (MLA150, Heidelberg Instruments) with a UV-light (375 nm wavelength) dose of 900 mJ/cm^2^, developed by stirring in AZ 726 MIF developer (AZ Electronic Materials GmbH) for 8 min, and rinsed in DI water. Using this mask to protect the silicon below, the Bosch process was performed in a RIE system with Inductively Coupled Plasma (SI 500 ICP-RIE Plasma System, Sentech Instruments GmbH). For the etching steps, SF6 was used at an ICP power of 950 W, an etching RF power of 25 W and a gas flow of 129.9 sccm, with an etching time of 3 s. For the passivation steps, C4F8 was used at an ICP power of 800 W, a passivation RF power of 5 W and a gas flow of 80 sccm, with a passivation time of 2 s. The substrate was kept a constant temperature of 10 °C. These steps were repeated for 200 cycles to achieve a total depth (resin and etched silicon) of around 50 µm.

#### Device preparation

The devices were replica-molded as previously described (*8*). In short, the devices were prepared by casting a mix of PDMS base and curing agent (Sylgard™ 184, Dow) in a 10:1 proportion on the mould and baking it for 2 h at 70 °C. To facilitate pealing the PDMS, the wafer was previously silanized with chlorotrimethylsilane (Merck KGaA) for 30 min. After carefully pealing the PDMS, the individual devices were cut and 0.75 mm inlet and outlet perforations were created by a pneumatic puncher. Finally, the patterned PDMS and glass coverslides (#1.5, VWR) were irreversibly bonded by oxygen plasma activation. Diaphragm deflection versus pressure was previously calibrated for both mixtures (*72*).

#### Animal loading into a microfluidic trap

Loading of the animals in the body wall chip was performed as described in detail elsewhere (*73*). For imaging PDE in control and *mec-4*^-/-^ conditions, we used MSB661 and MSB636, respectively. For imaging DVA neuronal activity secondary to direct mechanical stimulation of the body wall, we used MSB273 and MSB1410 in the control and *mec-4* mutant background, respectively. Briefly, 2-3 young adult animals were transferred to a M9-filtered droplet. The worms were then aspirated through an SC23/8 gauge metal tube (Phymep) connected to a 3 mL syringe (Henke Sass Wolf) with a PE tube (McMaster-Carr). Once the tube was inserted into the chip inlet, the animals were loaded into the waiting chamber by applying gentle pressure with the syringe.

#### Calcium imaging and mechanical stimulation

*In vivo* calcium imaging of TRNs was performed by positioning the worm-loaded microfluidic device in a Leica DMi8 microscope with a 40x/1.1 water immersion lens, Lumencor Spectra X LED light source, fluorescence cube with beam splitter (Semrock Quadband FF409/493/573/652), a Hamamatsu Orca Flash 4 V3 sCMOS camera and HCImage software (version 4.4.2.7). Cyan-488 nm (≈6.9 mW) and yellow-575 nm (≈12.6 mW) illuminations were used to excite the green and red fluorescence of TRN::GCaMP6s and mtagRFP, which was used to correct possible artifacts of animal movement and identification of TRN. The incident power of the excitation light was measured with a Thorlabs microscope slide power meter head (S170C) attached to the PM101A power meter console. Emission was split with a Hamamatsu Gemini W-View with a 538 nm edge dichroic (Semrock, FF528-FDi1-25-36) and collected through two single band emission filters, 512/525 nm for GCaMP6s (Semrock, FF01-512/23-25) and 620/60 for mKate (Chroma, ET620/60m). The emission spectra were split by the image splitter, allowing different exposure times for each signal. For the application of mechanical loads to the animal’s body wall, the stimulation channel was connected to a piezoelectric pressure controller (OB1-MK3, Elveflow) as described (*71*). To follow calcium transients, videos were taken at 10 frames-per-second with 80 ms exposure time, using the master pulse of the camera. The camera SMA trigger out was used to synchronize the stimulation protocol in Elveflow sequencer, which consisted of three stimulation, 20 s pre-stimulation, 2 s stimulation (2500 mbar buzz). Each stimulus was separated by 20 s.

#### Calcium analysis

Images were processed using ImageJ and intensity signals were extracted as described (*71*). First, the neuron of interest was manually labeled on the basis of the calcium insensitive channel mtagRFP. The position was automatically tracked in the following frames and used to extract the GCaMP intensity. A smooth filter (moving average filter) was applied. The calcium sensitive signal was background subtracted and then normalized to the first 100 frames pre-stimulation ((*F* − *F*_0_)*/F*_0_). After arithmetic averaging all recordings, the *t*-test statistics for each timepoint in the timeseries was calculated according to 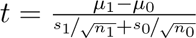 where *µ*_1_ and *µ*_0_ are the mean values of each group at time *t*, *s*_1_ and s_0_ are the standard deviations for each group at time t and n equals the number of recording for each group. The t-test statistics was then used to calculate the p-value considering the degrees of freedom *n*_1_ + *n*_0_ − 2 and the null hypothesis was rejected if the absolute value of *t* was larger than its critical value at the level of significance *α*=0.01. All operations were performed in R using custom routines.

### 1.7 Traction Force Microscopy

#### 1.7.1 Polyacrylamide gels preparation

In order to simplify the imaging, reduce the potential artifacts, and ensure the estimation of the tractions on the surface of the gels, we assured placement of the fluorescent beads on the surface of polyacrylamide (PAA) gels. The gels were prepared following the protocol from (*74*), unless stated otherwise. In short, the cover-slip glasses were functionalized with poly-L-lysine, air-dried, coated with fluoSpheres 0.2*µm* carboxylated beads (Fisher, F8809), with spectral properties of 540/560 diluted with miliQ H_2_O in a ratio of 1:2000, and sonicated for 30 min. Polyacrylamide gels were prepared in three different concentration ratios of bisacrylamide and acrylamide, as described above. The spectral properties of the beads were chosen in order to minimize the effect of the excitation light on the worm’s behavior.

#### 1.7.2 Experimental setup

The image processing pipeline has been developed in a way that allows the use of various setups, without the need for a dual view camera, as long as segmentation of the worm silhouette is possible from the videos. The simultaneous stage position acquisition is not necessary, which is a caveat when using most of commercially available image acquisition software, and making a manual worm following a feasible option. The stage position movement is estimated from the background image matching between the consecutive frames. For best information content, each frame should include a full image of a worm and a sufficiently dense bead image underneath.

#### 1.7.3 Image preprocessing

Each frame was divided into three parts, 1) background ‘far away’ from the worm, where the beads image is unaffected by traction forces exerted by the worm, which can be used to estimate offset stage movement; 2) worm silhouette that can be used to find the behavioral characteristics of the worm, such as body curvature; and finally, 3) background region in the close vicinity of the worm empirically chosen to be around 1.5 worm width, where the beads image is affected by the traction forces exerted by moving worm.

#### 1.7.4 Worm segmentation

The worm videos were first segmented to find the worm silhouette, using tracker annotator from ‘Segment Anything for Microscopy’ napari plugin (*75*) using the ViT-L model. The parameters for each video were chosen individually, based on the movement of the worm, to maximize the automatic segmentation of the worm. The incorrectly segmented frames were further manually curated and the worm images were kept for further analysis. The worm masks were further processed as described in Section 2.4.3.

#### 1.7.5 Offset correction

‘Far-away’ background regions were further used to estimate offset stage movement in the pairs of the consecutive frames using subpixel maxima in cross-correlation landscapes, implemented in openpiv packages and self-prepared scripts.

#### 1.7.6 Particle image velocimetry

The pairs of the consecutive frames, containing regions of the image in the close vicinity of the worm, were used to estimate particle image velocimetry, using the openpiv package (*76*). The parameters chosen were 42 pixels of window size, with 43 search windows, and 60% overlap, and a pixel size of 0.65 *µm*. The PIV in the worm and in the background regions was set to zero, potentially irrelevant for the analysis.

#### 1.7.7 Traction estimation

The traction forces were estimated using the pyTFM package (*77*), which uses the Fourier transform to estimate the traction fields (*78*). The parameters used: Poisson ratio of the substrate, *σ* = 0.457, height of the substrate, *h* = 1*mm* and Young’s modulus of the substrate 1.25, 6 or 30 kPa.

#### 1.7.8 Traction quantification and behavioral analysis

The profiles of the traction fields along the body of the worm were further estimated, using geometrical image transformations: that is, the images of the traction fields were straightened along the skeletons. The images were processed using self-prepared functions and map coordinates from the scipy.ndimage package. The profiles were obtained separately for magnitudes, tangential and normal components with respect to skeleton midlines. For each profile, for each body point, the 97.5 percentile of the absolute value of the array was selected, and later was assigned back its sign, in order to preserve the negative values. For the time point and for each midline of the skeleton, the body angle was found and normalized between −*π* and *π*.

The codes described in sections 2.7.3 - 2.7.8 can be found under the link: https://gitlab.icfo.net/NMSB/traction-force-worm

#### 1.7.9 Statistical Analysis of the traction profiles

To account for repeated measurements along the body of each worm, tractions were modeled using a linear mixed-effects model implemented with the `lme4` package in R. Traction was the dependent variable, with *strain* and *body.position* (normalized from head to tail) included as fixed effects, along with their interaction (*strain* × *body.position*) to capture region-specific differences between strains. Random intercepts were included for each worm to account for multiple body points measured per individual. The model can be expressed as:

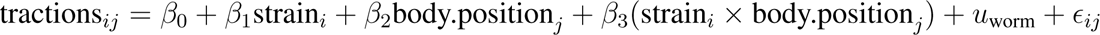

where 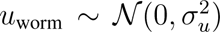 represents worm-specific random effects and *ɛ_ij_* ∼ *N* (0*, σ*^2^) is the residual error. Statistical significance of fixed effects was assessed using Type III ANOVA. Denominator degrees of freedom were estimated using Satterthwaite’s approximation to account for uncertainty in variance component estimation within the mixed-effects framework.. Post-hoc comparisons along the body axis were performed using estimated marginal means with multiple-testing correction where appropriate.

### 1.8 Statistics and reproducibility

Statistical modeling and hypothesis testing were performed in MATLAB and R (v. 4.2.2). No statistical methods were used to predetermine the sample sizes, but our sample sizes are similar to those reported in previous publications. Data distributions were assumed to be normal, unless obviously not, but this was not formally tested. All datasets were acquired in a randomized fashion, and when they were not, the data were not biased by the history. Some data were not collected blind to the genotype, and analyzed blind to the experimental condition (Fig. 1).

#### 1.8.1 Multilevel analysis

For multilevel hierarchical analysis, data did not meet the assumptions of normality, (Fig. 1), as indicated by the results of the Shapiro-Wilk test. Consequently, a mixed-effect linear model (*79*) was employed as an alternative to the adjusted t-test, for hierarchical data structures, where indicated. The average from individual worms as a first level, and different plates as a second level. The computations were performed using `statsmodel` Python module. A linear mixed-effects model can be expressed as:

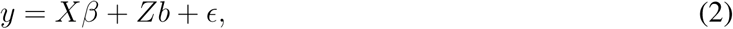

where *y* is the observed data. In our case, y is the statistic estimated for each individual worm. *Xβ* term accounts for fixed effects, *β*_0_ corresponds to baseline value in the control group, while *β*_1_ accounts for the difference between the groups. *X* contains information about the experimental conditions at the highest level, (e.g. group affiliation of each worm). *Zb* term accounts for random effects, *b* = *b_x_, x* ∈ {*plate*_1_*,…, plate_n_*}, and *Z* contains information about the experimental conditions at the lower level, (e.g. plate affiliation of each worm). *ɛ* corresponds to residual error. *β*, *b*, plate-level variability, individual worm variability are determined by maximizing the restricted likelihood. *β*_1_ stores the information about the difference between group means. *P*-value for the fixed effect is then obtained by comparing *β*_1_ to its standard error and using *t*-test. *t*-test assesses whether the group difference, *β*_1_ significantly deviates from zero.

##### Touch response

The analysis was performed in R using the lme4 package. Binary “yes/no” responses to mechanical touch were analyzed using a mixed-effects logistic regression to account for repeated trials within individual animals, with tetracycline dose as a fixed effect and individual animals as random intercepts to account for repeated touches on the same subject. Let *Y_ij_* denote the binary response (0/1) for trial *j* from animal *i*, with response probability *p_ij_* = Pr(*Y_ij_* = 1). We assumed

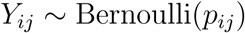

and modeled the log-odds of response using a logit link function:

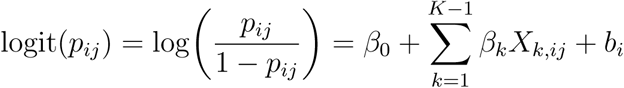

where *β*_0_ is the fixed intercept, *X_k,ij_* are indicator variables encoding the experimental conditions (with one condition serving as the reference level), and *β_k_* are the corresponding fixed-effect coefficients. The term *b_i_* represents a random intercept for animal *i*, accounting for within-animal correlation across repeated trials.

Random effects were assumed to be normally distributed:

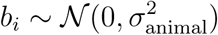

where *b*_0_*_i_* represents animal-specific baseline variation and b*_ki_* represents animal-specific deviations in the effect of condition. Random effects were assumed to be normally distributed.

#### 1.8.2 Surrogate testing

To statistically validate the cross-covariance between neuronal activity and velocity, we applied intersubject surrogate testing alongside a control group (*80, 81*). Specifically, to differentiate genuine concurrent activity (*H*1) from coincidental cross-covariance peaks arising due to the intrinsic properties of each time series (*H*0), we computed the cross-covariance between the neuronal activity of one worm and the velocity of another worm within the same group. This process was repeated for all possible pairings. The 2.5th and 97.5th percentiles of the surrogate cross-covariance peak values were used as reference thresholds corresponding to a *p*-value of 0.05. If the observed cross-covariance exceeded the 97.5th percentile, *H*0 could be rejected at the *α*=0.05 significance level, indicating that the peaks likely result from genuine concurrent activity in calcium dynamics and tangential velocity.

#### 1.8.3 Calculation of the effect size

To quantify the effect of the genetic and pharmocological perturbation on animal locomotion, we resorted to estimation statistics of non-normally distributed data. Cliff’s delta (*δ*) is a non-parametric measure of effect size that quantifies the degree of difference between two independent distributions. It is defined as the probability that a randomly chosen observation from one group is larger than a randomly chosen observation from the other group, minus the reverse probability:

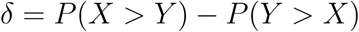

where X and Y are values from the two groups.

For every pair of observations (x*_i_*, y*_j_*), assign:

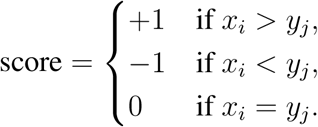

All scores were summed across all pairs and divided by the total number of comparisons (*n_X_*× *n_Y_*) to obtain *δ*:

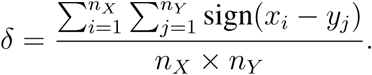

The value of *δ* ranges from −1 (all values in group *Y* are higher than *X*) to +1 (all values in group *X* are higher than *Y*), with 0 indicating no difference. Calculations were done using the `dabestr` library in R (*82*).

##### Inference

Bootstrapped 95% bias-corrected and accelerated (BCa) confidence intervals were computed from 5000 resamples of the data to quantify the uncertainty in *δ*.

## 2 Supplementary Discussion

### 2.1 Mathematical modeling of neural network and behavioral proxy

#### 2.1.1 Formulation

The model is based on an earlier work (*29, 30, 83*). The realistic *C. elegans* synaptic and gap junction connectome is used in conjunction with a single compartment neural model to simulate the dynamics of the network. Following (*29, 30, 83*), neurons were modeled with an isopotential single compartment membrane model, where the membrane potential of the i-th neuron V*_i_* is given by the equation:

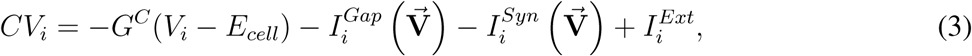

where *C* is the whole-cell membrane capacitance, *G_C_* is the membrane leak conductance and *E_cell_* is the leakage potential. The external input current is given by 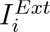, while neural interactions via gap junctions and synapses are modeled by input currents 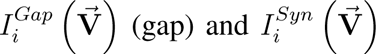 (synaptic), described by following equations:

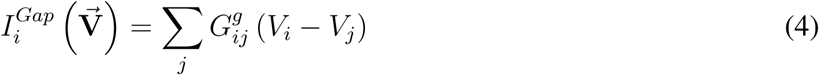

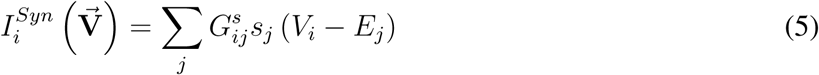

Gap junctions are taken as ohmic resistances connecting each neuron, where 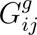 is the total conductivity of the gap junctions between i and j. The synaptic current is modeled as a diffusive coupling and depends on the displacement from the reversal potentials 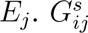 is the maximum total conductivity of synapses to *i* from *j*, modulated by the synaptic activity variable *s_i_*, whose temporal evolution is governed by:

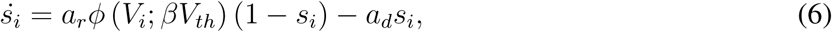

where *a_r_* and *a_d_* correspond to synaptic activity rise and decay time, and *ϕ* is sigmoid function, given by:

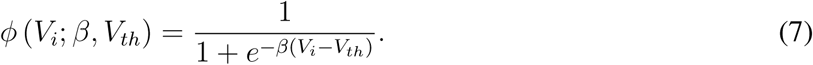

The parameters are selected as in (*29*), unless started otherwise.

#### 2.1.2 Network driver selection

We have tested several neuronal candidates receiving the constant current for drivers of the network, namely: AVB (as it delivers command signals to B-type motor neurons and it is known to drive forward locomotion, (*35*)), PLM (as it mediates the gentle touch response, triggers forward escape reflex and it was originally proposed as a network driver for this model (*29*)), AVF (as it was already reported as a persistent neuronal oscillator (*34*)). To find the feasible input current amplitudes that generate the oscillatory activity in the network Jacobian eigenspectrum was estimated using Matlab for various amplitude ranges. The values of input current amplitudes, for which at least one pair of the real part of eigenvalues was changing its sign, were selected to be used later as drivers. The oscillations in the network were the most stable when PLMR/L were used a network drivers. Therefore the following simulations were performed using PLMR/L neurons pair as a network driver.

#### 2.1.3 Estimating the oscillatory modes representing forward locomotion

The constant input current applied to PLM neurons, 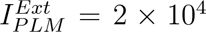, results in the emergence of the oscillations in the whole neural network, as visualized in Fig 2c. Following the procedure originally proposed in (*29*), the singular value decomposition is then applied to B & D type motor neurons activity, which reveals an existence of the dominant oscillatory modes representing forward locomotion (*31*), indicated as a_1_ & a_2_ on Fig. 2d, and allows to find the weights corresponding to the contribution of each motor neurons in the locomotion modes.

#### 2.1.4 Mechanosensitive current implementation

The model was originally used to computationally investigate the effect of injury or ablations on network dynamics. In this research, it was applied to investigate the effect of ‘removing’ the external input that represents the mechanosensitive current from the TRNs. An important assumption to mention is that the synapses were imposed half open at equilibrium membrane potentials (mean) (where *ϕ*(*V_eq_*) = 0.5, *V̇_i_* = 0, *ṡ* = 0), following (*29, 84*). This can be achieved by adjusting *V_th_* in Eq.7. In the first part of the simulation, 0-15 s, representing an intact worm receiving the mechanosensitive current, in addition to applying the PLM driving current, one of the TRNs (AVM, ALM or PVM) received a small constant input current with an amplitude varying between 0.5 and 5% of the driving current amplitude. The average membrane potential and thus the synaptic equilibrium, as well as the neuronal weights to each singular value, were established taking into account both the driving current and an additional ‘mechanosensitive’ input applied to one of the TRNs. In the second part of the simulation (20 - 30 s) this additional current was removed, which mimicked the loss of the mechanosensitive current. This resulted in an increase in the amplitude of the oscillations in the motor neuron membrane potential, leading to an increase of the body curvature, or a total extinction of these oscillations, resulting in the lack of motion, or in other words lethargus. This suggested that both increase in body curvature and lethargus, seemingly very different phenotypes, can have a common underlying origin in loss of mechanosensation in TRN. In the third part of the simulation (30-25 s), synaptic equilibrium values were adjusted to new values, by modifying *V_th_* in Eq. 7 and taking into account driving current only (but no mechanosensitive currents), then the synapses were imposed half-open at new equilibrium values. This part represents adapted *mec* mutants.

#### 2.1.5 Parameters

All the parameters are taken from the original paper (*29*), except where stated otherwise. Individual gap junction and synapse was assumed to have approximately the same conductance, roughly *g* = 100*pS*. Each cell has a smaller membrane conductance (taken as 10*pS*) and a membrane capacitance of about *C_i_* = 1*pF* (*83*). Leakage potentials are all taken as *E_c_* = −35*mV* (*84*). Reversal potentials *E_j_* are 0*mV* for excitatory synapses and −45*mV* for inhibitory synapses. For the synaptic variable, we choose *a_r_* = 1, *a_d_* = 5 and define the width of the sigmoid by *β* = 0.125*mV* ^−1^. *V_th_* is found by imposing that the synaptic activation *ϕ* = 1*/*2 at equilibrium (*84*).

The simulation codes are available: https://gitlab.icfo.net/NSMB/net-sim-worm

### 2.2 Details of the worm locomotion model

Here we resort to the geometrically exact beam formulation in statics (*85*) and two dimensions, extended with active bending torques m*_ac_*(s, t) at midline coordinate s and time t, and a wet friction distribution, in agreement with resistive force theory (*43, 86, 87*). The latter is modeled as a distributed force on the beam proportional to velocity ***ẋ***, and expressed as −***ηẋ***, with 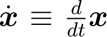 the velocity of the midline position ***x***, and ***η*** a frictional tensor given by (*88*)

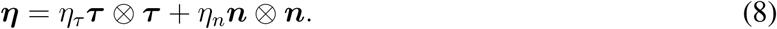

Parameters *η_τ_* and *η_n_* correspond to tangential and normal frictional coefficients, and ***τ*** and ***n*** the tangential and normal vectors at each point of the worm midline *s* ∈ [0*, L*], with *L* the length of the worm (see Figure 3.2). Consequently, the beam equations read

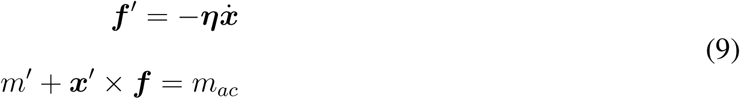

where 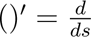, and vector ***f*** = **R*F*** denotes the axial forces at each cross-section, while *m*(*s, t*) are the elastic bending moments. Rotation matrix 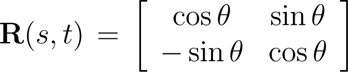 defines the orientation of the cross-section, with *θ*(*s, t*) the angle of the rotation, as shown in Figure 3.2.

Note that in view of the friction tensor in (8), the external forces at the rhs of the first equation of (9) are equal to −***ηẋ*** = −*η_t_**v**_τ_* − *η_n_**v**_n_*, with ***v****_τ_*= (***ẋ***· ***τ***)***τ*** and ***v****_n_*= (***ẋ***· ***n***)***n*** the tangential and normal components of the velocity vector ***v*** ≡ ***xẋ***, respectively.

We assume an active elastic body of the worm, that is, the unrotated axial forces ***F*** and moment *m* are assumed to obey linear elastic constitutive laws, conjugate to strain measures **R***^T^ **x***^′^ − **E**_1_ and *θ*^′^,

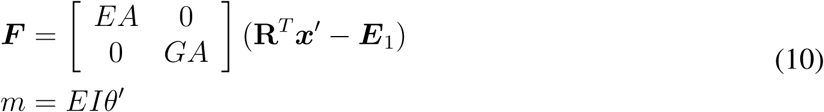

Constants *E, G, A, I* are material and geometrical parameters corresponding to Young modulus, shear modulus, cross-section area and moment of inertia of the worm.

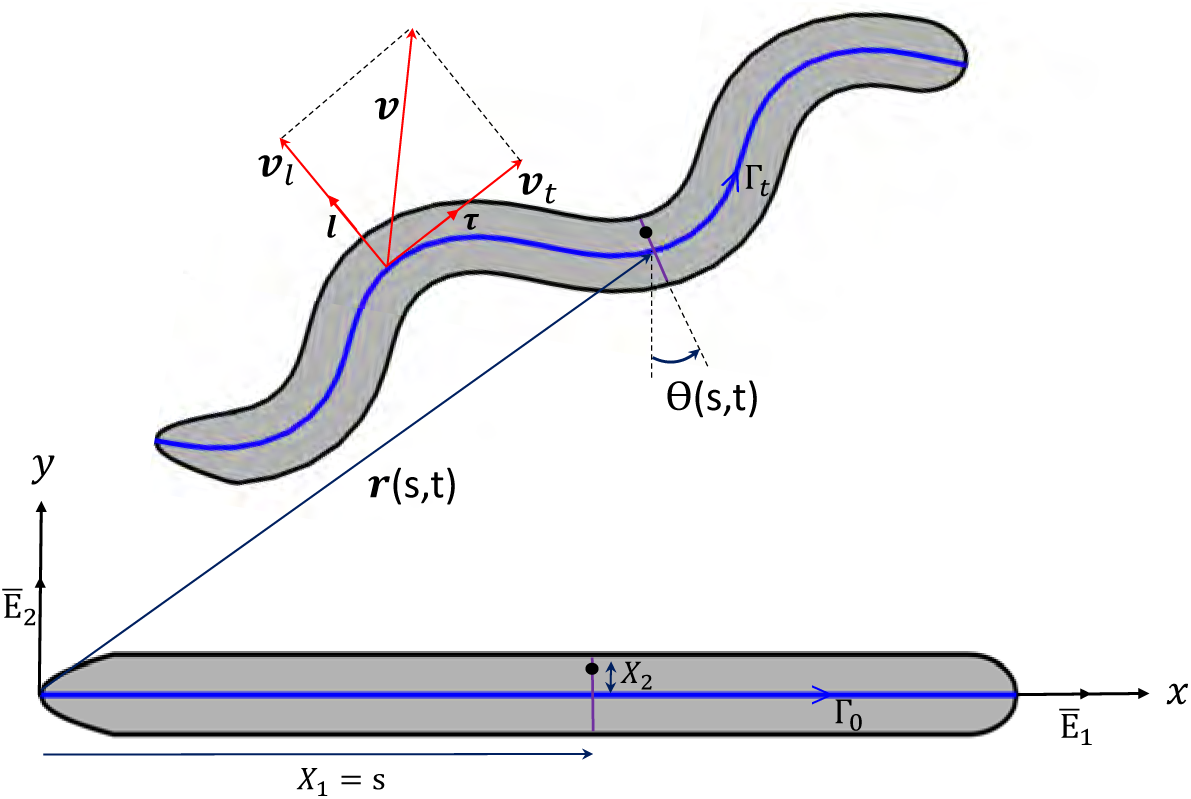
Scheme of slender body kinematics, indicating normal vector ***v****_n_* and tangential component ***v****_τ_* (*s, t*), with ***v*** = ***ṙ*** the velocity vector.

The direct problem of the worm consists of, given a time-history of spatial distributions of viscous parameters *η_t_τ*(*s, t*) and *η_n_*(*s, t*), and a torque distribution *m_ac_*(*s, t*), finding the sequence of positions ***x***(*s, t*) and angles *θ*(*s, t*), for an initial configuration ***x***_0_(*s*) = ***x***(*s,* 0) and *θ*_0_(*s*) = *θ*(*s,* 0).

We solve this direct problem resorting to a Finite Element (FE) discretisation of unknowns ***x*** and *θ* from a set of nodal values ***y****_i_* = {***x****_i_, θ_i_*}, *i* = 1*,…, N*. The FE method allows us to write the partial differential equations (PDEs) in (9), together with the constitutive law in (10), as a set of ordinary differential equations (ODEs), which may be written in a compact manner as,

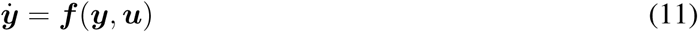

where ***y*** = {***y***_1_*,…,* ***y****_N_* } denotes the set of nodal general coordinates, and vector ***u*** = {***u***_1_*,…,* ***u****_N_* } a set of nodal triplets ***u****_i_* = {*η_τ_, η_n_, m_ac_*}*_i_* containing the values of friction and torques at node *i*. We note that for writing the static beam equations in the form (11), we include a small wet friction contribution also to the rotations, that is, we add a term −*η_m_θ̇* to the rhs of the second equation in (9).

### 2.3 Inverse analysis through Optimal Control Theory

We consider the following inverse problem: given a time-history of worm kinematics (i.e., nodal positions and rotations ***y****_exp_*(*t*)), deduce the set of torques and frictional parameters ***u****_i_* that best agree with the measured values ***y****_i_* while satisfying beam ODEs in (11). Since the rotations *θ_i_* cannot be experimentally measured, we assume that each *θ_i_* corresponds to the orientation of the tangent of the midline, that is, we assume no shear deformations.

While direct problems are in general well-posed and have a unique solution, inverse problems may have multiple solutions and be very sensitive to initial data. For this reason, we adapt the formulation of optimal control problems (*89,90*) to i) compute the optimal deformations for the locomotion of the worm (a new set of ***y****_i_* that approximates ***y****_exp_*), and ii) deduce muscle activity from experimental measurement (best matching values of ***u****_i_*).

This inverse problem is formulated as the solution of the following optimization problem (*44, 91*):

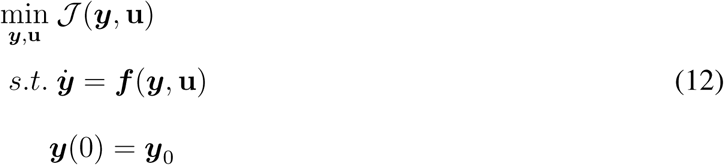

with ***y***_0_ the initial nodal positions and angles of the midline, and J (***y***, **u**) the functional

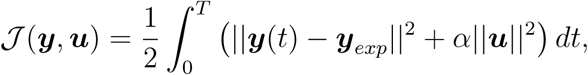

which aims to minimize the distance from the calculated position to ***y****_exp_*, while also minimizing the cost ***u*** through the weight parameter *α*. The second (regularization) term also aims to furnish unique solution to the optimal control problem (OCP) in (12). This OCP can be solved by adding the adjoint variable ***λ***, which also reveals a symmetric structure and form,

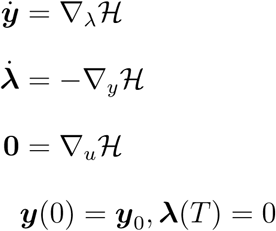

with 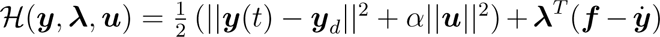 the Hamiltonian of the OCP. The equations above form a two-point Hamiltonian boundary value problem that we will solve resorting to a Forward-Backward Sweep Method (*89, 92, 93*).

In our optimization problem, the control variable **u** includes the active moments *m_ac_* in eqn. (9), and the unknown friction coefficients *η_t_* and *η_n_*. We note that the three quantities depend on space and time, and are defined at each node of the FE discretisation, except *m_ac_* which is equal to 0 at the end nodes.

The resulting inferred values in **u**, and optimal positions *mathbfy* that best match the experimental coordinates ***y****_exp_* allow us to compute the total dissipated and bending energy of the worm along the time span [0, *T*] as,

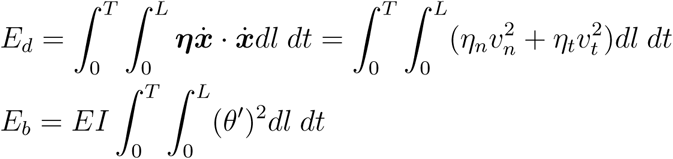

where v*_t_* = ||***v****_t_*|| and v*_n_* = ||***v****_n_*|| are the norms of the normal and tangential velocity components. We have taken the ratio *E_d_*/*E_b_* as a measure of lost energy during locomotion, which as shown in Fig. 6k is higher fo the *mec-4* mutant.

### 2.4 Notes on the time-series analysis of calcium activity and tangential locomotion

Since the seminal contribution of the Chalfie lab in determining the role of individual touch receptor neurons to the mechano-escape reflex (*9, 35*), the consensus in the field is that PVM does not contribute to the behavioral pattern generation, despite being mechanosensitive in itself (*8, 33*). Behavioral experiments carried out on animals with individually laser-ablated neurons have revealed that anterior neurons ALML/R and AVM are mediating a reversal escape response when animals are stimulated on the anterior body segments, PLML/R have been shown to be crucial for acceleration when animals are stimulated on the posterior body part. In contrast, killing PVM has little consequences on touch-evoked behavior, despite PVM being mechanoresponse to body touch (*8,33*). In particular, from the work presented in reference (*35*) paper we learned: *“Ablation of PVM has no detectable effect upon touch sensitivity. Killing all the microtubule cells, or all except PVM, results in animals that are touch insensitive over their entire length, except at the very front of the head.”* Thus, functional significance for PVM in the touch reflex has yet to be demonstrated, and remains unknown (*94, 95*). In addition, a recent investigation aiming to decipher the role of individual TRNs on behavior suggested that PVM has a modulatory, rather than a driving role (*36*).

In our experiments which address the role of TRNs in locomotion behavior, we observed that the calcium dynamics showed a complex and context dependent behavior.

The plots in Figure 4 and Supplementary Figure 4, we display the distribution of cross-covariance values between calcium activity in the sensory neuron and locomotion velocity across lags (*τ*). Cross-covariance was computed for each recording trace, and the distributions for control and *mec-4* animals are shown as density maps (center) with their respective marginal distributions projected onto the top and right axes. Importantly, cross-covariance preserves the absolute magnitude of joint fluctuations and is therefore sensitive to changes in signal amplitude, making it well suited for detecting causal structure and asymmetric coupling. In contrast, cross-correlation normalizes the signals and highlights only relative similarity, which can obscure amplitude-dependent effects. Because our goal was to capture how variations in locomotor dynamics scale with, and potentially activate, changes in TRN activity, cross-covariance provided a more informative measure than correlation.

Lags are defined such that positive *τ* values indicate velocity leading calcium, and correspond to sensory encoding, where changes in velocity precede and are followed by changes in neuronal calcium. In contrast, negative *τ* values indicate calcium leading velocity and the left quadrants reflect motor command generation, where neuronal calcium elevations precede/cause behavioral changes in crawling velocity.

The density of the cross-covariance values near *τ* = 0 in PVM indicates a tight temporal coupling between sensory neuron activity and ongoing locomotion. Whereas control animals show a more pronounced and consistent covariance peak at negative and positive lag, *mec-4* mutants generally exhibit a broader and less coherent distribution, suggesting reduced or more variable coupling when mechanotransduction is impaired.

Whereas the function of TRNs is well studied and they mediate escape reflex to mechanical stimuli, the function of the PVM neuron is currently not known. We investigated the hypothesis that it encodes aspects of the animal’s movement (responding to velocity). The covariance structure therefore reflects a combination of sensory encoding (velocity → Ca; *τ >*0) and motor command generation (Ca → velocity change; *τ <* 0).

In control animals, the sharp covariance structure suggests:

1. Reliable sensory encoding: changes in velocity are consistently followed by changes in calcium at short positive lags.
2. Effective motor output: increases in Ca often precede behavioral change in locomotion at short negative lags.

In *mec-4* mutants, the flat covariance patterns indicate an impaired mechanosensory drive: the neuron’s calcium signal tracks velocity less reliably. It also suggests a weaker motor influence: Ca events do not translate into consistent reversal initiation or velocity changes.

The overall reduction in covariance amplitude and temporal precision in *mec-4* animals is consistent with a loss of mechanosensory input through the MEC-4 channel, affecting both how movement is encoded and how sensory activation is converted into behavioral output.

As displayed in the Supplementary Figure 4a below, the individual quadrants can be explained by the following:

1. Q1: The positive lag of the positive cross-covariance indicates that movement triggers a change in calcium activity. This is consistent with the hypothesis outlined here that PVM is activated by velocity-dependent substrate sensing (e.g. traction-dependent activity).
2. Q2: The negative lag of the positive covariance indicates that calcium activity triggers forward movement. This is consistent with the idea that PVM causes accelerations similar to other posterior touch receptor neurons such as PLM.
3. Q3: The negative lag of the negative covariance indicates that increase in calcium activity in PVM triggers decrease in velocity. This is neither inconsistent with the idea of the standard touch response nor with our hypothesis, and indeed this is the quadrant with the least observations in our data.
4. The positive lag of the negative covariance indicates that the decrease in locomotion velocity triggers a PVM calcium transient. This is consistent with the hypothesis that PVM is activated by velocity-dependent substrate sensing (e.g. traction-dependent activity). The notion that PVM is activated by forward and backward locomotion points towards a direction-invariant traction sensing mechanism. Further studies need to be conducted to investigate if traction is larger during forward or backward crawling.

We have highlighted how these quadrants map onto the raw calcium and tangential velocity traces in the Supplementary Figure 4b.

Because of the high covariance in quadrant 1, it gives first direct hints that PVM has a role on substrate and velocity sensation as studied in detail in this paper. The high covariance in Q2 also suggests that PVM may drive acceleration and reversals. Together, we put the idea forward that PVM has a dual context dependent role as a substrate mechanosensor to adjust proprioception and motor output.

## 4 Supplementary Figures

**Supplementary Fig. 1.**
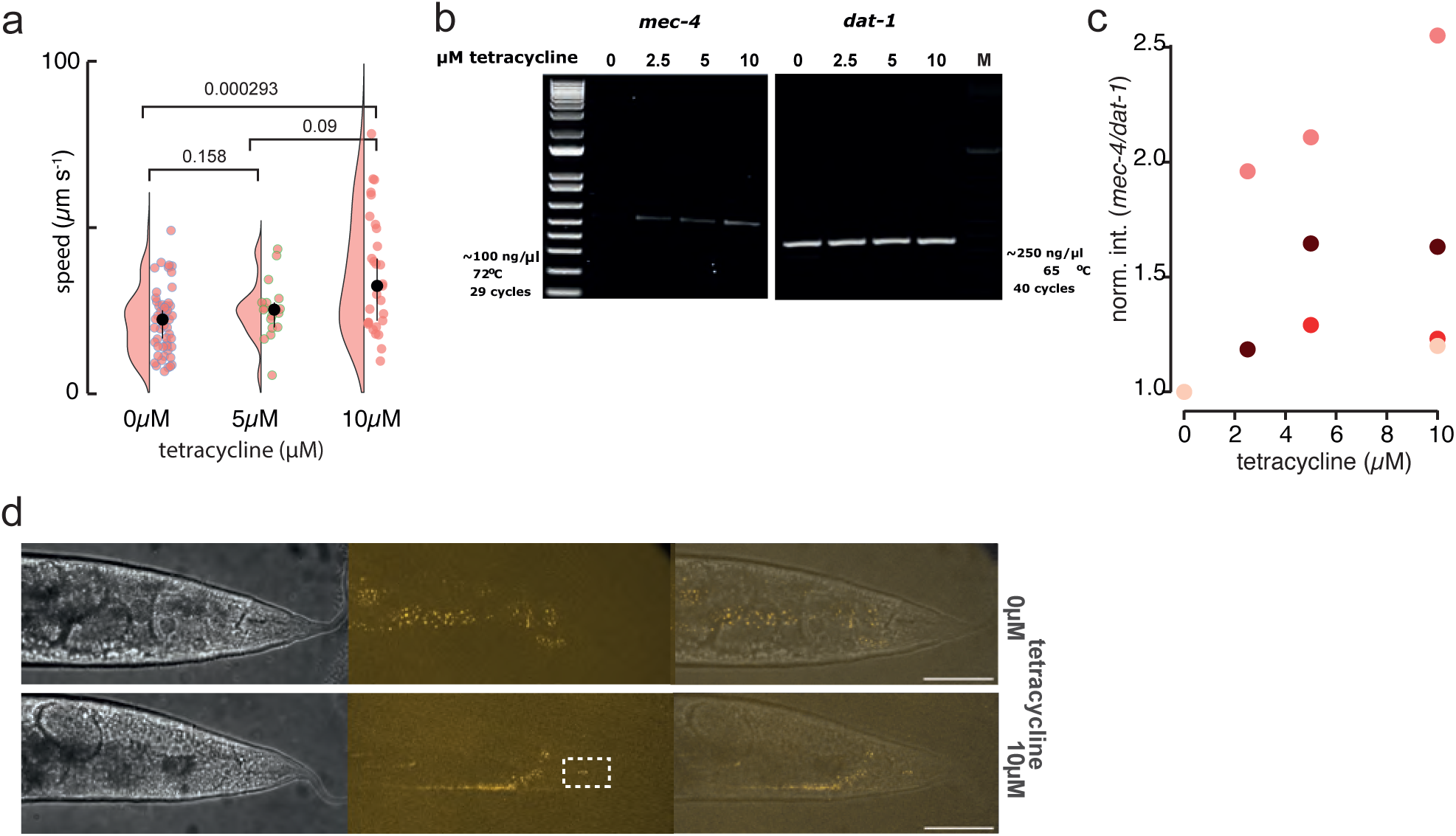
Temporal *mec-4* regulation through a tetracycline-dependent ribozyme. **a,** Average crawling speed of wild-type animals exposed to varying concentrations of tetracycline. p-values above the bracket show the result of a two-tailed Kolmogoroff-Smirnoff test for a pairwise comparison of the indicates conditions. **b,** Representative gel image of tetracyclin-dependent *mec-4* expression and *dat-1* as a tet-independent control. **c,** Quantification of *mec-4* gene expression. Independent experiments are represented as a different color. N = 4. Each data point was normalized to untreated control. **d,** Brightfield and fluorescent micrographs of an animal with and without tetracycline addition. The PLM cell body is indicated in the dotted box. Image is representative for N=10 animals that have been examined. Scale bar = 40 µm.

**Supplementary Fig. 2.**
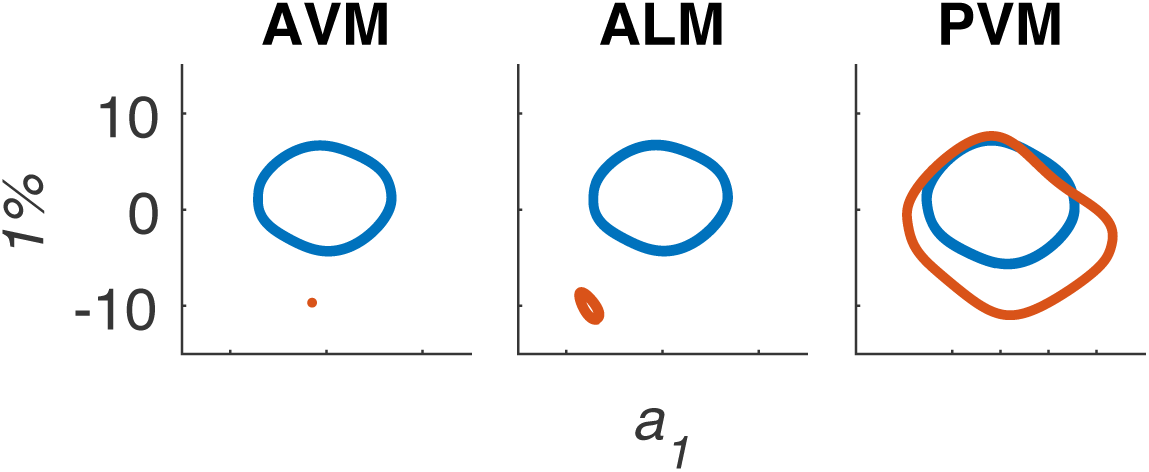
Simulation of the motor output in the *mec-4* mutant. Representation of the combined motor neuron activity in the low dimensional subspace after driving forward locomotion through current injection into PLM (see also ref. (*29*)). We simulated TRN sensory capacity by injecting a baseline current of 1% of the driving current into AVM, ALM and PVM (blue limit cycle). Disruption of the basal current (e.g. simulating *mec-4* mutant condition) leads to a loss of motor neuron activity, visible as fixed points in the subspace of AVM and ALM, and the described increase in the limit cycle in the PVM subspace.

**Supplementary Fig. 3.**
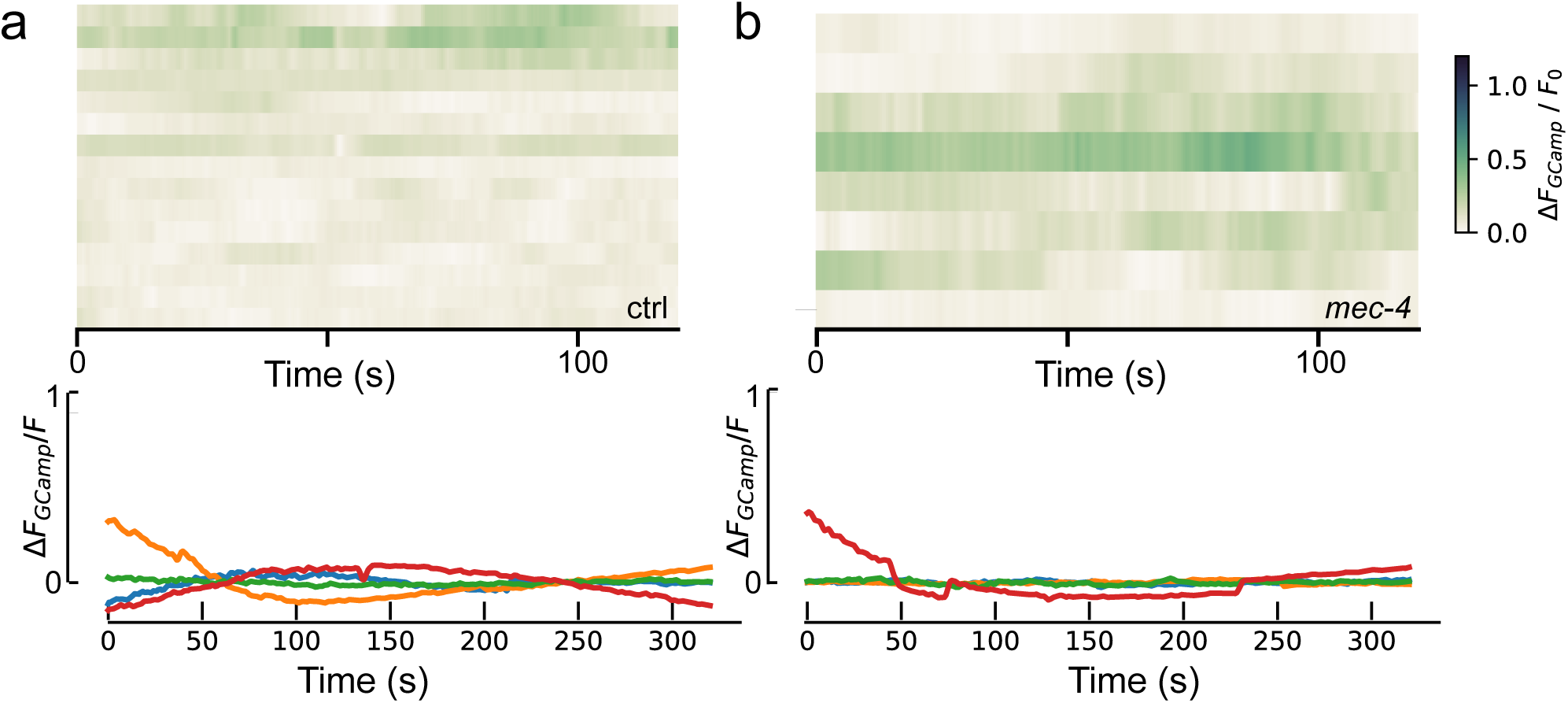
TRNs do not mobilise calcium in immobilized animals. **a, b,** Kymograph (i) and selected traces (i) of the calcium activity measured in PLM cell body of immobilized (a) wild-type, and (b) *mec-4* mutant animals, as opposed to spontaneous, locally confined and non-propagating calcium transients observed in TRN neurites (Extended Data Fig. 2.)

**Supplementary Fig. 4.**
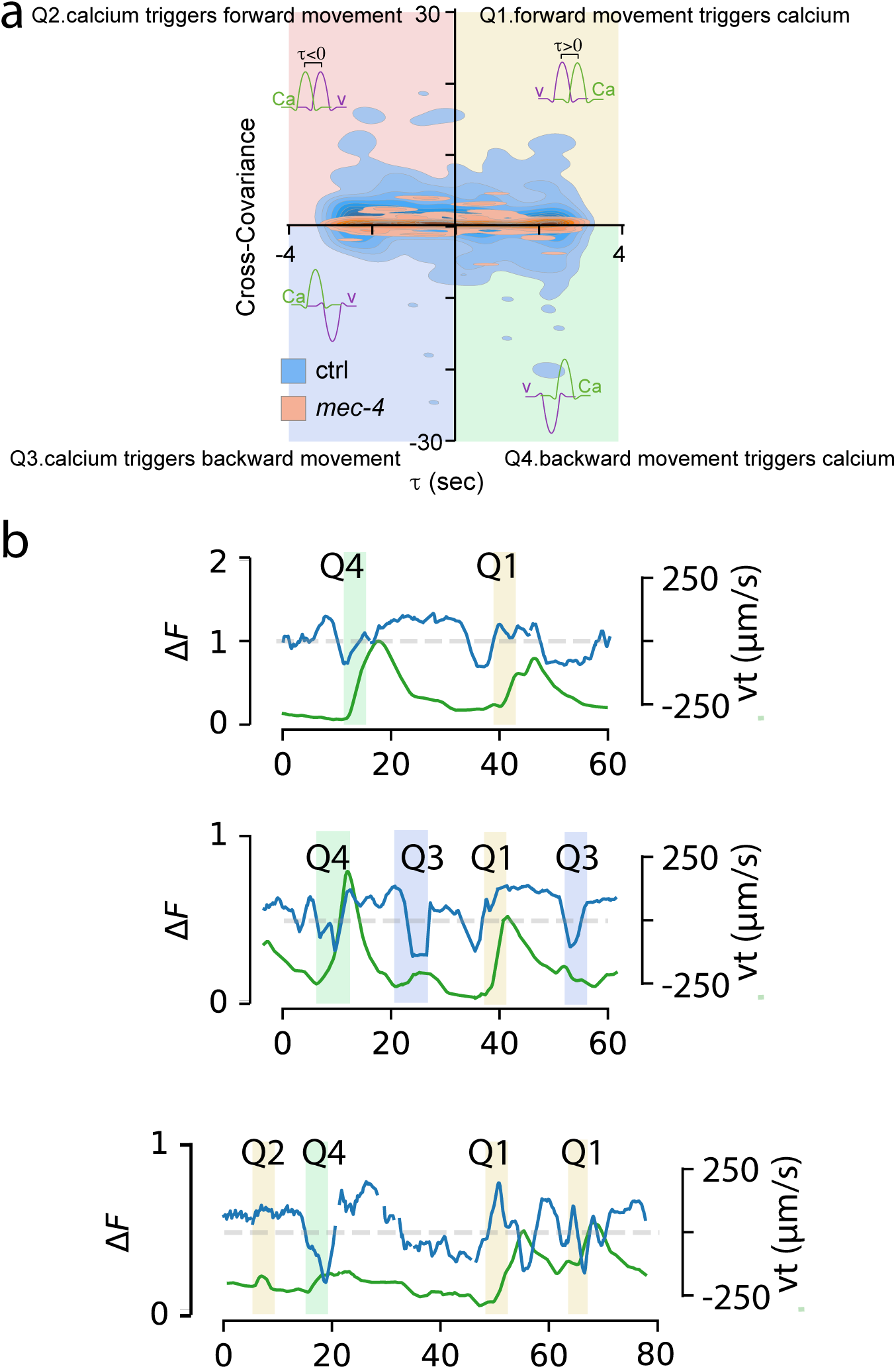
Individual events and how they contribute to the Calcium - velocity cross-covariance. **a,** Detailed description and case examples for the cross-covariance computed between PVM:GCaMP intensity and animals crawling velocity. **b,** Highlighted events that map into the cross covariance.

## 5 Supplementary Tables

**Table 1.** Plasmids used in this study.

| plasmid name | promoter | ORF | 3'UTR |
| --- | --- | --- | --- |
| pNM3 | *mec-4* | cGal4 | *let-858* |
| pNS50 | *mec-4* | mScarlet1-10 | *unc-10* |
| pNMSB184 | *flp-8* | N-CRE | *tbb-2* |
| pNMSB185 | *twk-49* | C-CRE | *tbb-2* |
| pNMSB186 | *mec-4* | mScarlet::SL2::TeTx | *let-858* |
| pNS121 | *mec-18* | *mec-4*(cDNA)::SL2::mCherry | *unc-10* |

**Table 2.**
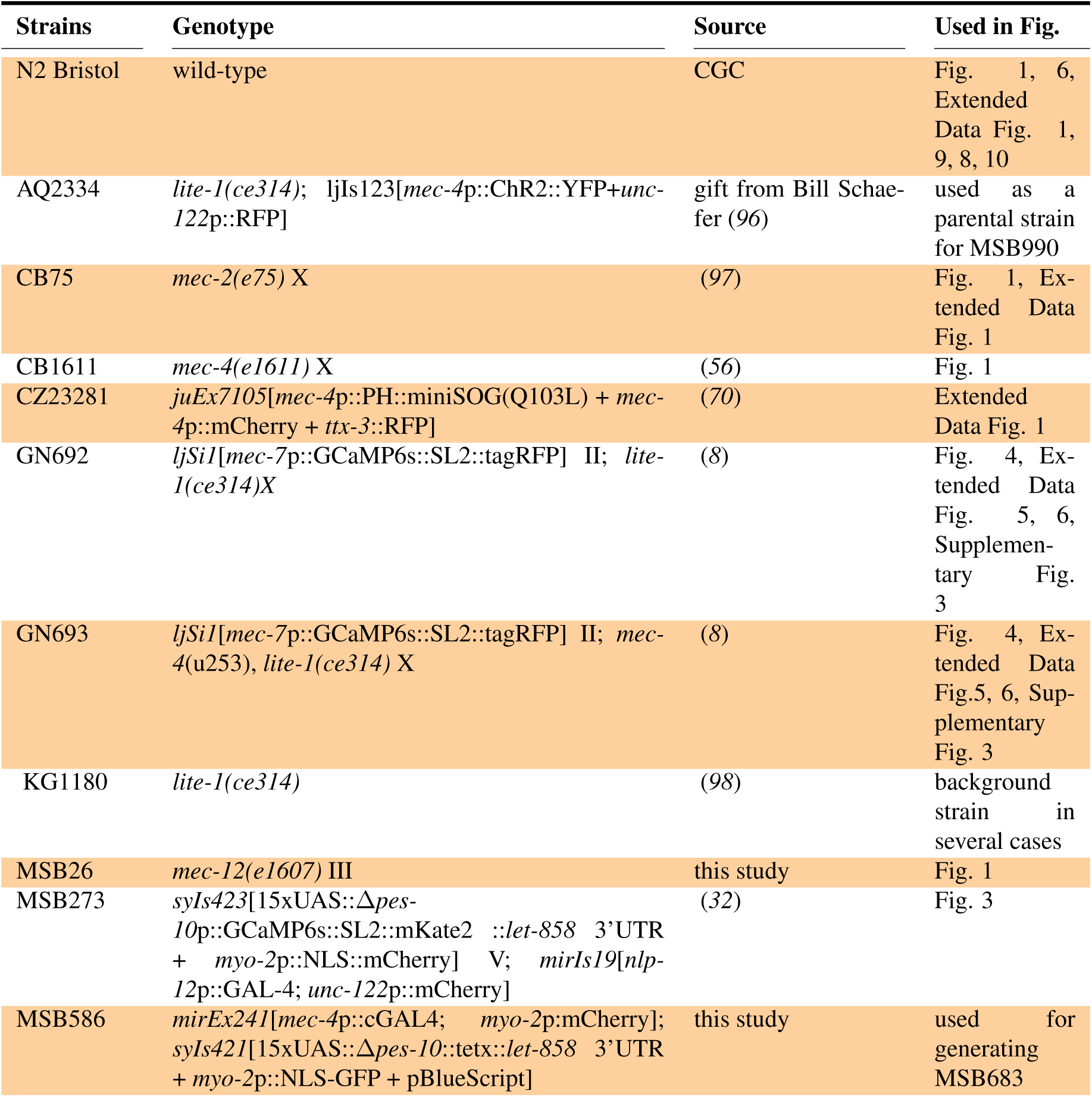
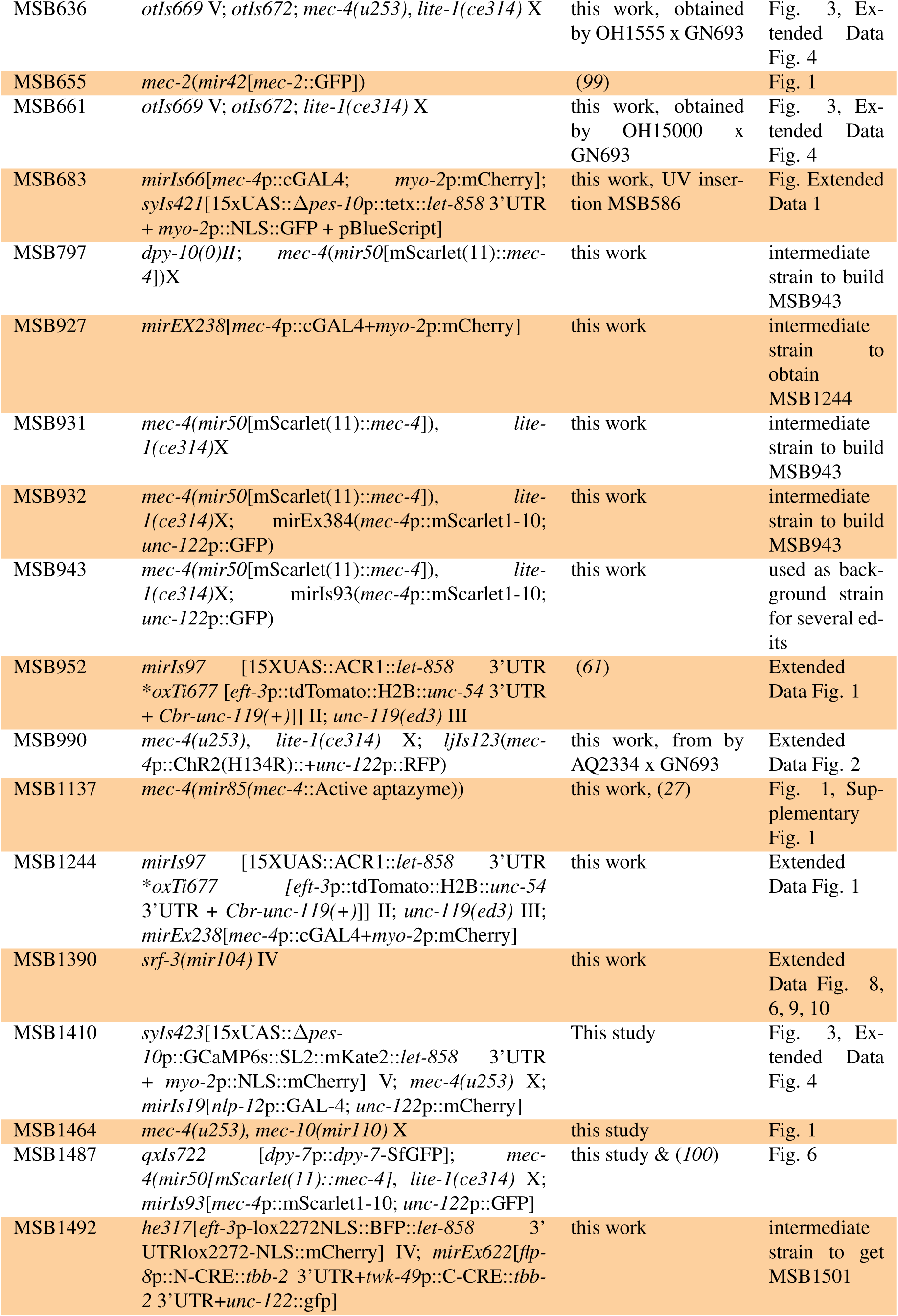
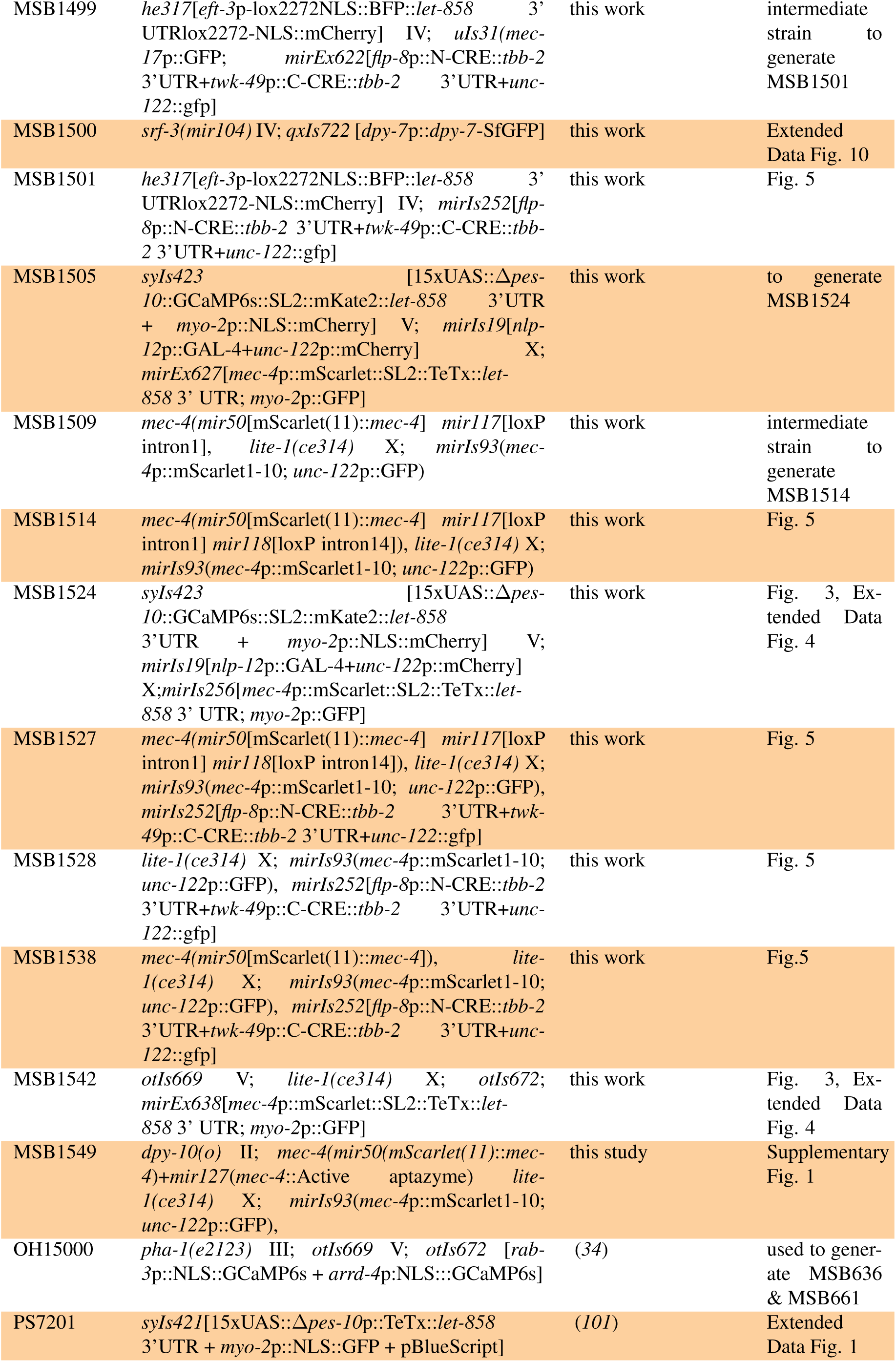
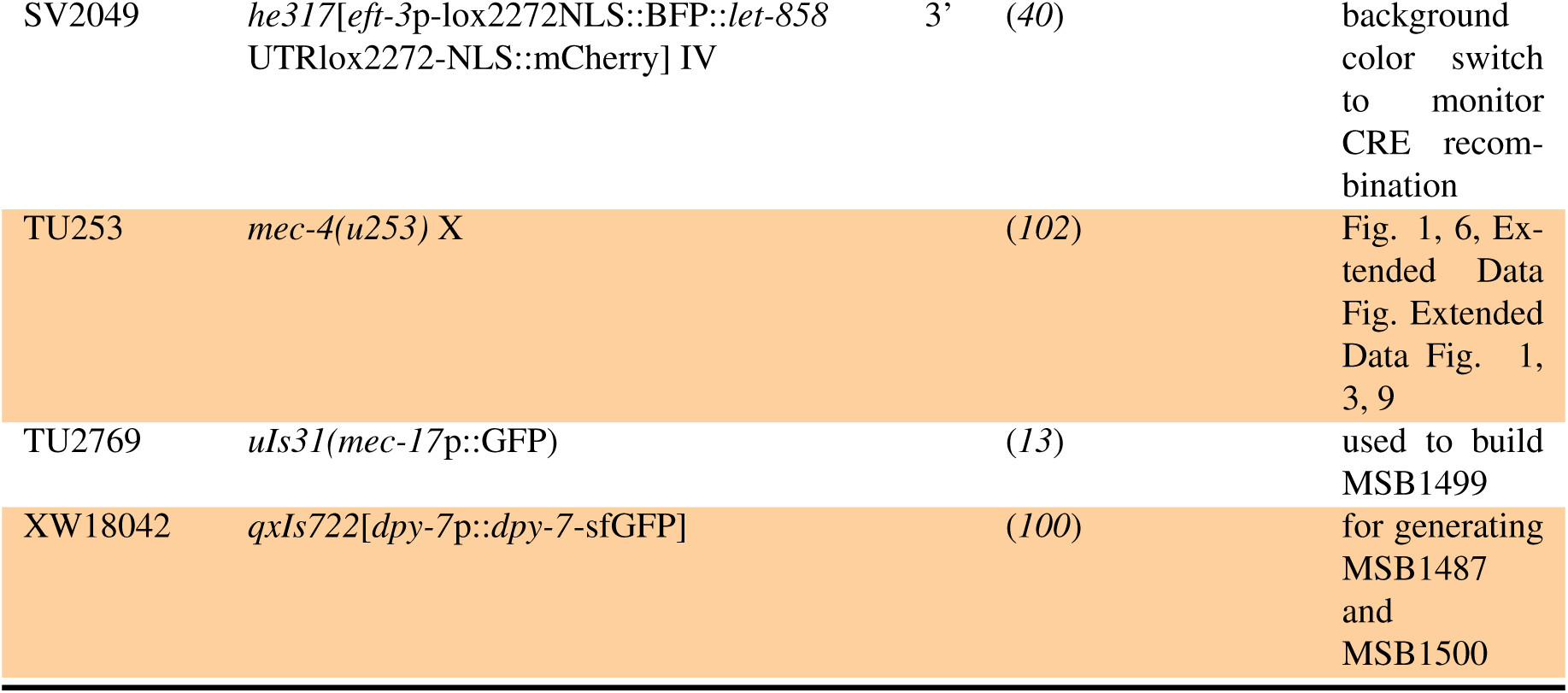
Strains used in this study.

**Table 3.**
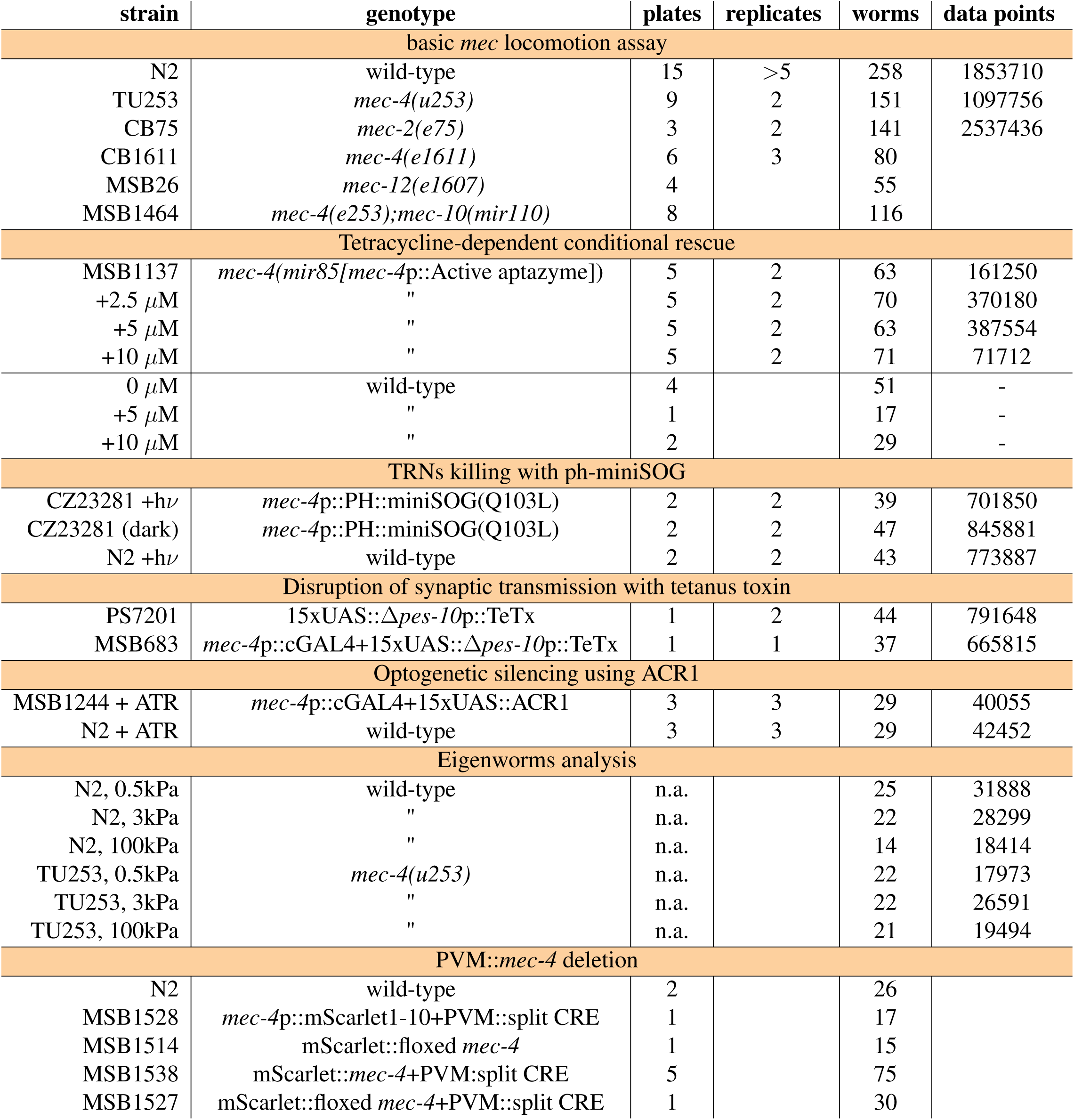
Worm locomotion data: Number of plates, days, animals and frame analyzed from the locomotion experiments.

| strain | genotype | plates | replicates | worms | data points |
| --- | --- | --- | --- | --- | --- |
| basic *mec* locomotion assay |  |  |  |  |  |
| N2 | wild-type | 15 | >5 | 258 | 1853710 |
| TU253 | *mec-4(u253)* | 9 | 2 | 151 | 1097756 |
| CB75 | *mec-2(e75)* | 3 | 2 | 141 | 2537436 |
| CB1611 | *mec-4(e1611)* | 6 | 3 | 80 |  |
| MSB26 | *mec-12(e1607)* | 4 |  | 55 |  |
| MSB1464 | *mec-4(e253);mec-10(mir110)* | 8 |  | 116 |  |
| Tetracycline-dependent conditional rescue |  |  |  |  |  |
| MSB1137 | *mec-4(mir85[mec-4p::Active aptazyme])* | 5 | 2 | 63 | 161250 |
| +2.5 μM | " | 5 | 2 | 70 | 370180 |
| +5 μM | " | 5 | 2 | 63 | 387554 |
| +10 μM | " | 5 | 2 | 71 | 71712 |
| 0 μM | wild-type | 4 | | 51 | - |
| +5 μM | " | 1 | | 17 | - |
| +10 μM | " | 2 | | 29 | - |
| TRNs killing with ph-miniSOG |  |  |  |  |  |
| CZ23281 +hν | *mec-4p::PH::miniSOG(Q103L)* | 2 | 2 | 39 | 701850 |
| CZ23281 (dark) | *mec-4p::PH::miniSOG(Q103L)* | 2 | 2 | 47 | 845881 |
| N2 +hν | wild-type | 2 | 2 | 43 | 773887 |
| Disruption of synaptic transmission with tetanus toxin |  |  |  |  |  |
| PS7201 | 15xUAS::*$\Delta pes-10p::TeTx$* | 1 | 2 | 44 | 791648 |
| MSB683 | *mec-4p::cGAL4+15xUAS::$\Delta pes-10p::TeTx$* | 1 | 1 | 37 | 665815 |
| Optogenetic silencing using ACR1 |  |  |  |  |  |
| MSB1244 + ATR | *mec-4p::cGAL4+15xUAS::ACR1* | 3 | 3 | 29 | 40055 |
| N2 + ATR | wild-type | 3 | 3 | 29 | 42452 |
| Eigenworms analysis |  |  |  |  |  |
| N2, 0.5kPa | wild-type | n.a. |  | 25 | 31888 |
| N2, 3kPa | " | n.a. |  | 22 | 28299 |
| N2, 100kPa | " | n.a. |  | 14 | 18414 |
| TU253, 0.5kPa | *mec-4(u253)* | n.a. |  | 22 | 17973 |
| TU253, 3kPa | " | n.a. |  | 22 | 26591 |
| TU253, 100kPa | " | n.a. |  | 21 | 19494 |
| PVM::*mec-4* deletion |  |  |  |  |  |
| N2 | wild-type | 2 |  | 26 |  |
| MSB1528 | *mec-4p::mScarlet1-10+PVM::split CRE* | 1 |  | 17 |  |
| MSB1514 | mScarlet::*floxed mec-4* | 1 |  | 15 |  |
| MSB1538 | mScarlet::*mec-4*+PVM:split CRE | 5 |  | 75 |  |
| MSB1527 | mScarlet::*floxed mec-4*+PVM::split CRE | 1 |  | 30 |  |

**Table 4.** Number of worms/points used covariance and cross-covariance plots. Overall 163 neurons from 158 different worms, 7 different experimental days, each recording lasted 2 min.

| Ca<sup>2+</sup> dynamics in the TRNs freely moving worms |  |  |  |  |
| --- | --- | --- | --- | --- |
| strain | neuron | 1.25 kPa | 6 kPa | 30 kPa |
| GN692 | PLM | 13/162 | 8/92 | 13/187 |
| GN693 |  | 17/171 | 17/114 | 14/171 |
| GN692 | PVM | 8/65 | 7/39 | 7/118 |
| GN693 |  | 11/48 | 7/15 | 10/103 |
| GN692 | ALM | 9/81 | 6/28 | 13/133 |
| GN693 |  | 12/73 | 9/64 | 11/62 |
| GN692 | AVM | 4/18 | 5/13 | 8/45 |
| GN693 |  | 4/8 | 5/39 | 7/27 |

**Table 5.** Number of traction force experiments performed in the different conditions.

| Traction Force Microscopy |  |  |  |
| --- | --- | --- | --- |
| strain | 1.25 kPa | 6 kPa | 30 kPa |
| N2 | 8 | 8 | 6 |
| TU253 | 9 | 6 | 8 |
| MSB1390 | 9 | 7 | 7 |

## 6 Supplementary Videos

**6.1 Supplementary Video 1**

Representative video of N2 control, *mec-4* and *mec-2* mutant animals on NGM agar supplemented with food.

**6.2 Supplementary Video 2**

Representative video of the crawling N2 control animal on 0.5kPA PAA hydrogel.

**6.3 Supplementary Video 3**

Representative video of the *mec-4* mutant animal crawling on 0.5kPA PAA hydrogel.

**6.4 Supplementary Video 4**

Representative video of control animal expressing GCaMP6s in PDE, PVM, PVD and SDQ, subjected to three consecutive mechanical stimuli anterior to the PVM cell body (left most stimulator). Posterior to the right.

**6.5 Supplementary Video 5**

Representative video of the control animal expressing GCaMP6 in DVA, subjected to three consecutive mechanical stimuli to the PVM neurites.

**6.6 Supplementary Video 6**

Representative calcium recording of control animals expressing GCaMP6s in TRNs crawling on 1.25 kPA PAA hydrogel.

**6.7 Supplementary Video 7**

Representative calcium recording of *mec-4* mutant animals expressing GCaMP6s in TRNs crawling on 1.25 kPA PAA hydrogel.

**6.8 Supplementary Video 8**

Representative recording of a traction force microscopy experiment using wild-type control animal crawling on 6kPA PAA hydrogel. Displacement is visualized using gel-embedded microspheres.

**6.9 Supplementary Video 9**

Representative result of the model used to infer the friction coefficients, showing the locomotion of the worm on 6kPa substrate and the underlying traction components. The static dashed line is the initial body posture, thick line is the worm’s backbone with inferred non-homogeneous friction and the thin blue line is the hypothetical worm’s backbone with homogeneous friction coefficient.

## Notes

### Competing Interest Statement

The authors have declared no competing interest.

